# Template-Independent Enzymatic RNA Synthesis

**DOI:** 10.1101/2024.10.09.617423

**Authors:** Nilesh B Karalkar, Tatiana Kent, Taylor Tredinnick, Leonardo Betancurt-Anzola, Marc Delarue, Richard Pomerantz, Steven A Benner

## Abstract

A route to prepare ribonucleoside triphosphates featuring a 3’-aminoxy (3’-O-NH_2_) removable blocking group is reported here. We then show that versions of two DNA polymerases, human DNA polymerase theta (Polθ) and mimiviral PrimPol, accept these triphosphates as substrates to add single nucleotides to an RNA primer under engineered conditions. Cleaving the O-N bond in the 3’-O-NH_2_ group within the extended primer regenerates the 3’-OH group, facilitating subsequent polymerase cycles that add a second, selected, nucleotide. These enzymes and triphosphates together enable template-independent enzymatic RNA synthesis (TIERS) exploiting a cyclic reversible termination framework. The study shows that this process is ready for instrument adaptation by using it to add three ribonucleotides in three cycles using an engineered Polθ. This work creates a new way to synthesize RNA with a de novo defined sequence, without requiring the protecting groups, hazardous solvents, and sensitive reagents that bedevil phosphoramidite-based RNA synthesis.

## INTRODUCTION

RNA molecules have long been “chemically” synthesized using the same phosphoramidite reagents that are today used for DNA synthesis.^1^ This is especially true for short RNA molecules,^2^ which are used for a spectrum of biochemical and biological applications.^3^ Thus, direct synthesis supplements a synthetic process that makes RNA by transcribing synthetic DNA by an RNA polymerase. This process is used to make longer RNA molecules, including vaccines.^4^

Phosphoramidite-based synthesis has become so routine that some synthetic biologists have come to believe that “DNA synthesis should cost next to nothing”^5^. In contrast, phosphoramidite-based synthesis of RNA still presents many challenges that are not presented by DNA synthesis. These arise from the additional 2’-OH groups in the RNA building blocks, which require special protection. Further, RNA chains are susceptible to cleavage under the alkaline conditions typically used in DNA synthesis to release synthetic products from solid supports and to deprotect nucleobases. Despite considerable work in this area, protecting groups, deprotection conditions, and workflows for phosphoramidite-based synthesis of oligomeric RNA are still expensive and limited. Thus, improving methods for RNA synthesis is an important goal in biotechnology.

Enzymatic reactions, of course, do not require protecting groups^6^ to manage reactivity in multi-functional substrates. Instead, enzymes achieve specificity by binding substrates in active sites with controlled geometry. This geometry guides catalytic functionality to only the desired locations. Enzymatic solvents are simply buffered water. The byproducts of enzymatic reactions are generally innocuous. This mitigates the cost and the environmental impact with enzyme-dependent process.^7^

Accordingly, many efforts are today directed towards developing enzyme-dependent processes to make RNA without a previously synthesized DNA template.^8^ Conceptually, this involves a stepwise process that might begin when a selected ribonucleoside triphosphate is presented to an enzyme that can use these as substrates to elongate an RNA or single-strand DNA (ssDNA) primer. After this first ribonucleotide is added, subsequent ribonucleotides might be added in a step-wise fashion. The specific ribonucleoside triphosphate introduced for each addition step would determine the sequence of the resulting RNA.

Certain polymerase enzymes, such as poly-A polymerase and poly-U polymerase, catalyze such untemplated primer extension reactions with RNA substrates and their respective NTPs.^9^ However, if standard ribonucleoside triphosphates with a free 3’-OH group are used as substrates, nucleotide extension does not stop after a single nucleotide is added. Rather, primer extension continues to form a homopolymer.

Returning to an analogy to enzyme-assisted DNA synthesis, some time ago, we noted that the smallest removable 3’-O blocking group consistent with the Periodic Table and rules of chemical binding was 3’-O-NH_2_, the “aminoxy” group.^10^ When presented on an incoming triphosphate, the aminoxy group blocks further primer extension. However, the blocking group can be removed under mild conditions (for example, sodium nitrite buffered at pH 6).

The aminoxy group was developed to be used in DNA sequencing that exploits a cyclic reversible termination architecture.^11^ However, it was then adopted for enzyme-dependent DNA synthesis,^12^ most notably by DNA Script, which now sells an instrument (*Syntax*^TM^)^13^ that uses the aminoxy group as a reversible 3’-blocking group for enzyme-dependent DNA synthesis.^14^ The workflow in *Syntax*^TM^ avoids the complex protection, hazardous solvents, and toxic reagents in phosphoramidite-based DNA synthesis.

Motivated by the success of 3’-aminoxy groups as removable terminators in template-independent enzymatic *DNA* synthesis, we considered using the 3’-aminoxy group as a reversibly terminating blocking group in template-independent enzymatic RNA synthesis (TIERS).

Unfortunately, the additional complexity of RNA building blocks over DNA building blocks proved to make the synthesis of 3’-O amino ribonucleoside triphosphates also more challenging than for the corresponding 3’-O amino *2’-deoxy*ribonucleoside triphosphates.

In this study, we report the successful synthesis of ribonucleoside triphosphates featuring a 3’-O aminoxy blocking group. This enabled production of all four ribonucleoside triphosphates with 3’-ONH_2_ aminoxy blocking groups. Subsequently, these synthetic compounds were used as substrates to evaluate a set of primase-polymerase (PrimPol) and other polymerase enzymes able to work in an untemplated fashion, investigating their compatibility in controlled primer extension reaction cycles.

The synthesis of 3’-ONH_2_-NTPs allowed us to explore their suitability as substrates for these polymerases. Additionally, we developed methods that allow for stepwise ribonucleotide additions using 3’-ONH_2_ aminoxy blocking groups and an engineered DNA polymerase θ (Polθ)· Together, these findings lay the foundation for a platform using enzymes to enhance the efficiency of RNA synthesis, setting the stage for its application with automation techniques.

## RESULTS AND DISCUSSION

Architectures for enzyme-dependent RNA synthesis generally require that the 3’-OH of the ribonucleoside triphosphate substrates be reversibly blocked by a moiety that prevents further addition after the first ribonucleotide is added. Many have turned to the 3’-O removable blocking groups that were developed for cyclic reversible termination architectures used in DNA sequencing. These include the azidomethyl,^15^ methoxymethyl,^16^ and allyl groups (**Fig. 1**).^17^

**Figure 1.**
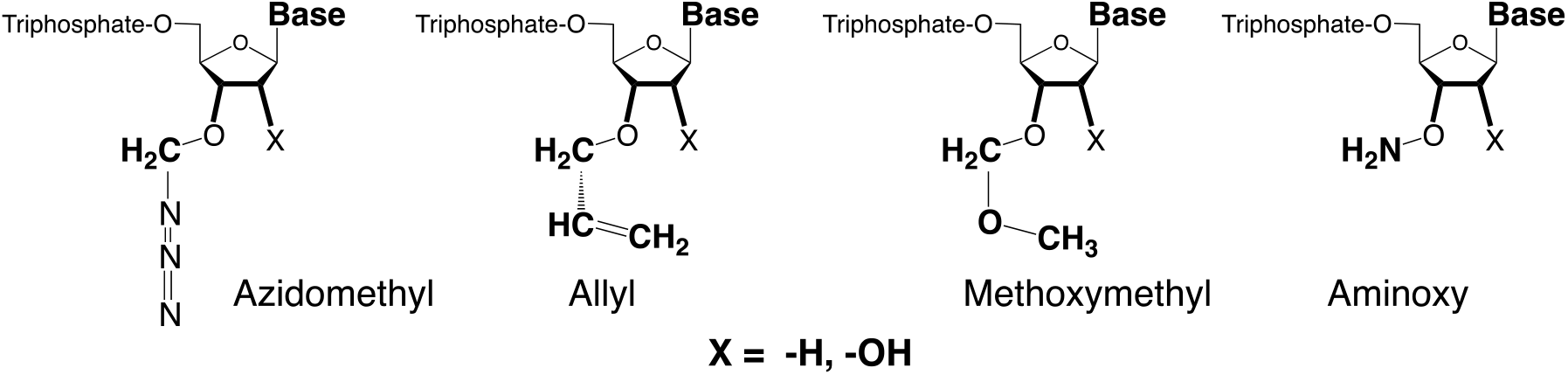
Four removable blocking groups might be recruited from next generation sequencing to support enzyme-dependent RNA synthesis. EnPlusOne is working with the 3’-O-allyl group.^18^ Work reported here exploits the aminoxy group (far right). Since the aminoxy group is smaller, more hydrophilic, and recapitulates the hydrogen bonding potential of the native –OH group, engineering enzymes to accept the aminoxy group is less difficult than with the other removable blocking groups.

Unfortunately, these groups are substantially larger than the hydrogen atom that they replace. This means that they do not fit easily into natural polymerase active sites. Further, polymerases have evolved for billions of years to closely inspect the 2’, 3’ region of their triphosphate substrates, so as to discriminate deoxyribonucleoside triphosphates (dNTPs) from ribonucleoside triphosphates (NTPs), both present together in cellular environments. These close inspection mechanisms for 2’ and 3’, selected over billions of years of evolution, make it likely that polymerases will discriminate against nucleotides with *any* 3’-O blocking modifications. Indeed, getting polymerases to accept 2’- and 3’-modified triphosphates has often involved protein engineering.^19^

### Synthesis of Reversibly Terminating Nucleotide Building Blocks

Our first efforts to synthesize ribonucleoside triphosphates with 3’-ONH_2_ blocking groups sought to follow routes that we had used previously to make 3’-ONH_2_ 2’-deoxyribonucleoside triphosphates from the 2’-deoxyribonuceosides. Here, we sought to apply two Mitsunobu-type reactions to ribonucleosides that had their 2’- and 5’-hydroxyl groups protected (**Fig. 2**):

**Figure 2.**
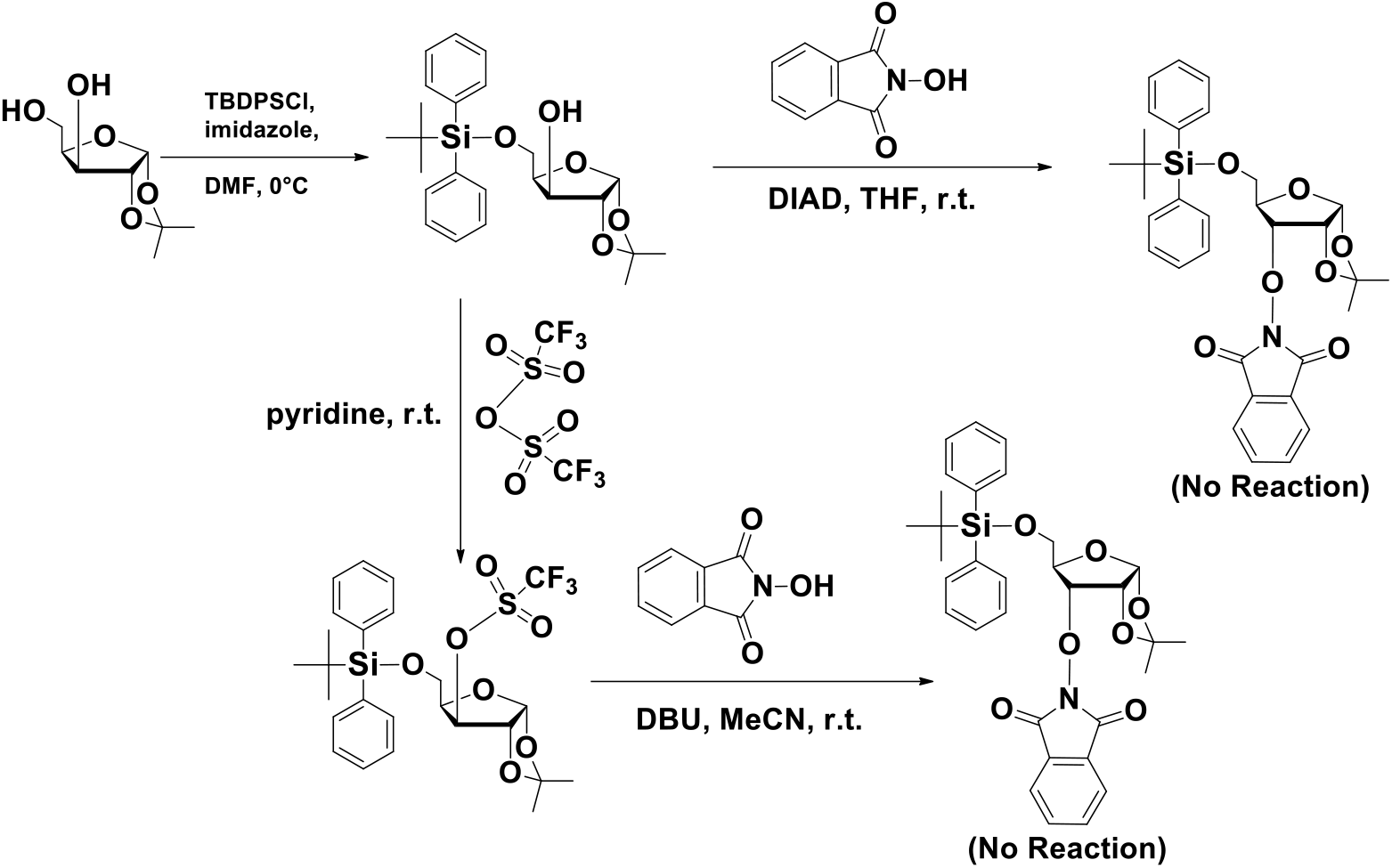
Failed routes to make ribonucleosides with a 3’-ONH_2_ moiety. (**Top**) Attempted inversion by a Mitsunobu reaction with N-hydroxyphthalimide as a nucleophile. (**Below**) Attempted S_N_2 displacement of a trifluoromethylsulfonate using N-hydroxyphthalimide as a nucleophile.

- In the first Mitsunobu reaction, a benzoate nucleophile was used to give a beta-benzoate with inversion of stereochemistry at the 3’-position.
- After hydrolysis, in the second Mitsunobu reaction, N-hydroxyphthalimide was used as the nucleophile to give a second stereochemical inversion to yield the 3’-O aminoxy compound protected as its phthalimide, in the alpha configuration.

Unfortunately, this double Mitsunobu reaction sequence proved ineffective when applied to ribonucleosides.

In search of an alternative approach, we attempted a direct S_N_2 displacement of the 3-OH of a protected xylose derivative (**Fig. 2**). Here, the 3-hydroxyl group was derivatized as a trifluoromethanesulfonate. Then, we attempted to displace the sulfonate with stereochemical inversion using the N-hydroxyphthalimide nucleophile. Unfortunately, this also failed to yield the desired phthalimido derivative. These repeated failures align with a model that takes into account the extent to which the hydroxyl group, which we aim to react with, is sterically hindered within the ribose molecule.^20^ Notably, this hindrance is less pronounced in the 2’-deoxyribose analogs.

Therefore, we developed a novel oxidation-reduction approach starting with the inexpensive xylose acetonide (**Fig. 3**). Its primary hydroxyl group was protected as a tertiary-butyl-diphenylsilyl ether. This allowed clean oxidation of the secondary hydroxyl group to form a ketone, which was then reduced with inversion of stereochemistry to yield an intermediate with the ribose stereochemistry. The amino group was directly introduced onto the free hydroxyl group using an electrophilic sulfonate aminating reagent. The resulting aminoxy group was selectively protected as its oxime with acetone, the acetonide group was removed under acidic conditions, and the free hydroxyl groups were converted to acetate esters.

**Figure 3.**
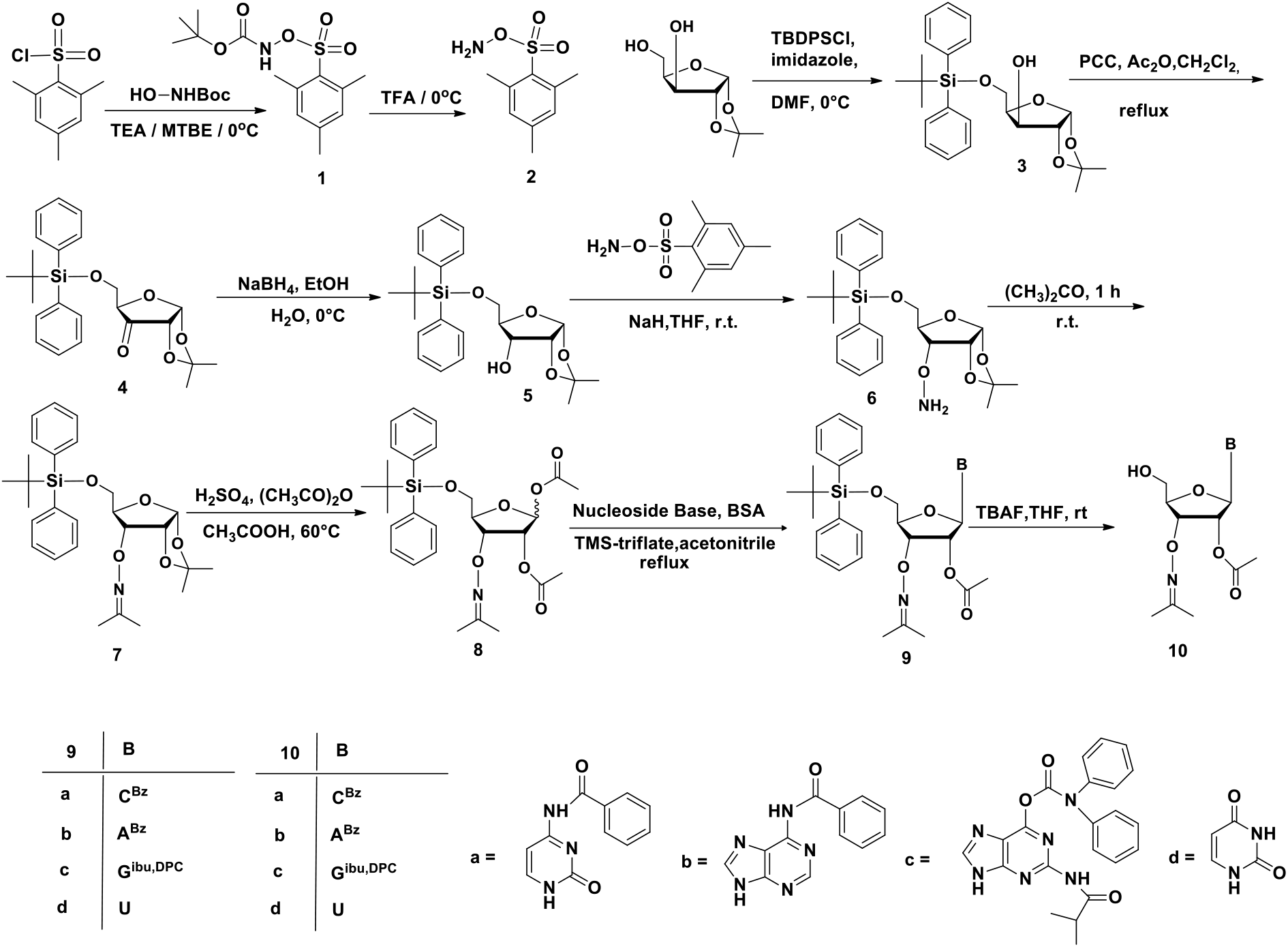
Synthesis of a common intermediate for all four standard nucleosides with 3’-ONH_2_ aminoxy removable blocking groups.

This sequence created multi-gram quantities of a common intermediate for all four standard ribonucleotides. This intermediate was directly combined with the nucleobases to form the 3’-O-NH_2_ nucleotides. The correct stereochemical orientation of the base (beta) assured by the participation of the 2’-acetoxy group. The 3’-O-NH_2_ nucleosides were then converted into their respective triphosphates by using the Ludwig Eckstein reaction (**Fig. 4**).^21^

**Figure 4.**
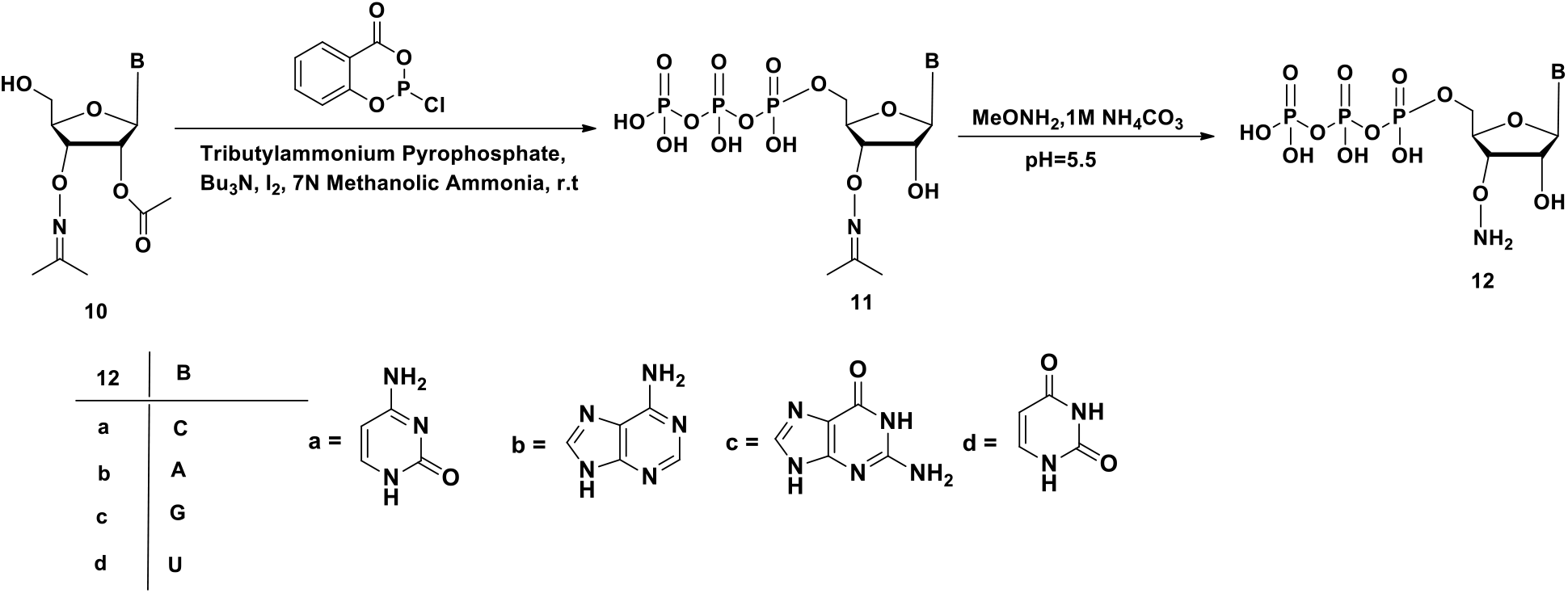
Synthesis of triphosphates of four standard nucleosides with 3’-ONH_2_ aminoxy groups.

### Polymerase Exploration

The 3’-O-NH_2_ moiety is the smallest removable blocking group possible on the Periodic Table in molecules that follow normal rules of bonding. Because of this, and because it recapitulates the hydrophilicity and hydrogen bonding potential of the 3’-OH group, the aminoxy moiety is the removable blocking group most likely to fit into the active sites of enzymes that accept standard nucleoside triphosphates as substrates.

Balancing this, natural enzymes must have evolved to distinguish ribo-from deoxyribo-nucleoside triphosphates, which occur together in their natural environments. This means that any enzyme that we might recruit to use 3’-O-NH_2_ nucleoside triphosphates has evolved to closely inspect the 2’,3’ region of their substrates, making it less likely that they will tolerate *any* substituent in this region.

This drove us to examine polymerase families^22,23^ that might naturally make this inspection less rigorously, in particular, polymerases that have evolved to tolerate bulky or non-canonical nucleotides such as translesion DNA polymerases^24^ and/or that can work in an untemplated fashion.^29,33,36^ Translesion polymerases have evolved to incorporate non-canonical nucleotides and promote replication opposite non-canonical damaged template nucleotides. Not all translesion polymerases, however, are likely to be active on single-strand RNA or efficiently incorporate ribonucleotides. Thus, we examined two polymerases that have been successfully engineered in prior studies and are known to accommodate multiple nucleic acid templates and exhibit untemplated synthesis activity.

The first was PrimPol from mimivirus,^25,26^ a polymerase displaying also primase activity. PrimPol is a member of the primase-polymerase superfamily whose members participate in many processes.^27^ These include the unprimed synthesis of RNA primers that are then extended by canonical DNA and restart DNA synthesis (or repriming) at damaged DNA replication forks in nuclei and mitochondria. Further, mutant forms of mimivirus PrimPol have been designed to alter substrate specificity without damaging catalytic activity.

A crystal structure of mimivirus PrimPol specifically has not been solved yet. However, some of its homologs are known. Thus, in previous work, a model for mimivirus PrimPol was built using AlphaFold2.^28,29^ The predicted structure was then overlapped with a solved structure of human PrimPol that has a docked double strand DNA molecule and an incoming nucleoside triphosphate (RMSD across 207 C-alpha pairs: 2.7 Å). Amino acid side chains in the vicinity of the 2’/3’ atoms of that triphosphate were listed by their geometric proximity in the model. Those that were highly conserved in homologous proteins were excluded from the list, and amino acids present in less highly conserved sites were replaced by amino acids with smaller side chains.

This provided a manifold of mimivirus PrimPol variants for testing, called: pLB001 (Wild type), pLB002 (L223V), pLB003 (L223T), pLB004 (L223S), pLB014 (L223G), pLB016 (L223F), pLB005 (L223A), pLB015 (L223P).

The second enzyme inspected was DNA polymerase theta (Polθ; gene name *POLQ*). Polθ is a multi-domain protein with an A-family DNA polymerase at its C-terminus^30^ closely related to Klenow fragment and *Thermus aquaticus* DNA polymerase^31, 32^.

Polθ performs translesion synthesis opposite various damaged DNA nucleotides and is able to accommodate non-canonical large deoxyribonucleotides.^33,34^ Polθ is also able to accommodate A-form DNA/RNA primer-templates, facilitated by a significant conformational change in the thumb subdomain that allows for active binding to broader DNA/RNA hybrids^35^. Further, Polθ has been explored as a polymerase able to incorporate canonical and non-canonical ribonucleotides and deoxyribonucleotides at the 3’-terminus of ssDNA and RNA^36^ , and the steric gate mutation (E2335G) of Polθ has been characterized as an efficient template-independent RNA polymerase incorporating various 2’-O modified NTPs^37^and a powerful DNA-dependent RNA polymerase.^38^

Building on these findings, we hypothesized that PolθE2335G would enable efficient untemplated RNA synthesis with various chemically modified ribonucleotides such as the 3’-O-aminoxy nucleotides.

### Assay For Nucleotidyl Transferase Activity Using PrimPol and Polθ Variants

Initial experiments showed that native PrimPol exhibited “terminal transferase” untemplated primer extension activity with both standard deoxyribonucleotides (dNTPs) and ribonucleotides (NTPs). These also showed that the wild-type mimiPrimPol (pLB001) incorporated ribonucleotides with various moieties attached to the 3’ end, such as methoxy, azido and aminoxy (**Fig. 5**). The mutants of mimiPrimPol showed diminished activity, even with standard nucleotides. 2’, 3’-Dideoxyribonucleoside triphosphates were not accepted as substrates in any of these reactions. Further, no improvement in nucleotidyl transferase function was observed in the mutants compared to the wild-type mimiPrimPol (**Fig. 5**).

**Figure 5.**
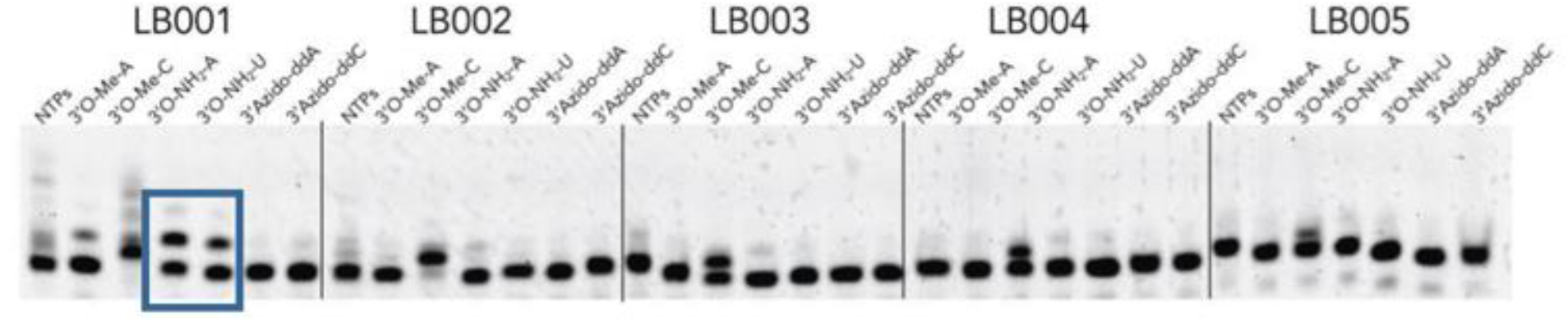
3’-O modified triphosphates (methoxy, aminoxy, azido) were tested with a variety of mutants, at 30 °C for 30 minutes. Only wildtype mimiPrimPol accepted 3’-O aminoxy derivatives with significant level of efficiency (blue box).

We asked whether the activity of native mimiPrimPol with 3’-aminoxyribonucleoside triphosphates could be modulated by changing the buffer or the divalent cation. Here, various divalent ions, pH levels, ionic strengths, detergents, and reducing agents were explored. This involved screening with just one aminoxy nucleoside triphosphates, ATP-3’-O-NH_2_, with NTPs as positive controls.

Notable from the data in **Fig. 6**, Mn^2+^ improved incorporation. Further, while Co^2+^ helps, it helps only at pH 7.5; at pH 8.5, Co^2+^ loses its ability to assist the incorporation. Additionally, the concentration of KCl significantly improves nucleotide addition; a concentration of 100 mM proved to be better than 150 mM. DTT substantially enhances the addition. BSA does not show any effect.

**Figure 6.**
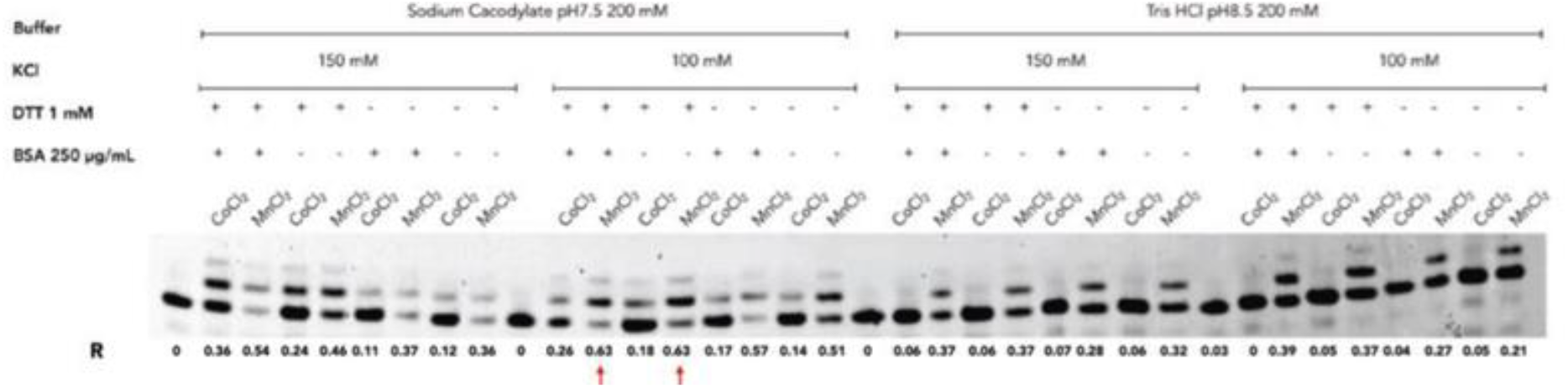
Incorporation of 3’-O-NH2 ATP with different buffers at pH 7.5 (left) and pH 8 (right) in the presence of different divalent ions using wildtype (LB001) mimiPrimPol and a FAM-labeled single stranded RNA primer. Extensions were for 30 min at 30 °C, with EDTA used as a quench prior to loading on a sequencing gel.

The results with Polθ were different and, ultimately, better. Initial experiments showed that the steric-gate variant efficiently extended a RNA primer with the aminoxy derivatives of C, U, and G. Curiously, only A was not accepted as a substrate (**Fig. 7**). The steric-gate mutant efficiently extended the RNA primer with all four canonical ribonucleotides as expected (**Fig. 7**).

**Figure 7.**
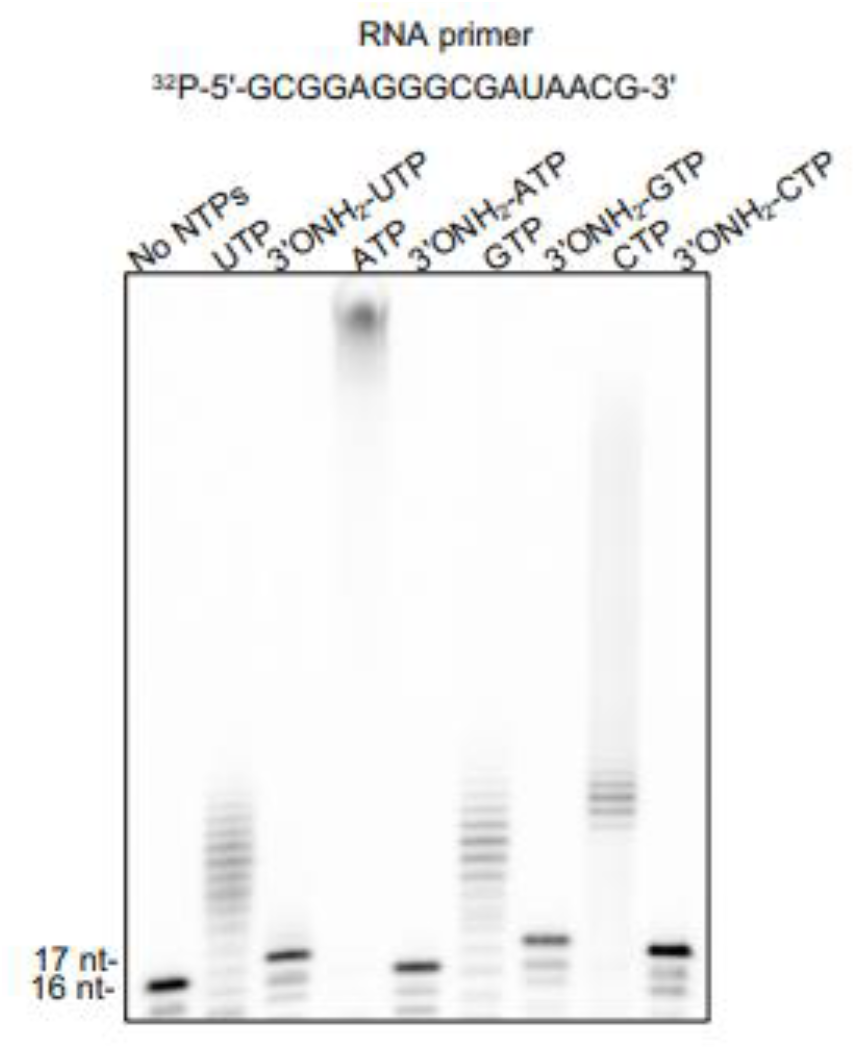
Extension of a RNA primer by Polθ E2335G with 3’-OH and 3’-aminoxy ribonucleoside triphosphates. Ladders were observed with all four standard nucleoside triphosphates having a free 3’-OH group, as expected. With a short RNA primer (^32^P-5’-GCGGAGGGCGAUAACG-3’) and 3’-ONH_2_ blocking group and the, controlled single nucleotide extension as observed with 3’-ONH_2_-CTP, 3’-ONH_2_-UTP, and 3’-ONH_2_-GTP, but not 3’-ONH_2_-ATP pH 7.5.

We envisaged that optimizing PolθE2335G enzymatic conditions and the primer nucleic acid construct may improve incorporation of 3’-aminoxy-AMP. For example, Polθ has been shown to exhibit both template-independent and dependent extension of ssDNA^36a^. The template-dependent process involves a “snap-back” replication mechanism whereby Polθ forms a transient hairpin loop that enables transient pairing of the 3’ ssDNA terminal base with a complementary base upstream along the ssDNA, resulting in a minimally paired pseudo primer-template structure. Thus, we envisaged having a 5’-polyT tail on the primer might improve improved incorporation of 3’-aminoxy-AMP.

We identified optimal conditions in which PolθE2335G incorporates all four 3’-aminoxy-NTPs using ssDNA as a primer. The conditions included Mn^2+^, relatively high concentrations of PolθE2335G and 3’-aminoxy-NTP substrates, and an ssDNA primer substrate containing a poly-T sequence at the 5’ terminus as predicted (**Fig. 8**). Using these optimized conditions, we demonstrate highly efficient PolθE2335G addition of all four 3’-aminoxy ribonucleotides (**Fig. 8**).

**Figure 8.**
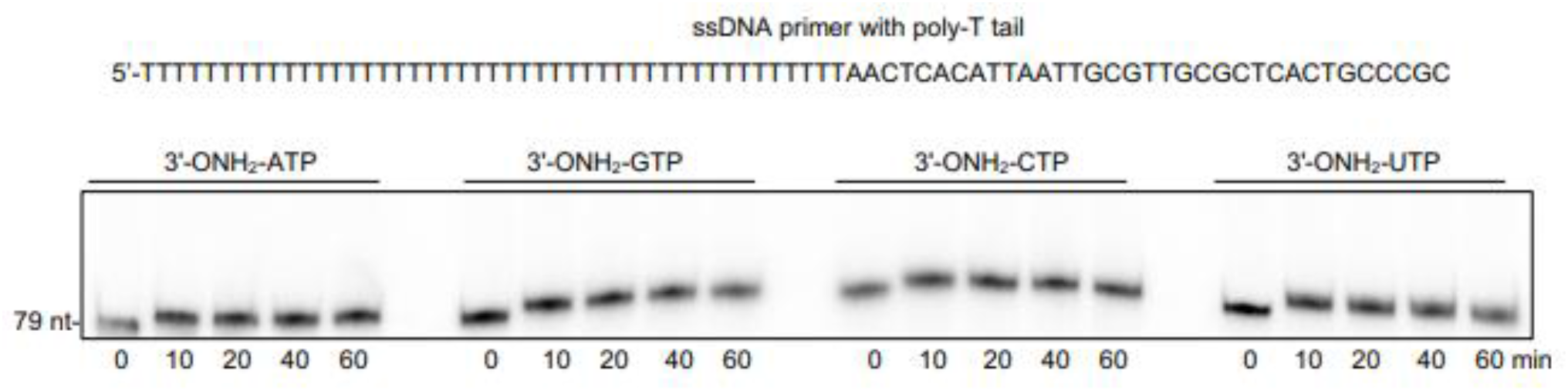
PolθE2335G incorporates all four 3’-aminoxy ribonucleotides under optimal conditions. Denaturing gel showing ssDNA extension by PolθE2335G in the presence of manganese , the indicated nucleoside triphosphate analogs, and a RNA primer with a poly-T appendage at the 5’-terminus pH 8.0. TTTTTTTTTTTTTTTTTTTTTTTTTTTTTTTTTTTTTTTTTTTTAACTCACATTAATTGCGTTGCGCTCACTGCCCGC

### Three cycles of enzyme-dependent RNA synthesis with Pol θ

The extension reactions shown above were sufficiently efficient to allow us to demonstrate proof-of-principle for multiple cycles of 3’-amonoxy-NMP addition, resulting in step-wise synthesis of two different RNA sequences 3 nt in length (**Fig. 9**). Here, PolθE2335G was incubated with ssDNA (5’-TTTTTTTTTGTGTGTGTGTGTGTGT ATTAATGCGTCACTGCG-3’) and 3’-aminoxy-CTP under optimal conditions with Mn^2+^ for 10 min. After purification of the ssDNA and removal of the 3’-aminoxy blocking group via incubation with aqueous sodium nitrate buffered at pH 6.0, a second addition step with 3’-aminoxy-ATP (**Fig. 9A**) or 3’-aminoxy-CTP (**Fig. 9B**) was performed. The ssDNA was again purified followed by the 3’-aminoxy deblocking step. Finally, a third addition step was performed with 3’-aminoxy-CTP. These results are a proof-of-principle showing that PolθE2335G can be used to perform step-wise enzymatic synthesis of two different 3 nt RNA sequences where a ssDNA primer does the initiation.

**Figure 9.**
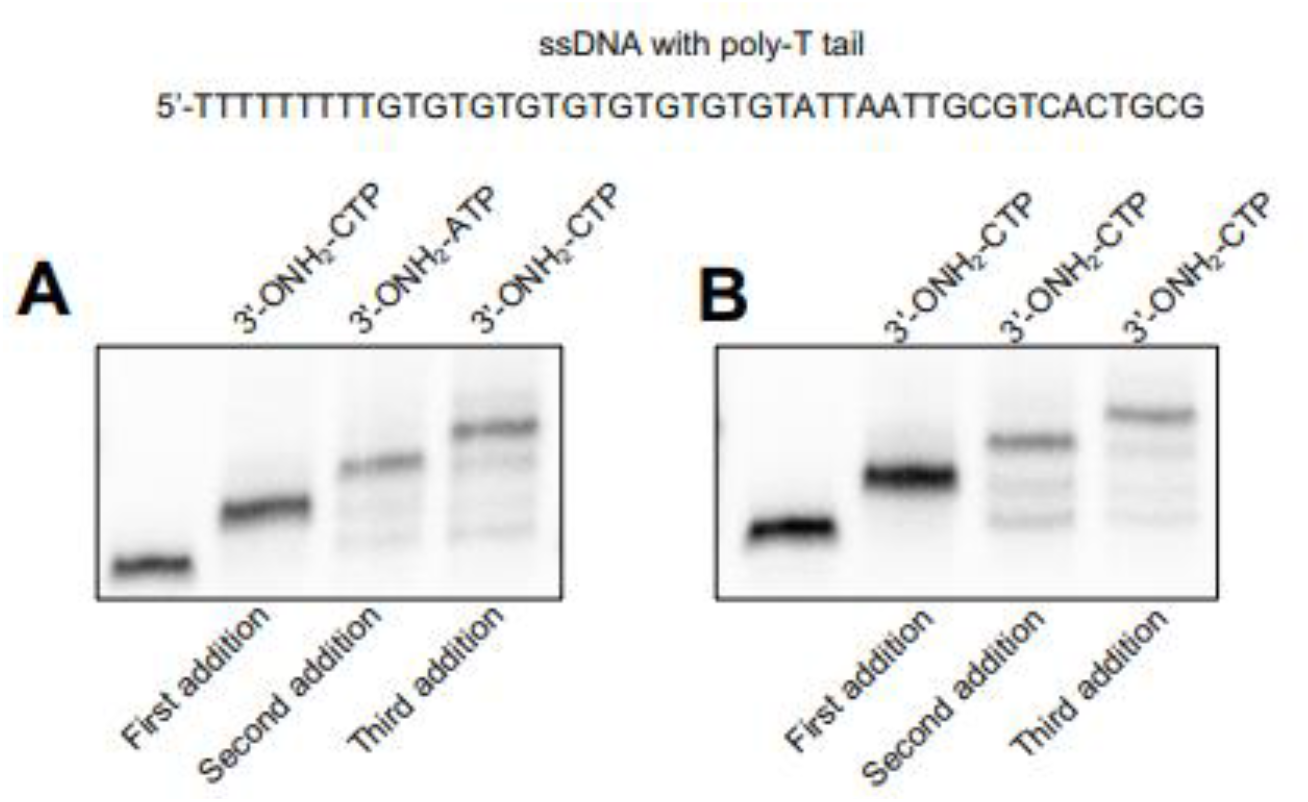
Three step-wise cycles of 3’-aminoxy-NMP addition using PolθE2335G. Denaturing gel showing step-wise addition of the indicated 3’-aminoxy-NMPs with PolθE2335G using the indicated ssDNA primer. Addition used 3’-aminoxy-CTP followed by deblocking, then 3’-aminoxy-ATP (**A**) or 3’-aminoxy-CTP (**B**) followed by deblocking, then 3’-aminoxy-CTP followed by deblocking, to a primer with a short 5’-oligoT extension (TTTTTTTTT GTGTGTGTGTGTGTGT ATTAATGCGTCACTGCG). Deblocking was performed with aqueous sodium nitrate buffered at pH 6.0. pH 8.0 for addition steps

### Scope and Utility

Preliminary data supporting a new approach for enzyme-dependent RNA synthesis are provided through a combination of synthetic organic chemistry, enzyme engineering, buffer selection, and divalent cation adjustment. Both PrimPol (preferably the native form) and Polθ (preferably the PolθE2335G variant) accept 3’-O-NH_2_ nucleoside triphosphates with various levels of efficiency.

Remarkably, the PolθE2335G variant exhibited robust acceptance of aminoxy triphosphates of cytidine, guanosine, and uridine, showcasing its substrate versatility. Notably, the optimization steps undertaken to facilitate the incorporation of 2’-ONH_2_-ATP included: (a) the strategic introduction of a 5’-polyT sequence, (b) a substantial elevation in both enzyme and nucleotide concentrations, and (c) the substitution of Mg^2+^ by Mn^2+^. These enhancements resulted in a successful adaptation of the enzyme, broadening its capacity for nucleotide incorporation.

These adjustments were adequate to support three cycles of manual cyclic reversible RNA synthesis using primer extension, adding (sequentially) C, then A or C, and then C. Cleaving the O-N bond in the 3’-O-NH_2_ group by buffered sodium nitrite, again manually, re-generates an extendable 3’-OH group, allowing the next cycle to add another selected nucleotide. This manual synthesis shows that this process is ready for instrument automation and further optimization using PolθE2335G or other related Polθ variants.

Because the PolθE2335G variant also incorporates well 2’-O modified NTPs (Ref. 37) it is likely that it will allow the synthesis of short or mid-sized 2’-O modified RNA that are resistant to nucleases.

As is common with enzymatic processes at this stage of development, the process reported here is best further co-developed with immobilization of the primer on a solid support, with reagents delivered and by-products removed via a liquid handling platform that will ultimately be used by the public. This allows further protein engineering, as necessary, to be optimized in the actual environment where it will be used.

If so developed, the chemistry and enzymology reported here holds the promise of revolutionizing the synthesis of small and mid-sized RNA molecules. The enzyme-dependent RNA synthesis circumvents the challenges posed by protecting groups, hazardous solvents, or sensitive reagents and complicated HPLC purification that often complicate standard phosphoramidite-based RNA synthesis. Consequently, this work introduces a valuable tool to the repertoire of techniques for the untemplated synthesis of RNA.

## Conclusion

Controlled enzymatic synthesis represents a potentially transformative improvement over conventional approaches to get RNA oligonucleotides. This paper delivers a synthesis of nucleotide triphosphates carrying 3’-O-aminoxy groups. The route proves to be simple and direct, providing these species for all four standard ribonucleotides. The paper further uses these to study this transient 3’-blocking group in the controlled enzymatic synthesis of RNA oligonucleotides. The cycle can be repeated by cleaving the O-N bond in the 3’-O-NH_2_ group on the extended primer with buffered sodium nitrite to re-generate an extendable 3’-OH group. Two DNA polymerases, mimiviral PrimPol and a variant of DNA polymerase theta (PolθE2335G), were shown to accept these triphosphates as substrates to add a single untemplated nucleotides to a single-strand nucleic acid primer under engineered conditions. Three cycles of addition were performed with PolθE2335G. Once transferred to a solid state platform and optimized, this combination has the potential to revolutionize how we make RNA oligonucleotides with defined sequences.

## Supporting information

https://drive.google.com/file/d/1Wf-D3DYZ3dXV26lp4vD4OBBahH-ahet8/view?usp=drive_link

## Acknowledgements

This work was supported by funding from the NHGRI Grant 1R01HG011669-01 (S.A.B., R.P.) and 1R35GM152198-01 (R.P.).

## Notes

### Competing Interest Statement

The authors have declared no competing interest.

