## Supplementary material for "Template-Independent Enzymatic RNA Synthesis": https://drive.google.com/file/d/1Wf-D3DYZ3dXV26lp4vD4OBBahH-ahet8/view?usp=drive_link

Supporting Information

Table of Contents

I. Supplementary Schemes and Figures ______________________________ S 2

II. Supplementary Synthetic Methods _______________________________ S 3-23

III. Supplementary Biological Methods _________________________ S 23

IV Supplementary Biological Methods ____________________ S 23-26

VI. References ______________________________________________________ S 27

V Supplementary 1H NMR and Mass Spectrum

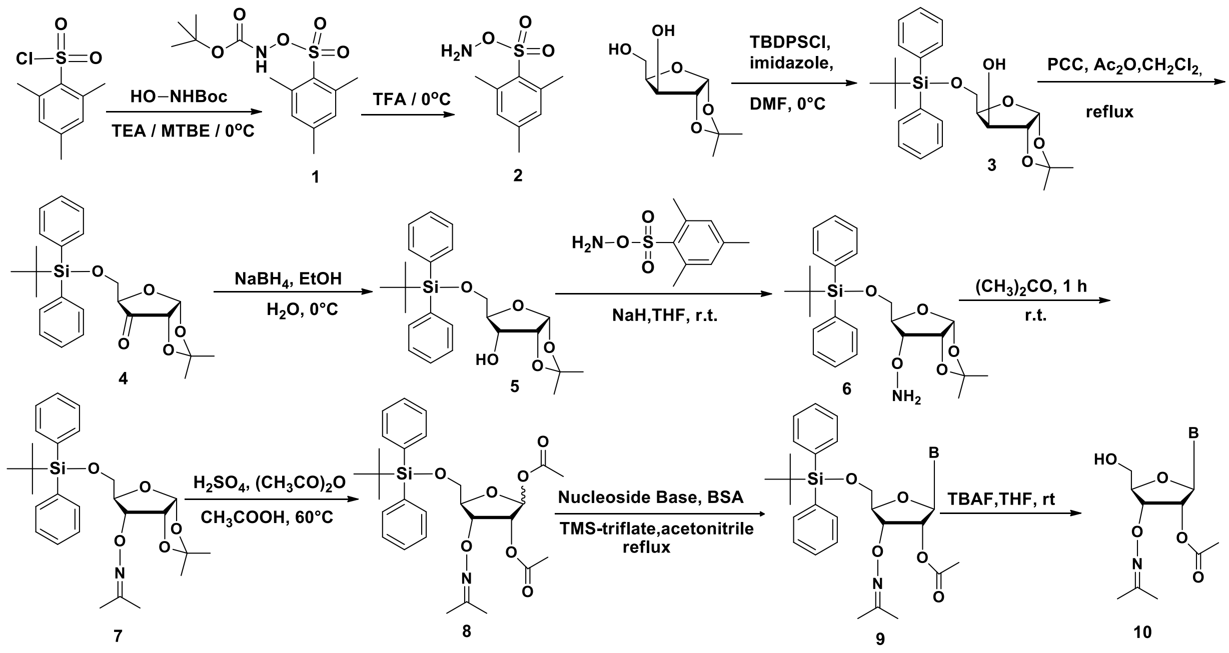

| 9 | B |
|---|---|
| a | $C^{Bz}$ |
| b | $A^{Bz}$ |
| c | $G^{ibu,DPC}$ |
| d | U |

| 10 | B |
|---|---|
| a | $C^{Bz}$ |
| b | $A^{Bz}$ |
| c | $G^{ibu,DPC}$ |
| d | U |

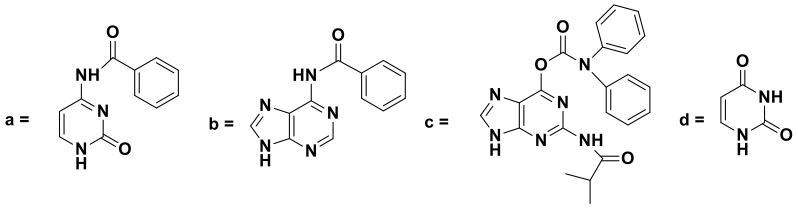

**Scheme No 1**

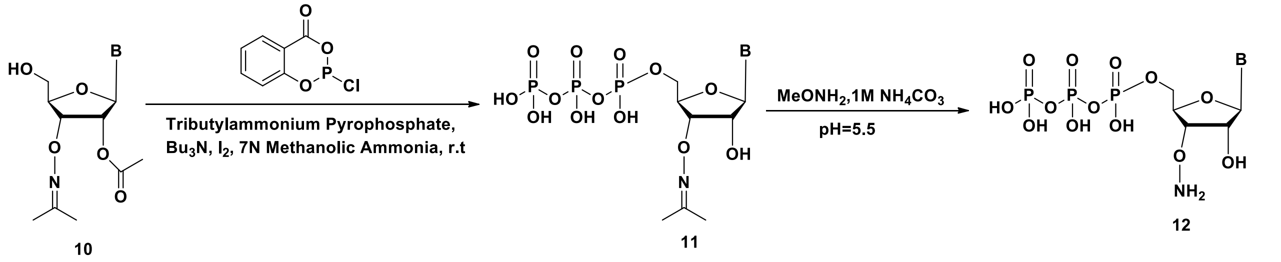

| 12 | B |
|---|---|
| a | C |
| b | A |
| c | G |
| d | U |

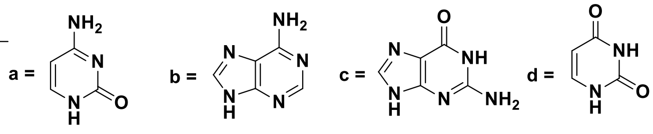

**Scheme No 2**

### Materials and General Methods

Chemical reagents were sourced from commercial suppliers and were employed without additional purification, unless otherwise specified. Anhydrous solvents were procured from commercial providers. All chemical reactions were conducted under a nitrogen atmosphere in glassware that had been oven-dried. Analytical thin-layer chromatography was carried out using glass-backed SiliCycle® silica gel 60 Å plates and was visualized either by staining with Cerium Molybdate Stain or by absorbance of UV light. Flash chromatography was performed utilizing SiliCycle® SilicaFlash® P60 silica gel. Solvent removal was accomplished under reduced pressure employing a rotary evaporator.

1H-NMR spectra were acquired utilizing Varian-300 with chemical shifts reported in parts per million (δ), and coupling constants (J) reported in hertz (Hz). High-resolution electrospray ionization (HRMS) mass spectra were acquired at the University of Florida (NIH S10 ODO21758-01A1). High-performance liquid chromatography (HPLC) was conducted using a Thermofisher instrument UHPLC equipped with an ion exchange column.

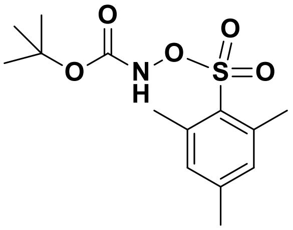

**1**

### N-Boc-O-mesitylenesulfonylhydroxylamine (Boc-MSH). (1)[i]

2-Mesitylenesulfonyl chloride (200 g, 917 mmol) and tert-butyl N-hydroxycarbamate (122.11g, 917 mmol) were dissolved in m-methyl tert-butyl ether (1L). The mixture was purged with nitrogen and cooled to 0 °C. Triethylamine (127 mL, 917 mmol) was added dropwise with stirring at 1 °C. The mixture was stirred for 2 h after addition. The reaction was monitored by TLC (n-hexane-ethyl acetate 7:3). After 2 h, $Et_3NHCl$ was filtered at the pump and washed with MTBE (1 L). Liquid phase was concentrated in Vacuo at 20 °C until ~250 mL of MTBE was left. n-Hexane (2 L) was added at 20 °C. After 5 min stirring a white solid appeared, and it was filtered and washed with n-hexane (500 mL). The liquid phase was concentrated, and more n-hexane was added (1 L). The solid was filtered and combined with the first one to give N-Boc-O-mesitylenesulfonylhydroxylamine **(1)**, (180 g, 70% yield) as a white solid. $^{1}$H NMR (300 MHz, $CDCl_3$) 7.66 (s, 1 H), 6.99 (s, 2 H), 2.68 (s, 6 H), 2.32 (s, 3 H), 1.32 (s, 9 H)

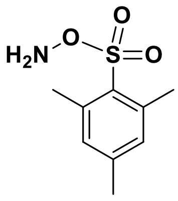

**2**

**O-Mesitylenesulfonylhydroxylamine (MSH) (2)**.

Trifluoroacetic acid (75 mL) was cooled to 2 °C. N-Boc-O-mesitylenesulfonylhydroxylamine **(1)** (20 g, 63.5 mol) was added portion wise over 1 h. Reaction was stirred at 2 °C for 90 min and monitored by TLC (n-hexane/ethyl acetate 8:2). After 90 min, TLC showed completed conversion. Cold water was added (~500 mL) followed by water (1 L). A white solid appeared. After 15 min, the solid was filtered, washed with water (20 L) until the pH ≈ 7, and dried of excess water for 15 min. O-Mesitylenesulfonylhydroxylamine (MSH, **2**) (11 g, 84% yield) was isolated as a white solid. $^{1}$H NMR (300 MHz, $CDCl_3$) 8.00 (s, 2 H), 6.92 (s, 1 H), 6.84 (s, 1 H), 2.59 (s, 3 H), 2.49 (s, 3 H), 2.27 (s, 3 H).

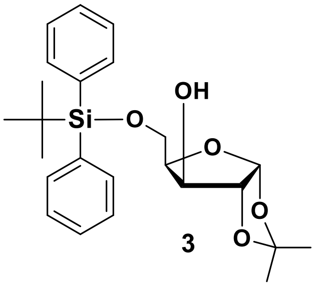

**3**

**5’-O-TBDPS-1’,2’-O-isopropylidene-α-D-xylofuranose (3)**[ii].

To a solution of TBDPS-Cl (36 mL, 144 mmol) in 50 mL of DMF at 0 ˚C was added 1,2-O-isopropylidene-α-D-xylofuranose (22.8 g, 120 mmol) and imidazole (11.4 g, 168 mmol).

After stirring for 10 min at 0 ˚C, the reaction mixture was warmed to r.t. and allowed to stir for 1 h. The reaction mixture was poured into 150 mL of ice water and diluted with 250 mL of EtOAc. After extracting the organic layer, it was washed sequentially with 150 mL of sat. $NH_4Cl$, 150 mL of sat. $NaHCO_3$, and 150 mL of brine. The resulting product was then dried over solid $Na_2SO_4$, filtered, concentrated in vacuo, and purified by silica gel column chromatography. (hexanes:EtOAc, 9:1 to 6:1 to 4:1) to afford pure **3** (28.7g, 56% yield). $^1$H NMR (300 MHz, $CDCl_3$): δ 7.77 - 7.64 (m, 4 H), 7.50 - 7.36 (m, 6 H), 6.02 (d, *J* = 3.7 Hz, 1 H), 4.55 (d, *J* = 3.7 Hz, 1 H), 4.38 (t, *J* = 2.7 Hz, 1H), 4.16 - 4.08 (m, 4 H), 1.47 (s, 3 H), 1.33 (s, 3 H), 1.06 (s, 9 H).

$^{13}$C NMR (75MHz, $CDCl_3$) : δ 135.7, 135.5, 132.5, 132.0, 130.0, 127.9, 111.5, 105.0, 85.5, 78.5, 77.5, 77.1, 76.8, 76.6, 62.8, 26.8, 26.7, 26.2, 19.1

HRMS $[M+NH_4]^+$ = 446.2378

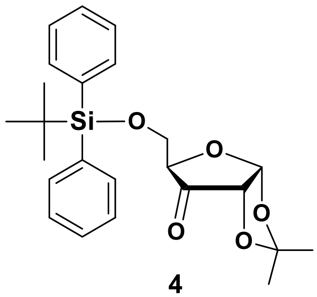


**4**

**5’-O-TBDPS-1’,2’-O-isopropylidene-α-D-xylo-3-ulose (4)**.

Pyridinium chlorochromate (18 g, 47.18 mmol) was suspended in dry methylene chloride (250 mL), acetic anhydride (21 mL, 221 mmol) was added. To this solution, 5’-O-TBDPS-1’,2’-O-isopropylidene-α-D-xylofuranose **(3)** (28.7 g, 67.05 mmol) in dry methylene chloride (100 mL) was added, and the mixture was refluxed under dry condition for 3 h. The mixture was filtered through celite, which was washed well with DCM. Evaporation gave the ketone as a colorless syrup and purified by silica gel column chromatography (hexanes: EtOAc, 4:1 to 1:1) to afford pure **4** (16 g, 57% yield). $^1$H NMR (300 MHz, $CDCl_3$): δ 7.72 - 7.65 (m, 2 H), 7.64 - 7.57 (m, 2 H), 7.48 - 7.35 (m, 6 H), 6.26 (d, *J* = 4.5 Hz, 1 H), 4.43 (d, *J* = 4.5 Hz, 1 H), 4.39 (d, *J* = 1.1 Hz, 1 H), 3.96 - 3.81 (m, 2 H), 2.10 (s, 1 H), 1.47 (d, *J* = 2.1 Hz, 6 H), 1.00 (s, 9 H).

$^{13}C$ NMR (75MHz , $CDCl_3$) : δ 210.9, 135.8, 135.5, 134.8, 132.3, 132.2, 130.0, 128.1, 127.9, 127.7, 114.2, 103.8, 81.5, 77.4, 77.1, 77.0, 76.6, 64.5, 27.7, 27.2, 26.7, 26.5, 19.1

HRMS $[M+NH_4]^+$ = 444.2198

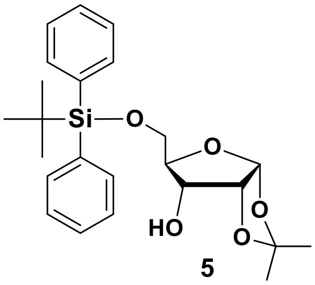


5

**5'-O-TBDPS-1',2'-O-isopropylidene-α-D-ribofuranose (5)**[2].

After dissolving **4** (25 g, 59 mmol) in 377 mL of ethanol: water (9:1), and cooling to 0 ˚C, sodium borohydride (8.0 g, 212 mmol) was added in four equivalents over 0.5 h. The reaction mixture was then allowed to stir for 3.5 h, at which point it was poured into 500 mL of water and extracted with EtOAc (500 mL). After drying the organic layer over solid $Na_2SO_4$, concentrating in vacuo, and purifying by silica gel column chromatography (hexanes: EtOAc, 15:1 to 10:1 to 5:1), the desired product, **5** (17 g, 67% yield), was obtained. $^1H$ NMR (300 MHz, $CDCl_3$): δ 7.75 - 7.65 (m, 4 H), 7.47 - 7.33 (m, 6 H), 5.85 (d, *J* = 3.5 Hz, 1 H), 4.60 (t, *J* = 4.4 Hz, 1 H), 4.22 - 4.09 (m, 1 H), 4.02 - 3.92 (m, 1 H), 3.90 - 3.79 (m, 2 H), 2.31 (d, *J* = 9.7 Hz, 1 H), 1.56 (s, 3 H), 1.38 (s, 3 H), 1.05 (s, 9 H).

$^{13}C$ NMR (75MHz , $CDCl_3$) : δ 135.7, 135.6, 135.6, 133.3, 133.2, 129.7, 127.7, 112.6, 104.2, 81.3, 78.8, 77.5, 77.1, 76.6, 71.3, 62.4, 26.9, 26.8, 26.6, 19.3

HRMS $[M+NH_4]^+$ = 446.2364

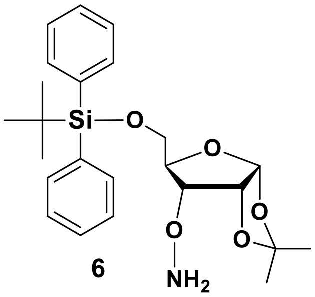


6

**5’-O-TBDPS-3’-O-(aminooxy)-1’,2’-O-isopropylidene-α-D-ribofuranose (6)**

To a stirring solution of 5’-O-TBDPS-1’,2’-O-isopropylidene-α-D-ribofuranose **(5)** (5.35 g, 12.5 mmol) in THF (30 mL) in an ice bath, sodium hydride (729 mg, 18.2 mmol, 60%w/w) was carefully added, and then the mixture was stirred for 20 min. To a resulting suspension was added a solution of O-(mesitylsulfonyl) hydroxylamine (4.20 g, 19.5 mmol) in THF (30 mL), which was used after drying over $NaSO_4$. After this was stirred for 16 h in an ice bath, ethyl acetate and water were added. The organic layer was washed with brine, dried over $NaSO_4$, filtered, and concentrated by evaporation. The residual oil was subjected to column chromatography on silica gel (hexanes: EtOAc, 4:1 to 1:4) to give **6** (3.7 g, 67% yield) as a colorless syrup. $^1$H NMR (300MHz , $CDCl_3$) 7.69 (m, 4 H), 7.46 - 7.32 (m, 5 H), 5.82 (d, *J* = 3.6 Hz, 1 H), 5.72 - 5.45 (br s, 1.4 H), 4.75 (t, *J* = 4.0 Hz, $^1$H), 4.25 (dd, *J* = 4.5, 8.5 Hz, 1 H), 4.11 (d, *J* = 7.1 Hz, 0.5 H), 4.05 - 3.91 (m, 2 H), 3.79 (dd, *J* = 3.1, 11.5 Hz, 1 H), 1.57 (s, 3 H), 1.38 (s, 3 H), 1.05 (s, 9 H).

$^{13}$C NMR (75MHz , $CDCl_3$) : δ 135.7, 135.6, 133.5, 133.3, 129.7, 127.7, 112.9, 104.3, 82.4, 79.0, 78.6, 77.5, 77.0, 76.6, 62.8, 26.9, 26.8, 26.7, 19.3

HRMS $[M+H]^+$ = 444.2222

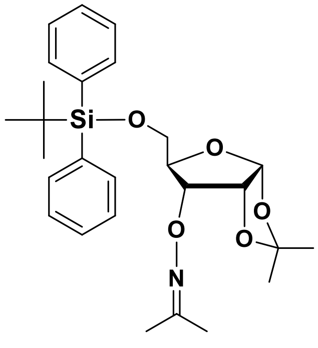


**7**

**5’-O-TBDPS-3’-O-[(propan-2-ylideneamino)oxy]-1’,2’-O-isopropylidene-α-D-ribofuranose (7)**

Acetone (30 mL) was added to a flask containing 5’-O-TBDPS-3’-O-(aminooxy)-1’,2’-O-isopropylidene-α-D-ribofuranose **(6)** (3.3 g, 7.4 mmol). The solution was stirred at r.t. for

1 h. The solvent was removed on a rotavap. The residual oil was subjected to column chromatography on silica gel (hexanes: EtOAc, 1:1) to give **7** (3.3 g, 94% yield) as a colorless syrup. $^{1}$H NMR (300MHz , $CDCl_3$) 7.69 (m, 4 H), 7.37 (m, 6 H), 5.84 (d, *J* = 3.5 Hz, 1 H), 5.29 (s, 0.5 H), 4.77 (t, *J* = 4.0 Hz, 1 H), 4.62 ( dd, *J* = 9, 4.5 Hz, 1H) 4.14 (d, *J* = 8.0 Hz, 1 H), 4.01 - 3.95 (m, 1 H), 3.80 (dd, *J* = 3.5, 11.6 Hz, 1 H), 1.87 (d, *J* = 8.4 Hz, 6 H), 1.56 (s, 3 H), 1.38 (s, 3 H), 1.04 (s, 9 H)

$^{13}$C NMR (75MHz , $CDCl_3$) : δ 156.6, 135.7, 133.6, 133.4, 129.6, 127.6, 112.9, 104.3, 80.1, 78.7, 78.6, 77.5, 77.0, 76.6, 62.6, 26.9, 26.8, 21.8, 19.3, 15.9

HRMS $[M+H]^+$ = 484.2517

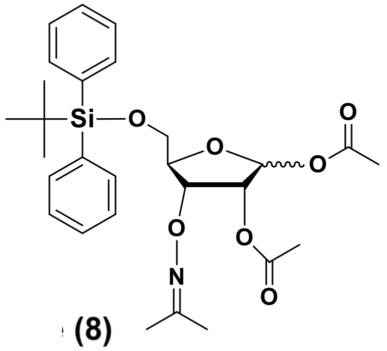


**(8)**

**3’-O-[(propan-2-ylideneamino)oxy]-5’-O-TBDPS-1’,2’-di-acetyl-α-and-β-D-erythro-pentofuranose (8)**

To a stirred solution of 5’-O-TBDPS-3’-O-[(propan-2-ylideneamino)oxy]-1’,2’-O-isopropylidene-α-D-ribofuranose **(7)** (82g,169.7 mmol) in acetic acid (1L) were added acetic anhydride (133.3 mL, 1209 mmol) and concentrated sulfuric acid (1.69 mL, 31.8 mmol). The solution was heated in an oil bath at 60°C for 18 h. The reaction was cooled and then reaction mixture was added drop-wise into a stirred aqueous solution of saturated $NaHCO_3$ (2L). The mixture was extracted with EtOAc (500 mL). Usual workup and purification by silica gel column chromatography (hexanes : EtOAc, 4:1 to 1:1) afforded 3’-O-[(propan-2-ylideneamino)oxy]-5’-O-TBDPS-1’,2’-di-acetyl-α-and-β-D-erythro-pentofuranose **(8)** (71 g, 80% yield) as a pale yellow oil.

$^{1}$H NMR (300MHz , $CDCl_3$) 7.74 - 7.65 (m, 4 H), 7.48 - 7.34 (m, 6 H), 6.20 (d, *J* = 1.5 Hz, 1 H), 5.42 (dd, *J* = 1.6, 4.8 Hz, 1 H), 5.30 (s, 1 H), 5.07 (t, *J* = 5.6 Hz, 1 H), 4.33 - 4.22

(m, 1 H), 3.97 - 3.85 (m, 1 H), 3.83 - 3.72 (m, 1 H), 2.10 (s, 3 H), 1.94 (s, 3 H), 1.86 (s, 3 H), 1.82 (s, 3 H), 1.1 (s, 9 H)

$^{13}$C NMR (75MHz , $CDCl_3$) : δ 169.7, 169.6, 156.3, 135.7, 135.6, 135.6, 133.2, 133.1, 129.7, 127.7, 98.6, 82.8, 78.7, 77.5, 77.1, 76.6, 75.3, 63.9, 63.8, 26.7, 21.6, 21.0, 20.7, 20.6, 19.3, 15.5

HRMS $[M+NH_4]^+$ = 545.2666

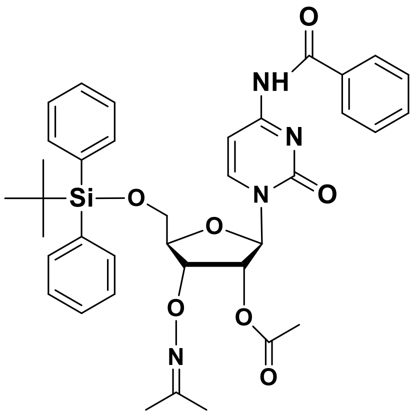

**9a**

**N$^4$-Benzoyl-5′-O-tert-butyldiphenylsilyl-2′-O-acetyl-3’-O-[(propan-2-ylideneamino)oxy]-cytidine (9a)**

Under $N_2$ atmosphere, trimethylsilyl trifluoromethanesulfonate (1.1 mL, 6.3 mmol) was added to a stirred solution of **8** (2.5 g, 4.7 mmol) and silylated *N*$^4$-benzoylcytosine (2TMS·C$^{Bz}$) (prepared from 1.2 g, 5.7 mmol of *N*$^4$-benzoylcytosine) in dichloroethane (60 mL) at room temperature. After having been refluxed for 5 h, the reaction mixture was cooled and partitioned with saturated $NaHCO_3$, and the mixture was extracted with $CH_2Cl_2$. Usual work-up and purification by $SiO_2$ column chromatography (hexanes : EtOAc, 4:1 to 0:1 to 1% MeOH in EtOAc) afforded **9a** (2.6 g, 83%) as a white foam.

$^1$H NMR (300MHz , $CDCl_3$) d = 8.77 - 8.66 (br s, 1 H), 8.34 (d, *J* = 7.5 Hz, 1 H), 7.89 (d, *J* = 7.4 Hz, 2 H), 7.70 (ddd, *J* = 1.7, 2.9, 7.7 Hz, 4 H), 7.59 (d, *J* = 7.2 Hz, 1 H), 7.54 - 7.35 (m, 8 H), 6.39 (d, *J* = 4.6 Hz, 1 H), 5.46 (t, *J* = 4.9 Hz, 1 H), 5.04 (t, *J* = 5.2 Hz, 1 H), 4.33 - 4.28 (m, 1 H), 4.12 (dd, *J* = 1.8, 11.9 Hz, 1 H), 3.88 (dd, *J* = 1.9, 11.9 Hz, 1 H), 2.10 (s, 3 H), 1.85 (d, *J* = 13.3 Hz, 6 H), 1.14 (s, 9 H)

$^{13}C$ NMR (75MHz , $CDCl_3$) : δ 169.4, 162.2, 156.6, 135.8, 135.4, 133.1, 132.6, 132.0, 130.1, 129.0, 128.1, 127.9, 127.6, 87.5, 82.6, 77.8, 77.5, 77.1, 76.6, 75.6, 63.1, 27.0, 21.7, 20.7, 19.3, 15.7

HRMS $[M+H]^+$ = 683.2922

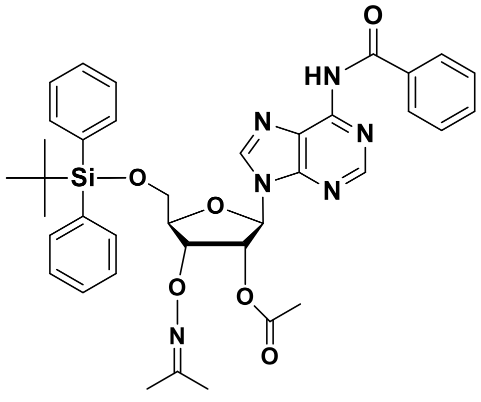

**9b**

**N6-Benzoyl-5′-O-tert-butyldiphenylsilyl-2′-O-acetyl-3’-O-[(propan-2-ylideneamino) oxy]-adenosine (9b)**

Under $N_2$ atmosphere, trimethylsilyl trifluoromethanesulfonate (0.8 mL, 4.5 mmol) was added to a stirred solution of **8** (1.8 g, 3.4 mmol) and silylated *N*4-benzoyladenosine (2TMS·$A^{Bz}$) (prepared from 992 mg, 4.1 mmol of *N*4-benzoyladenosine) in dichloroethane (60 mL) at room temperature. After having been refluxed for 3 h, the reaction mixture was cooled and partitioned with saturated $NaHCO_3$, and the mixture was extracted with $CH_2Cl_2$. Usual work-up and purification by $SiO_2$ column chromatography (hexanes: EtOAc, 4:1 to 1:1 to 1:4) afforded **9b** (1.7 g, 70%) as a white foam.

$^1H$ NMR (300MHz ,$CDCl_3$) 9.16 (s, 1 H), 8.78 (s, 1 H), 8.28 (s, 1 H), 8.01 (d, *J* = 7.3 Hz, 2 H), 7.68 (dd, *J* = 1.3, 7.7 Hz, 4 H), 7.63 - 7.46 (m, 4 H), 7.44 - 7.31 (m, 7 H), 6.46 (d, *J* = 6.9 Hz, 1 H), 5.82 (dd, *J* = 5.5, 6.8 Hz, 1 H), 5.19 (dd, *J* = 3.0, 5.3 Hz, 1 H), 4.45 - 4.40 (m, 1 H), 4.07 - 4.00 (m, 1 H), 3.92 - 3.84 (m, 1 H), 2.05 (s, 3 H), 1.90 (d, *J* = 3.9 Hz, 6 H), 1.11 (s, 9 H).

$^{13}C$ NMR (75MHz , $CDCl_3$) : δ 169.7, 164.6, 156.6, 153.0, 152.0, 149.6, 141.1, 135.8, 135.7, 135.5, 135.5, 133.7, 132.8, 132.4, 130.0, 129.9, 128.8, 127.9, 123.1, 85.2, 85.2, 83.7, 78.4, 77.5, 77.0, 76.6, 75.3, 64.0, 27.0, 21.8, 20.5, 19.3, 15.7

HRMS $[M+H]^+$ = 707.3001

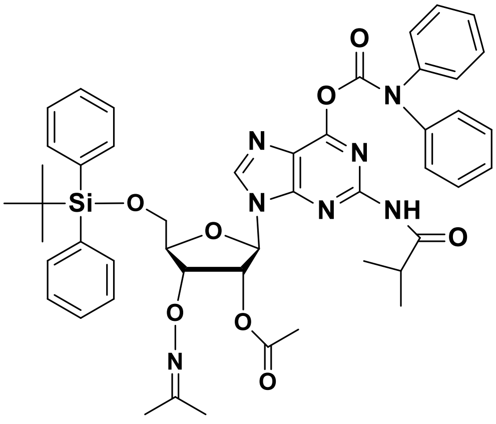

**9c**

**5′-O-tert-butyldiphenylsilyl-2′-O-acetyl-3'- O-[(propan-2-ylideneamino) oxy]- $N^2$-isobutryl-$O^6$-diphenyl-carbamoyl-guanosine (9c)**

Under $N_2$ atmosphere, trimethylsilyl trifluoromethanesulfonate (1.6 mL, 8.8 mmol) was added to a stirred solution of **8** (3.5 g, 6.6 mmol) and silylated $N^2$-isobutryl-$O^6$-diphenyl-carbamoyl guanine (3TMS·$G^{ibuDPC}$) (prepared from 3.4 g, 8.1 mmol of $N^2$-isobutryl-$O^6$-diphenyl-carbamoyl guanine) in dichloroethane (60 mL) at room temperature. After having been refluxed for 12 h, the reaction mixture was cooled and partitioned with saturated $NaHCO_3$, and the mixture was extracted with $CH_2Cl_2$. Usual work-up and purification by $SiO_2$ column chromatography (hexanes : EtOAc, 4:1 to 1:1) afforded **9c** (3.5 g, 60% yield ) as a white foam.

$^1H$ NMR (300MHz , $CDCl_3$) 8.20 (s, 1 H), 7.97 (s, 1 H), 7.68 (d, $J$ = 7.2 Hz, 4 H), 7.51 - 7.20 (m, 17 H), 6.36 (d, $J$ = 6.9 Hz, 1 H), 5.70 (t, $J$ = 6.2 Hz, 1 H), 5.15 (dd, $J$ = 2.9, 5.2 Hz, 1 H), 4.38 (d, $J$ = 2.4 Hz, 1 H), 4.12 (d, $J$ = 7.1 Hz, 1 H), 4.06 - 3.95 (m, 1 H), 3.91 - 3.81 (m, 1 H), 3.29 - 3.11 (m, 1 H), 2.05 (s, 3 H), 1.89 (d, $J$ = 6.7 Hz, 6 H), 1.28 - 1.20 (m, 6 H), 1.11 (s, 9 H)

$^{13}C$ NMR (75MHz , $CDCl_3$) : δ 169.6, 156.5, 156.2, 155.1, 152.2, 150.4, 141.8, 135.7, 135.6, 135.5, 132.7, 132.4, 129.9, 129.2, 127.9, 127.8, 121.1, 110.0, 85.0, 83.7, 78.3, 77.5, 77.4, 77.0, 76.6, 75.2, 64.0, 35.2, 27.0, 27.0, 21.8, 20.5, 19.2, 15.7

HRMS $[M+H]^+$ =884.3786

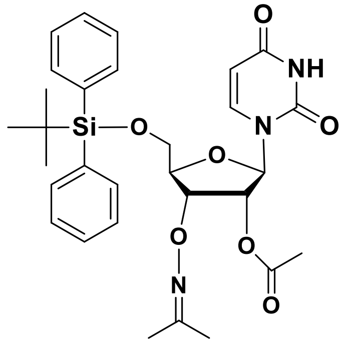

**9d**

**5′-O-tert-butyldiphenylsilyl-2′-O-acetyl-3’-O-[(propan-2-ylideneamino) oxy]-uridine (9d)**

Under $N_2$ atmosphere, trimethylsilyl trifluoromethanesulfonate (0.27 mL, 1.5 mmol) was added to a stirred solution of **8** (600 mg, 0.66 mmol) and silylated uracil (2TMS·U) (prepared from 154 mg, 1.4 mmol of uracil) in dichloroethane (60 mL) at room temperature. After having been refluxed for 1 h, the reaction mixture was cooled and partitioned with saturated $NaHCO_3$, and the mixture was extracted with $CH_2Cl_2$. Usual work-up and purification by $SiO_2$ column chromatography (hexanes : EtOAc, 4:1 to 1:1) afforded **9d** (0.350 g, 92 % yield) as a white solid.

$^1H$ NMR (300MHz , $CDCl_3$) 8.99 (br. s., 1 H), 7.78 (d, *J* = 8.2 Hz, 1 H), 7.66 (dt, *J* = 1.5, 7.9 Hz, 4 H), 7.49 - 7.34 (m, 6 H), 6.32 (d, *J* = 6.4 Hz, 1 H), 5.40 - 5.27 (m, 2 H), 5.01 (dd, *J* = 3.3, 5.4 Hz, 1 H), 4.33 - 4.23 (m, 1 H), 4.17 - 4.00 (m, 2 H), 3.86 (dd, *J* = 1.8, 11.7 Hz, 1 H), 2.09 (s, 3 H), 1.87 (s, 6 H), 1.12 (s, 9 H)

$^{13}C$ NMR (75MHz , $CDCl_3$) : δ 169.8, 163.1, 156.7, 150.5, 139.7, 135.7, 135.4, 135.3, 132.8, 131.9, 130.2, 130.1, 128.1, 128.0, 103.0, 85.6, 83.1, 78.3, 77.5, 77.1, 76.6, 74.9, 64.0, 27.0, 21.8, 21.7, 20.6, 19.4, 19.3, 15.8

HRMS $[M+H]^+$ = 580.2463

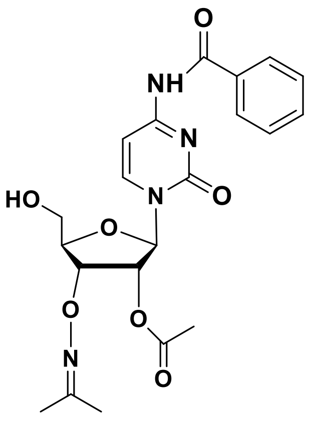

**10a**

**N[4]-Benzoyl-2′-O-acetyl-3’-O-[(propan-2-ylideneamino)oxy]-cytidine (10a)**

To a stirred solution of **9a** (2.6 g, 3.8 mmol) in dry THF (20 mL) was added to tetrabutylammonium fluoride (3.8 mL, 3.8 mmol, 1.0 M solution in THF) at room temperature. After having been stirred for 1h, the reaction mixture was concentrated under reduced pressure. The remaining residue was chromatographed on silica gel (0-4% $CH_2Cl_2$: MeOH) to afford **10a** (1.2 g, 70 % yield) as a white solid.

$^1$H NMR (300MHz , $CDCl_3$) 9.07 (br. s., 1 H), 8.34 (d, *J* = 7.5 Hz, 1 H), 7.83 (d, *J* = 7.5 Hz, 2 H), 7.61 - 7.36 (m, 4 H), 6.16 (d, *J* = 5.0 Hz, 1 H), 5.54 (t, *J* = 5.2 Hz, 1 H), 4.96 (t, *J* = 4.9 Hz, 1 H), 4.34 (d, *J* = 1.8 Hz, 1 H), 4.12 (br. s., 1 H), 4.06 - 3.95 (m, 1 H), 3.92 - 3.80 (m, 1 H), 2.03 (s, 3 H), 1.81 (d, *J* = 4.3 Hz, 6 H)

$^{13}$C NMR (75MHz , $CDCl_3$) : δ 169.7, 162.8, 156.9, 155.2, 146.0, 133.1, 133.0, 128.8, 127.8, 97.5, 89.8, 83.8, 78.4, 77.5, 77.1, 76.7, 76.6, 75.1, 61.7, 29.7, 21.7, 20.6, 15.7

HRMS $[M+H]^+$ = 445.1714

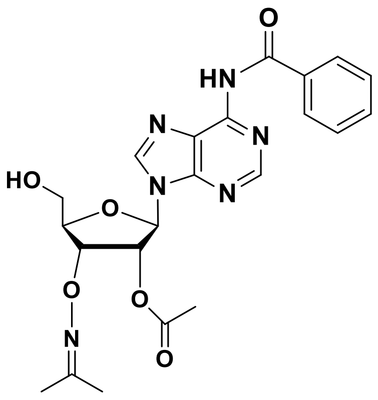

**10b**

**$N^6$-Benzoyl-2′-O-acetyl-3’-O-[(propan-2-ylideneamino) oxy]-adenosine (10b)**

To a stirred solution of **9b** (1.2 g, 1.7 mmol) in dry THF (20 mL) was added to tetrabutylammonium fluoride (2.0 mL, 2.0 mmol, 1.0 M solution in THF) at room temperature. After having been stirred for 45 minutes, the reaction mixture was concentrated under reduced pressure. The remaining residue was chromatographed on silica gel (5% (v/v) MeOH in $CH_2Cl_2$ to 9:1 $CH_2Cl_2$: MeOH) to afford **10b** (400 mg, 50% yield) as a white solid.

$^1$H NMR (300MHz , $CDCl_3$) 9.49 (s, 1 H), 8.66 (s, 1 H), 8.11 (s, 1 H), 8.00 - 7.91 (m, 2 H), 7.57 - 7.47 (m, 1 H), 7.47 - 7.37 (m, 2 H), 6.18 (d, *J* = 7.7 Hz, 1 H), 5.90 (d, *J* = 8.6 Hz, 1 H), 5.76 (dd, *J* = 5.6, 7.6 Hz, 1 H), 5.07 (dd, *J* = 1.2, 5.6 Hz, 1 H), 4.48 (d, *J* = 1.2 Hz, 1 H), 4.00 - 3.89 (m, 1 H), 3.84 - 3.68 (m, 1 H), 1.98 - 1.91 (s, 3 H), 1.91 (d, *J* = 4.5 Hz, 6 H)

$^{13}$C NMR (75MHz , $CDCl_3$) : δ 169.4, 164.9, 156.6, 152.2, 150.9, 150.3, 142.7, 133.4, 132.8, 128.8, 128.0, 124.5, 88.9, 86.1, 79.0, 77.5, 77.1, 76.7, 74.7, 63.1, 21.9, 20.4, 15.7

HRMS $[M+H]^+$ = 469.1840

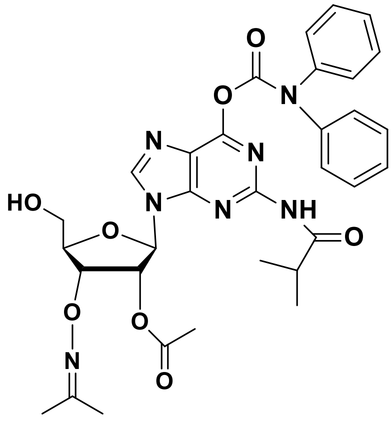

**10c**

**$N^2$-isobutryl-$O^6$-diphenyl-carbamoyl-2′-O-acetyl-3’-O-[(propan-2-ylideneamino)oxy]-guanosine (10c)**

To a stirred solution of **9c** (1.7 g, 1.9 mmol) in dry THF (20 mL) was added to tetrabutylammonium fluoride (1.9 mL, 1.92 mmol, 1.0 M solution in THF) at room temperature. After having been stirred for 45 minutes, the reaction mixture was concentrated under reduced pressure. The remaining residue was chromatographed on silica gel (0-4% $CH_2Cl_2$: MeOH) to afford **10c** (0.5 g, 42 % yield) as a white solid.

$^1$H NMR (300MHz , $CDCl_3$) 8.20 (s, 1 H), 8.09 (s, 1 H), 7.52 - 7.28 (m, 9 H), 7.28 - 7.17 (m, 3 H), 6.13 (d, *J* = 7.4 Hz, 1 H), 5.77 - 5.67 (m, 1 H), 5.14 (dd, *J* = 1.3, 5.4 Hz, 1 H), 4.77 (br. s., 1 H), 4.46 (d, *J* = 1.2 Hz, 1 H), 4.04 - 3.96 (m, 1 H), 3.83 (d, *J* = 8.6 Hz, 1 H), 2.90 - 2.75 (m, 1 H), 1.99 (s, 3 H), 1.88 (d, *J* = 2.1 Hz, 6 H), 1.23 (dd, *J* = 3.8, 6.8 Hz, 6 H)

$^{13}$C NMR (75MHz , $CDCl_3$) : δ 175.3, 171.2, 169.4, 156.5, 154.1, 151.8, 143.7, 129.2, 88.4, 85.3, 78.8, 78.7, 77.5, 77.0, 76.9, 76.6, 76.6, 74.6, 74.6, 62.7, 60.4, 36.3, 21.9, 21.8, 21.1, 20.4, 20.4, 19.3, 19.2, 19.2, 15.7, 15.6, 14.2, 14.2

HRMS $[M+H]^+$ = 646.2626

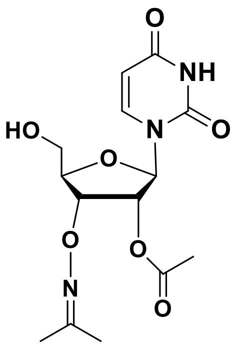

**10d**

**2′-O-acetyl-3’-O-[(propan-2-ylideneamino) oxy]-uridine (10d)**

To a stirred solution of **9d** (1 g, 1.72 mmol) in dry THF (20 mL) was added to tetrabutylammonium fluoride (2.1 mL, 2.1 mmol, 1.0 M solution in THF) at room temperature. After having been stirred for 45 minutes, the reaction mixture was concentrated under reduced pressure. The remaining residue was chromatographed on silica gel (0-5% (v/v) MeOH in $CH_2Cl_2$) to afford **10d** (500 mg, 85% yield) as a white solid.

$^1$H NMR (300MHz , $CDCl_3$) 8.91 (br. s., 1 H), 7.72 (d, *J* = 8.2 Hz, 1 H), 6.08 (d, *J* = 6.3 Hz, 1 H), 5.76 (d, *J* = 8.1 Hz, 1 H), 5.38 (t, *J* = 6.0 Hz, 1 H), 4.90 (dd, *J* = 3.5, 5.7 Hz, 1 H), 4.35 - 4.26 (m, 1 H), 4.00 - 3.80 (m, 2 H), 2.95 (t, *J* = 5.0 Hz, 1 H), 2.12 (s, 3 H), 1.92 (s, 6 H)

$^{13}$C NMR (75MHz , $CDCl_3$) : δ 170.1, 163.6, 157.1, 150.6, 141.1, 103.0, 87.7, 83.6, 78.6, 77.5, 77.0, 76.6, 74.5, 62.2, 21.7, 20.6, 15.8

HRMS $[M+NH_4]^+$ = 359.1574

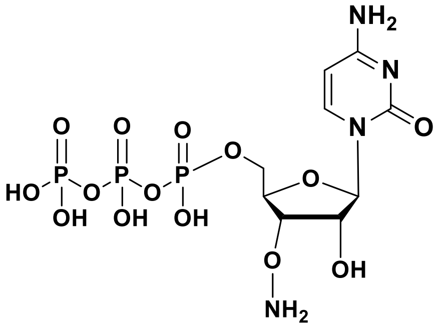

**12a**

**3'-O-Amino-cytidine-5'-triphosphate (12a)**.

To a solution of $N^4$-benzoyl-2′-O-acetyl-3’-O-[(propan-2-ylideneamino) oxy]-cytidine (250 mg, 0.56 mmol) in pyridine (3 mL) and dioxane in (6 mL) was added a solution of 2-chloro-4H-1,3,2-benzodioxaphosphorin-4-one (170mg, 0.84 mmol) in dioxane (6.0 mL) at r.t. After 15 min a mixture of tributylammonium pyrophosphate (368 mg, 0.672 mmol) in DMF (1.6 mL) and tributylamine (0.265 mL,1.12 mmol) was added. After 20 min a solution of iodine (226 mg, 0.89 mmol) and water (0.112 mL) in pyridine (5 mL) was added. After 20 min the reaction was quenched by the addition of aqueous $Na_2SO_3$ (10%, 0.5 mL). The solvents were removed in vacuo. The residue was dissolved in a mixture of water and acetonitrile (10 mL each) and kept at room temperature overnight. Volatile solvent were removed on a rotavap using vacuum at 20°C. The residue obtained was treated with 7N ammonia in methanol (25 mL) and stirred at room temperature for 24h. The ammonia was removed on a rotavap under vacuum to give an oil that was dissolved in 15 mL of water and the mixture was filtered (0.2 µm). Purification by ion-exchange HPLC (Dionex BioLC DNAPac PA-100, 22 x 250 mm, eluent A = water, eluent B = 1 M aq. $NH_4HCO_3$, gradient from 0 to 50% B in 30 min, flow rate = 10 mL/min). The product was recovered as a white solid (185.6 mg, 61 % yield) by lyophilization of fractions collected.

Fresh buffered $MeONH_2$ solution was prepared by diluting $MeONH_2$•HCl solution (3.1 M in Ar-saturated HPLC-grade water; 2 mL, 6.2 mmol) with water (HPLC, 30 mL), neutralizing it by the addition of aqueous ammonium bicarbonate buffer (1M, 5.5 mL). This solution is about 160 mM of buffered MeONH2. The final pH was 5.5 (using pH paper). Dilute terminator-oxime (185 mg, 0.34 mmol) with 7.4 mL of Ar-saturated HPLC-grade water to make a final concentration of 10 mM stock solution. Add 9.4 mL of buffered

MeONH2 solution to the 10 mM stock solution of so that the final terminator concentration is 5mM. Incubate this solution for 4 h with stirring, then neutralize the solution by adding ammonium bicarbonate buffer (1M, 8 mL). The mixture is stirred for 15 minutes at room temperature and then lyophilized. The lyophilized product was dissolved in 15 mL of HPLC water and the product was resolved by ion exchange preparative HPLC (Dionex BioLC DNAPac PA-100, 22 x 250 mm, eluent A = water, eluent B = 1 M aq. $NH_4HCO_3$, gradient from 0 to 30% B in 30 min, flow rate = 10 mL/min). The product was recovered as a white solid (100 mg, 60% yield) by lyophilization of fractions collected.

$^1$H NMR (300 MHz, $D_2O$) 7.80 (d, $J$ = 7.7 Hz, 1 H), 6.02 (d, $J$ = 7.7 Hz, 1 H), 5.89 (d, $J$ = 5.9 Hz, 1 H), 4.74 - 4.68 (m, 1 H), 4.67 - 4.59 (m, 26 H), 4.37 - 4.29 (m, 1 H), 4.28 - 4.21 (m, 2 H), 4.11 - 4.02 (m, 2 H)

$^{31}$P NMR (300 MHz, $D_2O$) -10.5 (d, $J$ = 18.5 Hz, 1P), -11.5 (d, $J$ = 19.2 Hz, 1P), -23.06 (t, $J$ = 18 Hz, 1P)

HRMS [M+H] = 499.0027

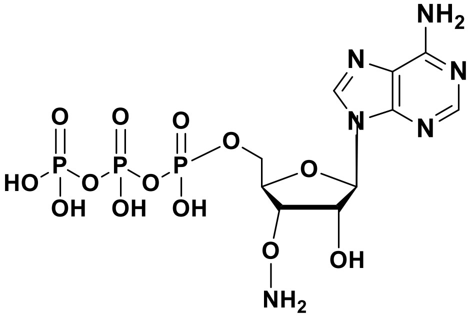


**12b**

**3'-O-Amino- adenosine-5'-triphosphate (12b)**.

To a solution of N$^6$-benzoyl-2′-O-acetyl-3’-O-[(propan-2-ylideneamino) oxy]-adenosine (0.436 g, 0.93 mmol) in pyridine (3 mL) and dioxane in (6 mL) was added a solution of 2-chloro-4H-1,3,2-benzodioxaphosphorin-4-one (0.282 g, 1.4 mmol) in dioxane (6.0 mL) at r.t. After 15 min a mixture of tributylammonium pyrophosphate (0.612 g, 1.12 mmol) in DMF (5 mL) and tributylamine (0.441 mL, 1.86 mmol) was added. After 20 min a solution of iodine (0.376 g, 1.5 mmol) and water (0.187 mL) in pyridine (9.2 mL) was added. After 20 min the reaction was quenched by the addition of aqueous $Na_2SO_3$ (10%, 1.0 mL).

The solvents were removed in vacuo. The residue was dissolved in a mixture of water and acetonitrile (10 mL each) and kept at room temperature overnight. Volatile solvent were removed on a rotavap using vacuum at 20$^{o}$C. The residue obtained was treated with 7N ammonia in methanol (25 mL) and stirred at room temperature for 24h. The ammonia was removed on a rotavap under vacuum to give an oil that was dissolved in 15 mL of water and the mixture was filtered (0.2 µm). Purification by ion-exchange HPLC (Dionex BioLC DNAPac PA-100, 22 x 250 mm, eluent A = water, eluent B = 1 M aq. $NH_4HCO_3$, gradient from 0 to 40% B in 30 min, flow rate = 10 mL/min). The product was recovered as a white solid (287 mg, 0.51 mmol) by lyophilization of fractions collected.

Fresh buffered $MeONH_2$ solution was prepared by diluting $MeONH_2$•HCl solution (3.1 M in Ar-saturated HPLC-grade water; 2 mL, 6.2 mmol) with water (HPLC, 30 mL), neutralizing it by the addition of aqueous ammonium bicarbonate buffer (1M, 5.5 mL). This solution is about 160 mM of buffered MeONH2. The final pH was 5.5 (using pH paper). Dilute terminator-oxime (191 mg, 0.34 mmol) with 7.6 mL of Ar-saturated HPLC-grade water to make a final concentration of 10 mM stock solution. Add 7.6 mL of buffered MeONH2 solution to the 10 mM stock solution of so that the final terminator concentration is 5mM. Incubate this solution for 4 h with stirring, then neutralize the solution by adding ammonium bicarbonate buffer (1M, 8 mL). The mixture is stirred for 15 minutes at room temperature and then lyophilized. The lyophilized product was dissolved in 15 mL of HPLC water and the product was resolved by ion exchange preparative HPLC (Dionex BioLC DNAPac PA-100, 22 x 250 mm, eluent A = water, eluent B = 1 M aq. $NH_4HCO_3$, gradient from 0 to 30% B in 30 min, flow rate = 10 mL/min). The product was recovered as a white solid (114 mg, 65% yield) by lyophilization of fractions collected.

$^{1}$H NMR (300MHz , $D_2O$) 8.40 (s, 1 H), 8.05 (s, 1 H), 5.95 (d, *J* = 7.5 Hz, 1 H), 4.87 - 4.78 (m, 1 H), 4.65 (s, 22 H), 4.38 (d, *J* = 3.8 Hz, 2 H), 4.15 - 4.04 (m, 1 H), 4.03 - 3.93 (m, 1 H), 3.01 - 2.88 (m, 26 H), 1.13 - 0.99 (m, 38 H)

$^{31}$P NMR (300 MHz, $D_2O$) -5.35 (d, *J* = 20.5 Hz, 1P), -10.4 (d, *J* = 19.2 Hz, 1P), -21.5 (t, *J* = 19.4 Hz, 1P)

HRMS [M+H] = 523.0144

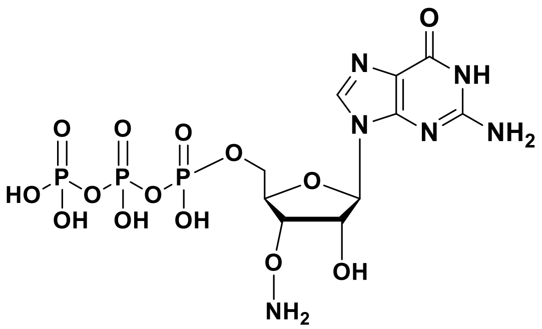

**12c**

**3'-O-Amino- guanosine-5'-triphosphate (12c)**.

To a solution of N[2]-isobutryl-O[6]-diphenyl-carbamoyl-2′-O-acetyl-3’-O-[(propan-2-ylideneamino) oxy]-guanosine (258 mg, 0.4 mmol) in pyridine (3 mL) and dioxane in (6 mL) was added a solution of 2-chloro-4H-1,3,2-benzodioxaphosphorin-4-one (132 mg, 0.65 mmol) in dioxane (6.0 mL) at r.t. After 15 min a mixture of tributylammonium pyrophosphate (0.285 g, 0.521 mmol) in DMF (5 mL) and tributylamine (0.21 mL, 0.868 mmol) was added. After 20 min a solution of iodine (0.176 g, 0.694 mmol) and water (0.086 mL) in pyridine (3.9 mL) was added. After 20 min the reaction was quenched by the addition of aqueous $Na_2SO_3$ (10%, 0.5 mL). The solvents were removed in vacuo. The residue was dissolved in a mixture of water and acetonitrile (10 mL each) and kept at room temperature overnight. Volatile solvent were removed on a rotavap using vacuum at 20°C. The residue obtained was treated with 7N ammonia in methanol (25 mL) and stirred at room temperature for 24h. The ammonia was removed on a rotavap under vacuum to give an oil that was dissolved in 15 mL of water and the mixture was filtered (0.2 µm). Purification by ion-exchange HPLC (Dionex BioLC DNAPac PA-100, 22 x 250 mm, eluent A = water, eluent B = 1 M aq. $NH_4HCO_3$, gradient from 0 to 50% B in 30 min, flow rate = 10 mL/min). The product was recovered as a white solid (154 mg, 0.27 mmol) by lyophilization of fractions collected.

Fresh buffered $MeONH_2$ solution was prepared by diluting $MeONH_2$•HCl solution (3.1 M in Ar-saturated HPLC-grade water; 2 mL, 6.2 mmol) with water (HPLC, 30 mL), neutralizing it by the addition of aqueous ammonium bicarbonate buffer (1M, 5.5 mL). This solution is about 160 mM of buffered MeONH2. The final pH was 5.5 (using pH paper). Dilute terminator-oxime (129.6 mg, 0.224 mmol) with 5.9 mL of Ar-saturated

HPLC-grade water to make a final concentration of 10 mM stock solution. Add 8.0 mL of buffered MeONH2 solution to the 10 mM stock solution of so that the final terminator concentration is 5mM. Incubate this solution for 4 h with stirring, then neutralize the solution by adding ammonium bicarbonate buffer (1M, 8 mL). The mixture is stirred for 15 minutes at room temperature and then lyophilized. The lyophilized product was dissolved in 15 mL of HPLC water and the product was resolved by ion exchange preparative HPLC (Dionex BioLC DNAPac PA-100, 22 x 250 mm, eluent A = water, eluent B = 1 M aq. $NH_4HCO_3$, gradient from 0 to 30% B in 30 min, flow rate = 10 mL/min). The product was recovered as a white solid (72 mg, 60% yield) by lyophilization of fractions collected.

$^1$H NMR (300 MHz, $D_2O$) 7.91 (s, 1 H), 5.73 (d, $J$ = 7.0 Hz, 1 H), 4.83 - 4.74 (m, 1 H), 4.69 - 4.60 (m, 27 H), 4.32 (t, $J$ = 2.4 Hz, 1 H), 4.15 - 3.96 (m, 2 H)

$^{31}$P NMR (300 MHz, $D_2O$) -10.8 (d, $J$ = 17.8 Hz, 1P), -11.5 (d, $J$ = 19.9 Hz, 1P), -21.5 (t, $J$ = 17 Hz, 1P)

HRMS [M+H] = 539.0090

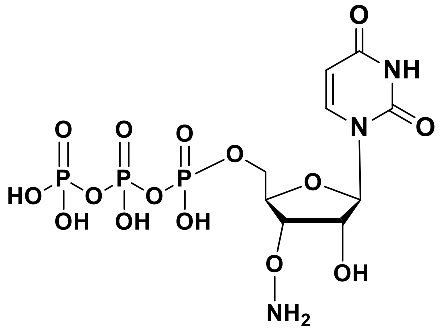


12d

**3'-O-Amino-uridine-5'-triphosphate (12d)**.

To a solution of 2′-O-acetyl-3’-O-[(propan-2-ylideneamino) oxy]-uridine (0.250 g, 0.73 mmol) in pyridine (3 mL) and dioxane in (6 mL) was added a solution of 2-chloro-4H-1,3,2-benzodioxaphosphorin-4-one (0.221 g, 1.1 mmol) in dioxane (6.0 mL) at r.t. After 15 min a mixture of tributylammonium pyrophosphate (0.480 g, 0.876 mmol) in DMF (5 mL) and tributylamine (0.346 mL, 1.45 mmol) was added. After 20 min a solution of iodine (0.295 g, 1.1 mmol) and water (0.146 mL) in pyridine (7.2 mL) was added. After 20 min the reaction was quenched by the addition of aqueous $Na_2SO_3$ (10%, 1.0 mL). The solvents were removed in vacuo. The residue was dissolved in a mixture of water and acetonitrile (10 mL each) and kept at room temperature overnight. Volatile solvent were removed on a rotavap using vacuum at 20°C. The residue obtained was treated with 7N ammonia in methanol (25 mL) and stirred at room temperature for 24h. The ammonia was removed on a rotavap under vacuum to give an oil that was dissolved in 15 mL of water and the mixture was filtered (0.2 µm). Purification by ion-exchange HPLC (Dionex BioLC DNAPac PA-100, 22 x 250 mm, eluent A = water, eluent B = 1 M aq. $NH_4HCO_3$, gradient from 0 to 50% B in 30 min, flow rate = 10 mL/min). The product was recovered as a white solid (231 mg, 0.42 mmol) by lyophilization of fractions collected.

Fresh buffered $MeONH_2$ solution was prepared by diluting $MeONH_2$•HCl solution (3.1 M in Ar-saturated HPLC-grade water; 2 mL, 6.2 mmol) with water (HPLC, 30 mL), neutralizing it by the addition of aqueous ammonium bicarbonate buffer (1M, 5.5 mL). This solution is about 160 mM of buffered MeONH2. The final pH was 5.5 (using pH paper). Dilute terminator-oxime (183 mg, 0.34 mmol) with 11.36 mL of Ar-saturated HPLC-grade water to make a final concentration of 10 mM stock solution. Add 11.36 mL of buffered $MeONH_2$ solution to the 10 mM stock solution of so that the final terminator concentration is 5 mM. Incubate this solution for 4 h with stirring, then neutralize the solution by adding ammonium bicarbonate buffer (1M, 13 mL). The mixture is stirred for 15 minutes at room temperature and then lyophilized. The lyophilized product was dissolved in 15 mL of HPLC water and the product was resolved by ion exchange preparative HPLC (Dionex BioLC DNAPac PA-100, 22 x 250 mm, eluent A = water, eluent B = 1 M aq. $NH_4HCO_3$, gradient from 0 to 30% B in 30 min, flow rate = 10 mL/min). The product was recovered as a white solid (106 mg, 63% yield) by lyophilization of fractions collected.

$^1$H NMR (300 MHz, $D_2O$) 7.66 (d, *J* = 7.8 Hz, 1 H), 5.85 (d, *J* = 6.0 Hz, 1 H), 5.75 (d, *J* = 7.8 Hz, 1 H), 4.68 - 4.50 (m, 21 H), 4.37 (br. s., 1 H), 4.23 (br. s., 1 H), 4.01 (d, *J* = 10.4 Hz, 2 H)

$^{31}$P NMR (300 MHz, D2O) -9.79 (d, *J* = 17.8 Hz, 1P), -10.5 (d, *J* = 19.6 Hz, 1P), -22.0 (t, *J* = 17 Hz, 1P)

HRMS [M+H] = 499.9869

**Mimiviral PrimPol purification**

The plasmid containing PrimPol (or variants) was used to transform One Shot BL21 Star (DE3) cells (ThermoFisher) following the manufacturers manual. Transformed cells were plated onto an LB agar plate containing Kanamycin and incubated at 37ºC overnight. After incubation, a single colony was used to start a 200 mL preculture with liquid LB medium containing Kanamycin and incubated overnight at 37ºC with constant shaking. The following day, the $OD_{600}$ was measured to start a 1 L culture at 0.1 OD units. Cells were incubated at 37ºC to reach an $OD_{600}$ of 0.6 units before induction with 1 mM IPTG final. Induction was carried out overnight at 20ºC with constant shaking. After induction, cells were harvested at 3500g for 30 minutes and resuspended in 100 mL buffer A (50 mM Hepes pH 7.5, 500 mM NaCl, 20 mM Imidazole), the buffer was supplemented with EDTA-free Pierce Protease inhibitor tablets (Thermo Scientific) as well as with 2 µL benzonase/nuclease (Millipore). Cells were then lysed with a cell disruptor at 1.4 kPa, three passages were carried out. Cell debris were pelleted at 20000 g for 30 minutes and the supernatant loaded onto a HisTrap column (Cytiva) previously equilibrated with buffer A. Elution was carried out with an Akta purifier (Cytiva) using a gradient from 0 to 100% buffer B (50 mM Hepes pH 7.5, 500 mM NaCl, 500 mM Imidazole). Fractions containing the protein were pooled and diluted 5x using 50 mM HEPES pH 7.5, the diluted and pooled fractions were loaded onto a Heparin column (Cytiva) previously equilibrated with buffer C (50 mM HEPES pH 7.5, 100 mM NaCl), elution was carried out using a gradient from 0% buffer C to 100% buffer D (50 mM HEPES pH 7.5, 1 M NaCl). Fractions containing the protein were pooled and incubated with His-TEV protease overnight at 4ºC under rotation. A second HisTrap step was carried out to separate the His-TEV protease from PrimPol, the flow through was retained and concentrated with Amicon ultra15 10k filters to obtain a volume of 500 µL. Finally, the concentrated fraction was loaded onto a Superdex 200 10/300 column (Cytiva) and the fractions containing the protein pooled.

**Nucleotidyl transferase tests with PrimPol**

To test the nucleotidyl transferase activity of PrimPol and mutants on single stranded RNA, we carried out a ssRNA extension assay. In a total volume of 10 µL, 250 nM Primpol were incubated with 100 nM 15-mer FAM-labelled RNA (FAM-CGGUGACCAUUGCGU) (Eurogentec), modified nucleotides (shown in the figures) were present at a concentration of 250 µM. The reaction buffer contained 20 mM Tris-HCl pH 8.8, 10 mM $(NH_4)_2SO_4$, 10 mM KCl, and 0.1% Triton. The reaction was started by adding 1 mM $MnCl_2$ and incubated at 30ºC for 30 minutes. The reaction was quenched by adding 10 µL of a solution of formamide containing 50 mM EDTA and bromophenol blue. RNA fragments were resolved by loading them onto a polyacrylamide sequencing gel.

DNA (5′- 3′)

486 - 5'-CACTGTGAGCTTAGTCACATTTCATCATGCAGGACAG-3’

15–5’-TTTTTTTTTTTTTTTTTTTTTTTTTTTTTTTTTTTTTTTTTTTTTAACTCACATTAATTGCGTTGCGCTCACTGCCCGC -3’

TKT45s9 – 5’-TTTTTTTTT GTGTGTGTGTGTGTGTGT ATTAATTGCGTCACTGCG-3’

RNA (5'-3')

635R - 5'-GCGGAGGGCGAUAACG-3’

**Protein**

PolθE2335G was purified as described[iii].

**3'ONH$_2$-NMP incorporation using PolθE2335G.**

Assays performed in Fig. 7.

0.1 µM PolθE2335G was incubated with 20 nM of 5'-radiolabeled 5'-GCGGAGGGCGAUAACG-3’ RNA for 30 min at 37°C in the presence of 0.2 mM of indicated NTPs or 3'ONH$_2$-analogs in a 20 µl volume of buffer A (20 mM Tris-HCl pH 7.5, 10% glycerol, 0.01% NP-40, 0.1 mg/ml BSA) with 5 mM $MnCl_2$. Reactions were terminated by the addition of 20 µl of 2X stop buffer (90% formamide, 50 mM EDTA). RNA was resolved by electrophoresis in urea polyacrylamide gels and then visualized by autoradiography.

Assays performed in Fig. 8

0.5 µM PolθE2335G was incubated with 50 nM of 5'-radiolabeled

5’-TTTTTTTTTTTTTTTTTTTTTTTTTTTTTTTTTTTTTTTTTTTTTAACTCACATTAATTGCGTTGCGCTCACTGCCCGC -3’ DNA for 0, 10, 20, 40 and 60 min at 42°C in the presence of 0.3 mM of the indicated 3'ONH$_2$-NTP analogs in a 20 µl volume of buffer A (20 mM Tris-HCl pH 8.0, 10% glycerol, 0.01% NP-40, 0.1 mg/ml BSA) with 5 mM $MnCl_2$. Reactions were terminated by the addition of 20 µl of 2X stop buffer (90% formamide, 50 mM EDTA). DNA was resolved by electrophoresis in urea polyacrylamide gels and then visualized by autoradiography.

**3'$ONH_2$-NMP step-wise incorporations using Pol$\theta$E2335G.**

Precipitation problems and inability to purify nucleic acid using ethanol due to the specific reaction and deblocking conditions caused significant loss of nucleic acid following each addition step. These technical problems prevented additional step-wise 3'-aminoxy-NMP incorporations. Primer immobilization to a solid support and automated methods are needed to overcome these technical problems and achieve quantitative step-wise enzymatic synthesis of relatively long RNA substrates using Pol$\theta$ E2335G. Assays results are shown in Fig. 9.

Step 1. PolE2335G (1 $\mu$M) was incubated with ~50 nM of 5'-radiolabeled DNA 5'-TTTTTTTTT GTGTGTGTGTGTGTGTGT ATTAATTGCGTCACTGCG-3' for 20 min at 42°C in the presence of 0.3 mM of 3'$ONH_2$-CTP in a 1000 $\mu$l volume of buffer A (20 mM Tris-HCl pH 8.0, 10% glycerol, 0.01% NP-40, 0.1 mg/ml BSA) with 5 mM $MnCl_2$ and 10 mM Methoxylamine hydrochloride (pH 7.0). The reaction was terminated by adding 5 U/ml Proteinase K, followed by incubation at room temp for 30 min and subsequent Proteinase K inactivation by incubating at 95°C for 10 minutes. the nucleic acid was purified at room temp by using RNA Clean & Concentrator-5 columns (Zymo Research) as follows: The nucleic acid sample was mixed with an equal volume of 4M Na-acetate pH 4.6 and 10 mM Methoxylamine hydrochloride, incubated and loaded onto the column followed by centrifugation at 350xg for 10 mins. The column was then washed with 600 ul of 10 mM Methoxylamine hydrochloride. Next, 10 mM TrisHCl pH 8.8 was added to the column, incubated for 1 hr and the nucleic acid was eluted by by centrifugation at 21,000xg. Deprotection steps used the buffer described in[iv] (0.7 M NaOAc and 1.0 M $NaNO_2$ pH 5.2) with sample incubation for 5 min at room temp. Next, the nucleic acid sample was purified as described above. 10% of the purified nucleic acid was measured for radioactivity by liquid scintillation counter for normalizing loading volumes for electrophoresis as described below.

Step 2. 90% of the volume of the purified nucleic acid from step 1 was used for the next addition of 3'$ONH_2$-AMP or 3'$ONH_2$-CMP as indicated. The reaction, purifications, and deprotonation were performed as described in step 1. Here, 50% of the volume of purified nucleic acid was measured for radioactivity via liquid scintillation counter for normalizing loading volumes for electrophoresis as described below.

Step 3. 50% of the volume of the purified nucleic acid from step 2 was used for the next addition of 3'$ONH_2$-CMP. The reaction, purifications, and deprotonation were performed as described above. Liquid scintillation counting and gel eletrophoresis

2 μl of each sample (purified nucleic acid) was mixed with 10 ml of Bio-Safe II solution (RPI corp.) and the total signal was measured by Liquid Scintillation Analyser Tri-Carb 4910 TR. The volumes of the samples for the electrophoresis gel were normalized based on the radioactive measurements obtained from liquid scintillation counting, resolved by electrophoresis in urea polyacrylamide gel and then visualized by autoradiography.

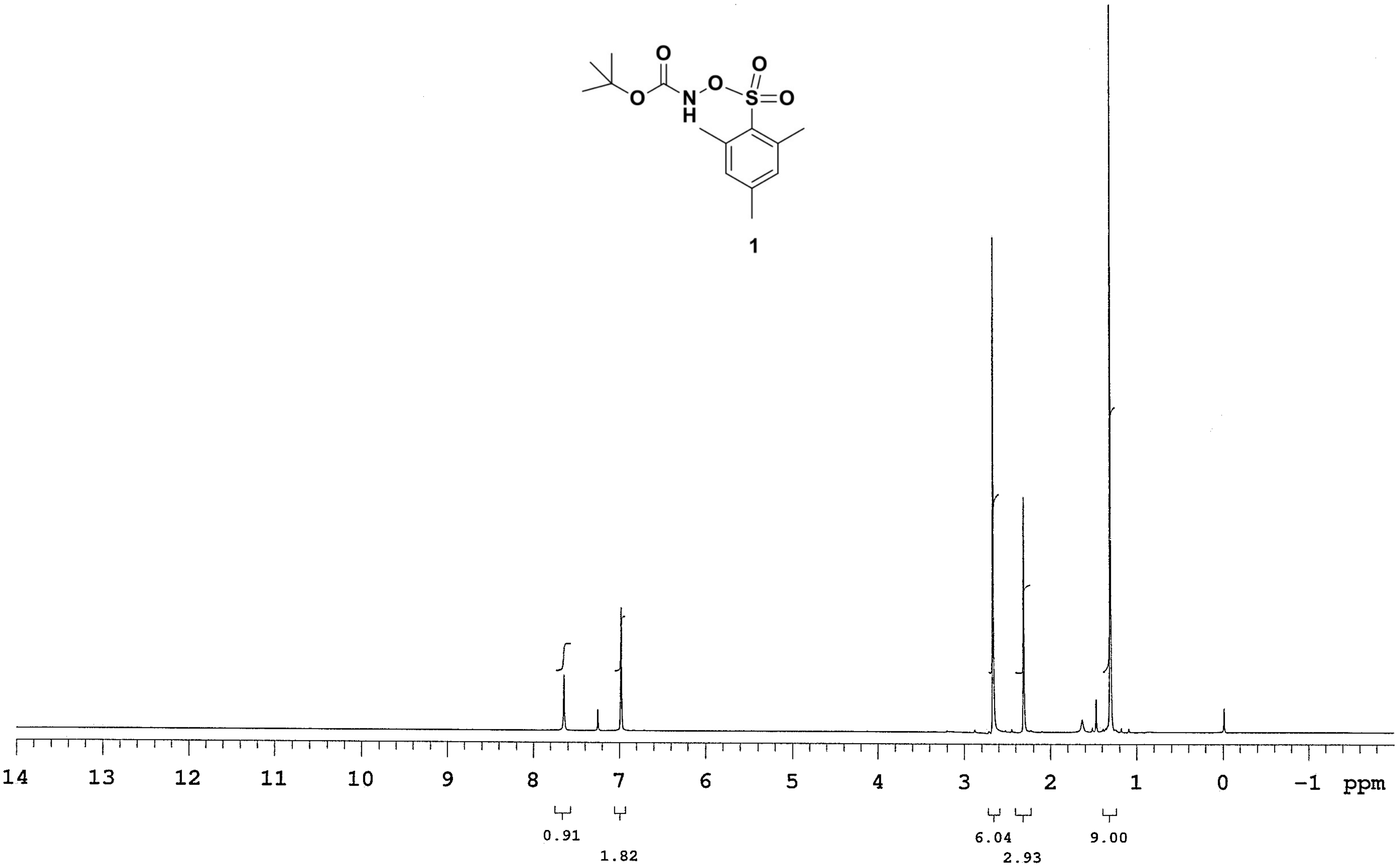

O
N
H
O
S
O
O
1
14
13
12
11
10
9
8
7
6
5
4
3
2
1
0
−1
ppm
0.91
1.82
6.04
2.93
9.00

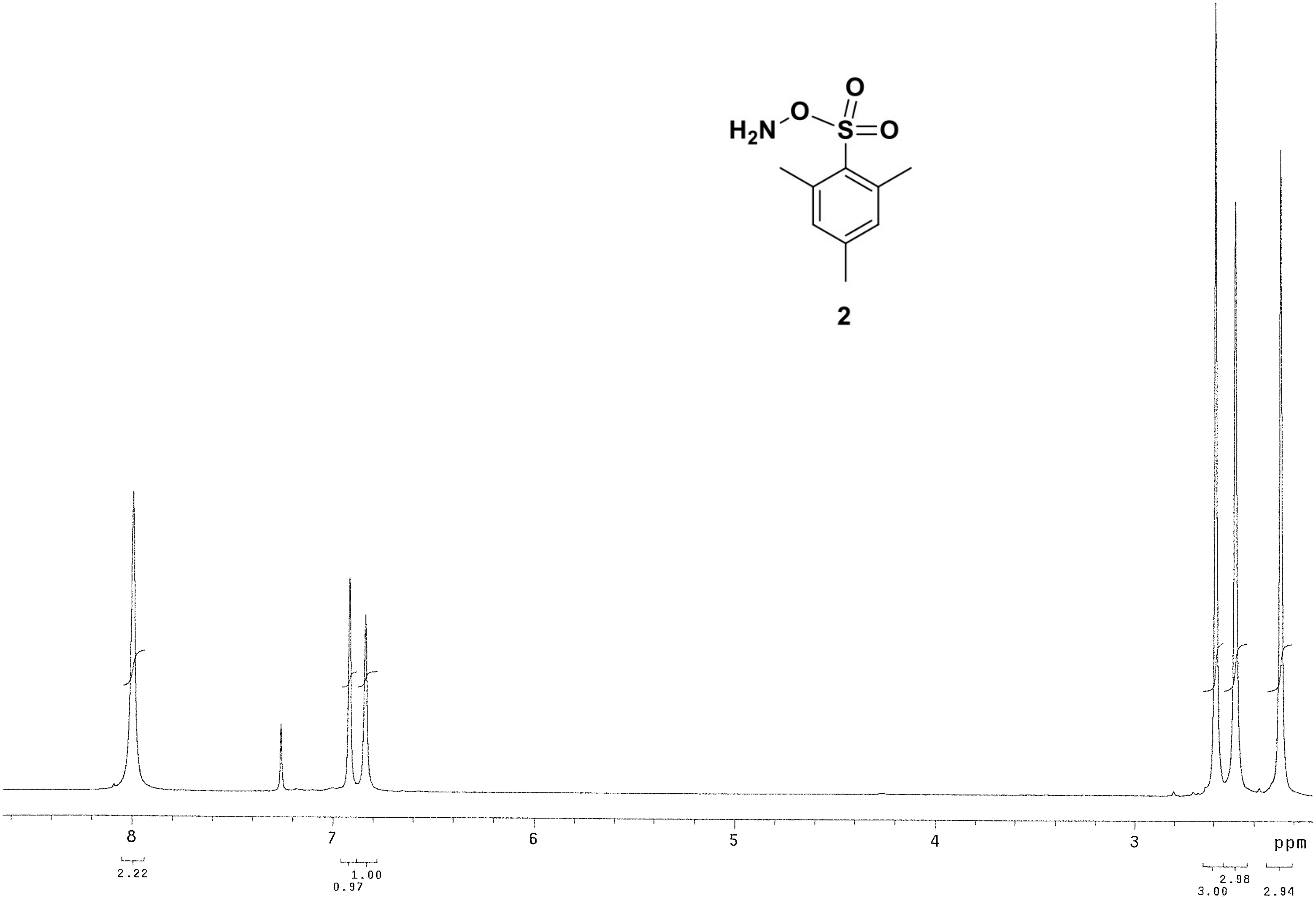


**2**

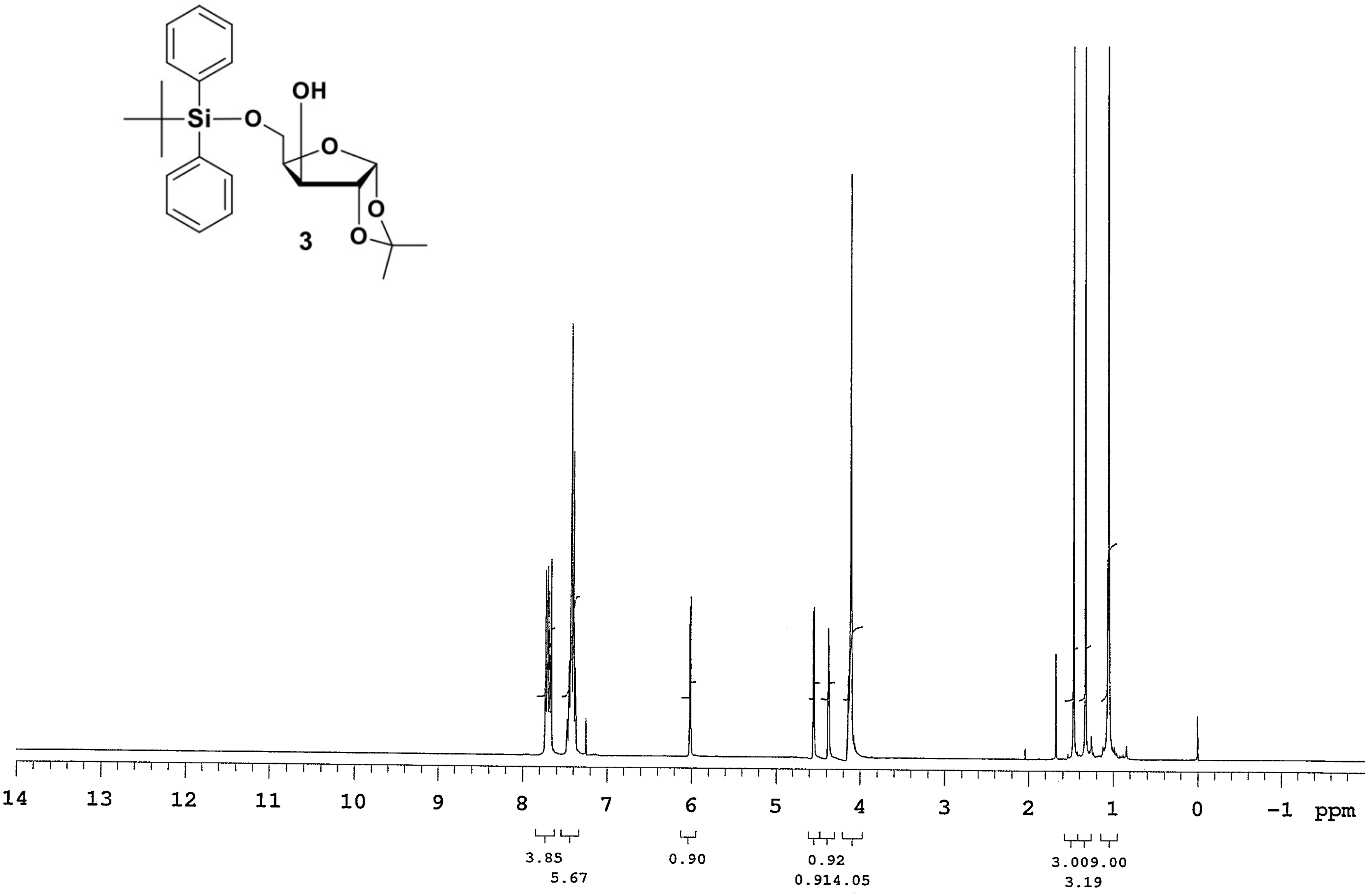

OH
Si
O
O
O
O
3
14
13
12
11
10
9
8
7
6
5
4
3
2
1
0
−1
ppm
3.85
5.67
0.90
0.92
0.914.05
3.009.00
3.19

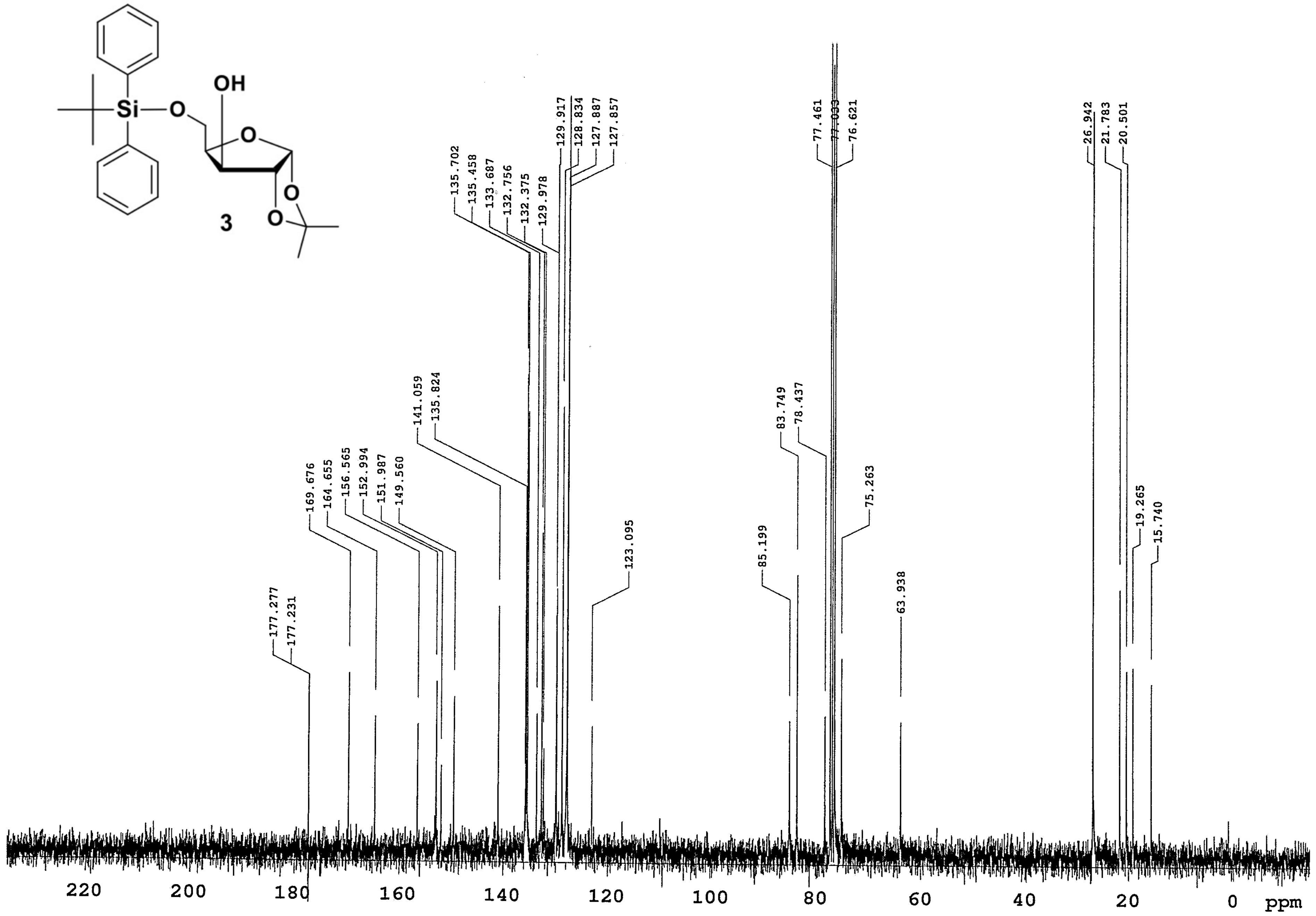
OH
Si
O
O
O
O
177.277
177.231
169.676
164.655
156.565
152.994
151.987
149.560
141.059
135.824
135.702
135.458
133.687
132.756
132.375
129.978
129.917
128.834
127.887
127.857
123.095
85.199
83.749
78.437
77.461
77.033
76.621
75.263
63.938
26.942
21.783
20.501
19.265
15.740
220
200
180
160
140
120
100
80
60
40
20
0
ppm


3

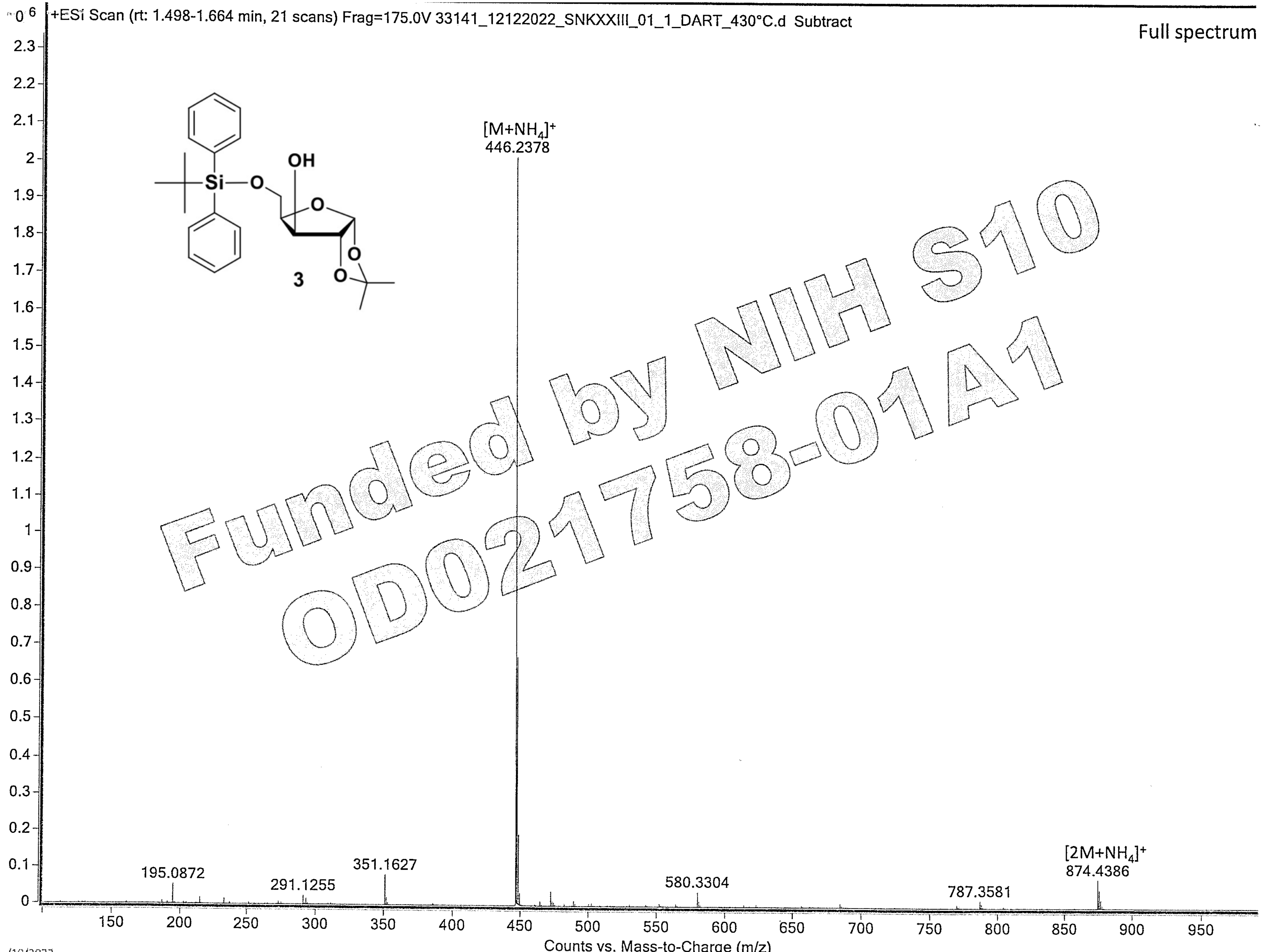


/10/2023

UF Mass Spectrometry Facility

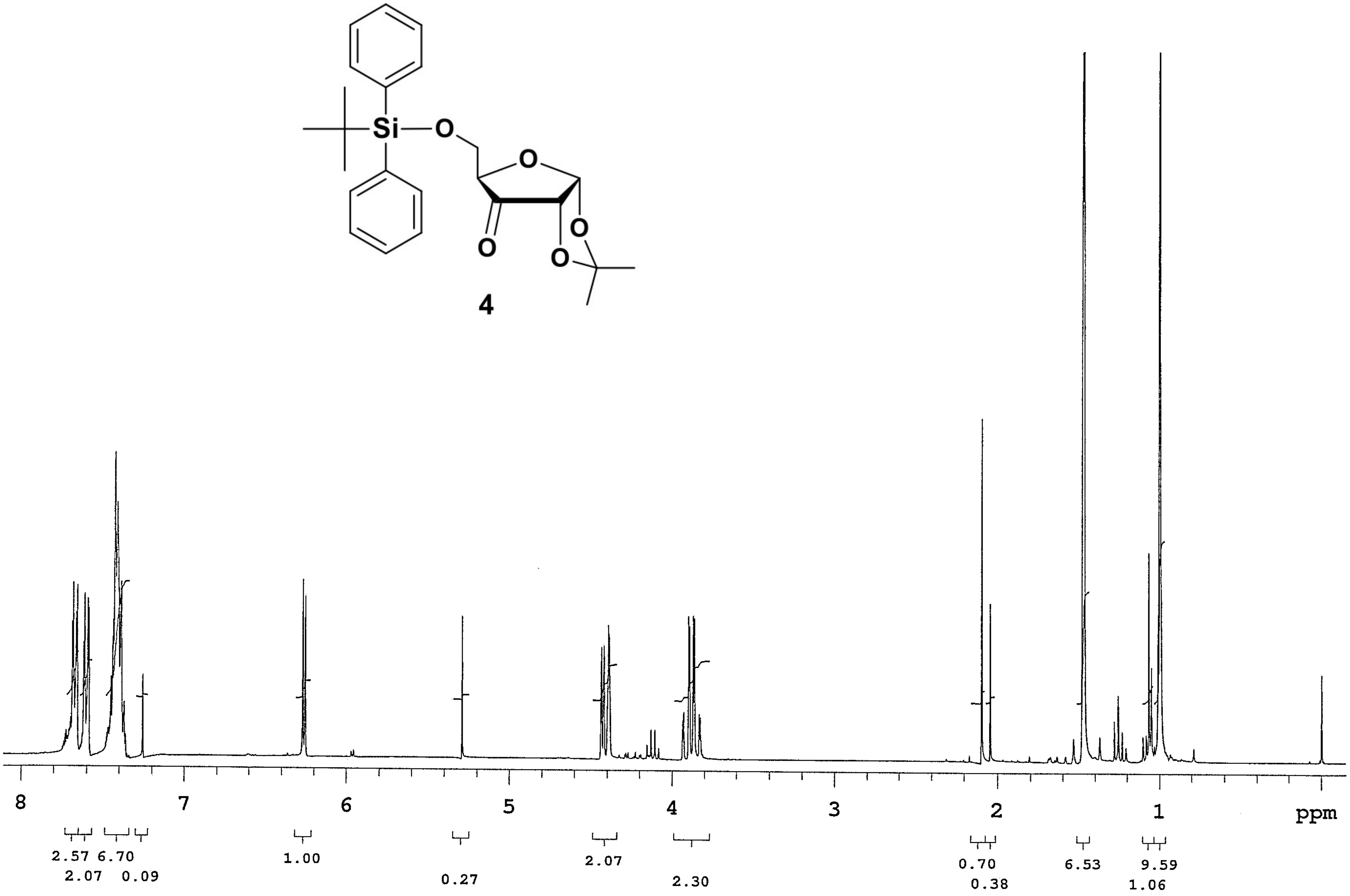
4
8
7
6
5
4
3
2
1
ppm
2.57 6.70
2.07 0.09
1.00
0.27
2.07
2.30
0.70
0.38
6.53
9.59
1.06

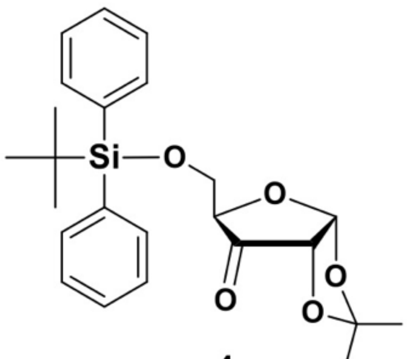

4

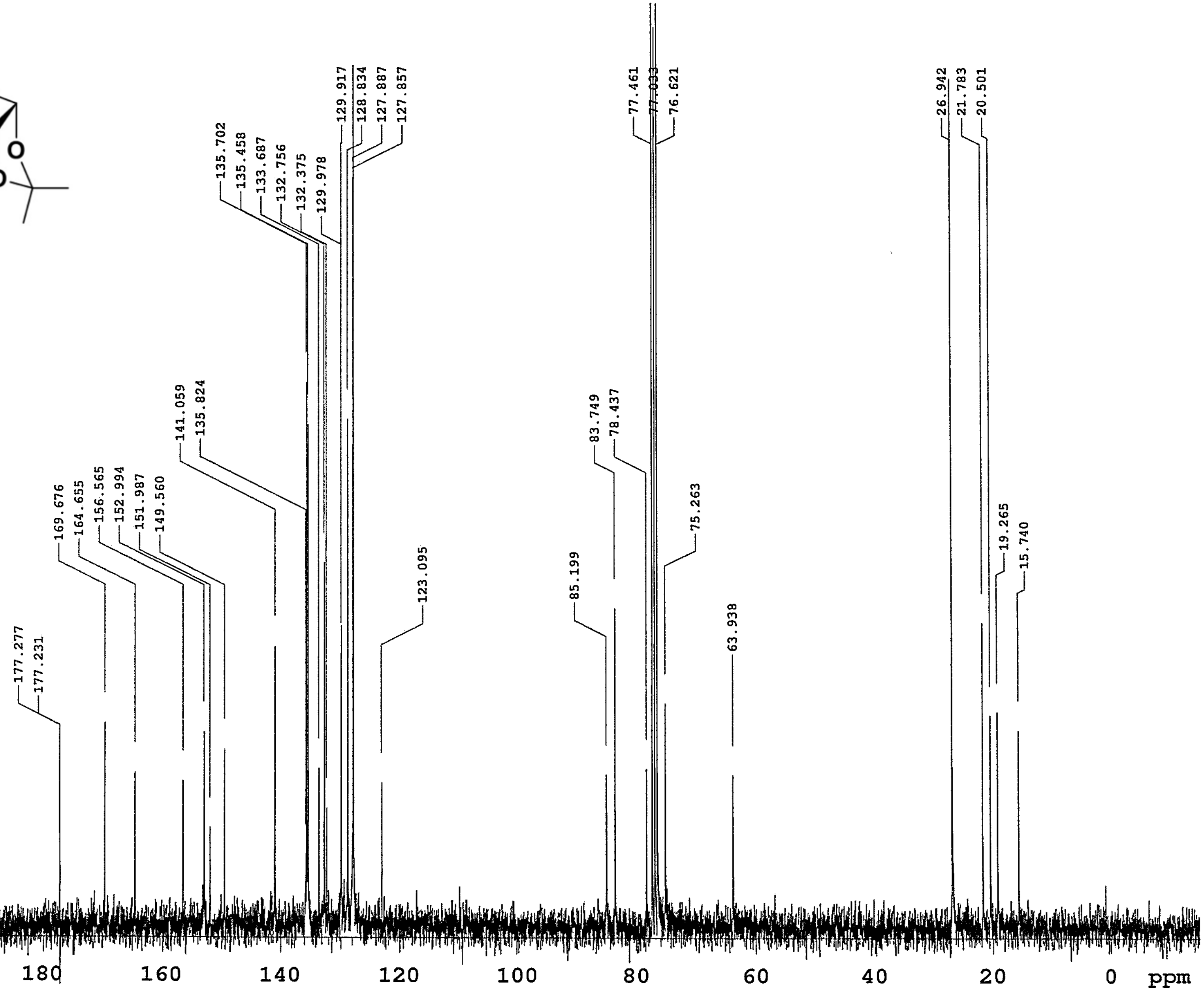
177.277
177.231
169.676
164.655
156.565
152.994
151.987
149.560
141.059
135.824
135.702
135.458
133.687
132.756
132.375
129.978
129.917
128.834
127.887
127.857
123.095
85.199
83.749
78.437
77.461
77.033
76.621
75.263
63.938
26.942
21.783
20.501
19.265
15.740
220 200 180 160 140 120 100 80 60 40 20 0 ppm

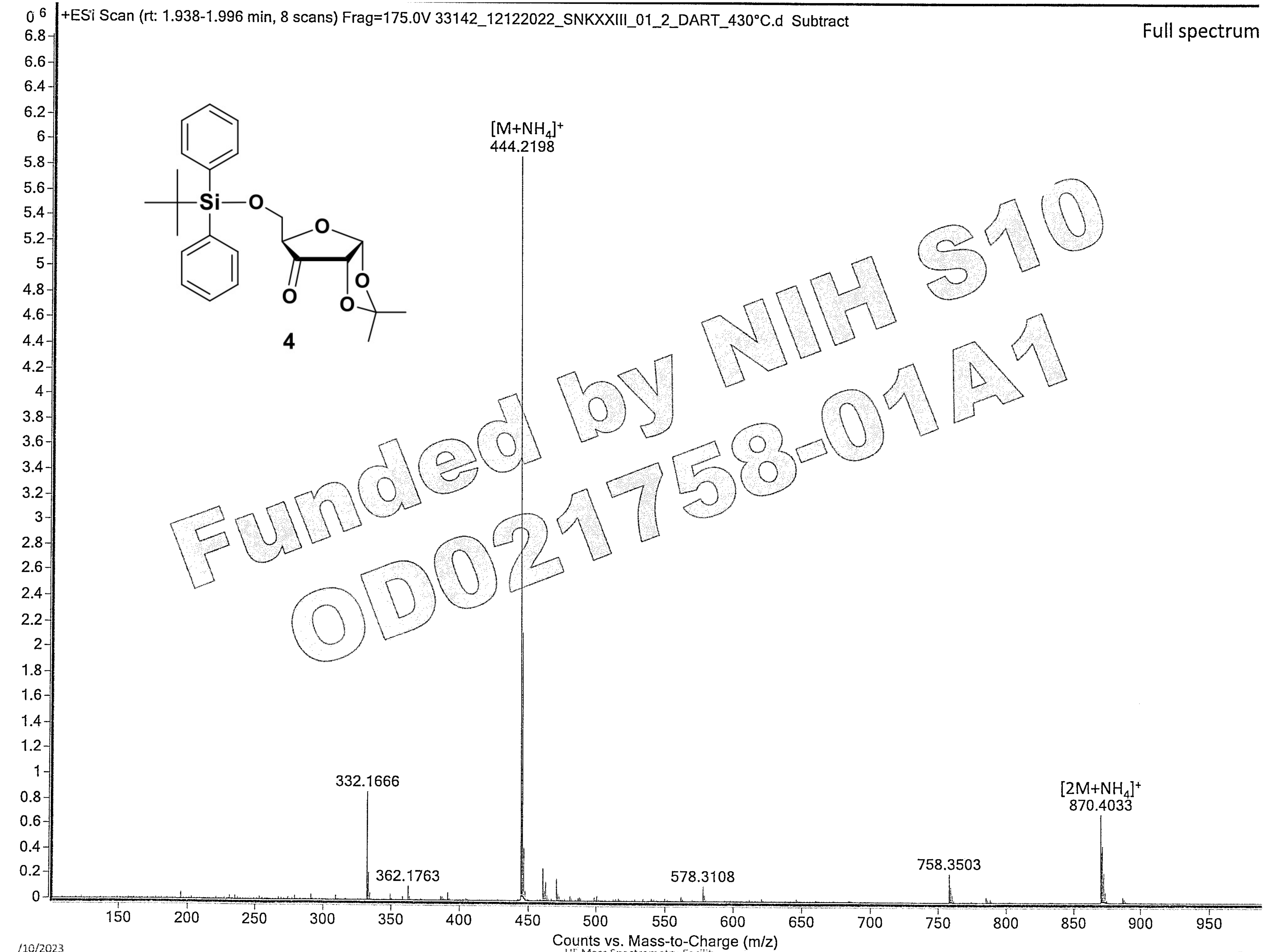

Funded by NIH S10 OD021758-01A1

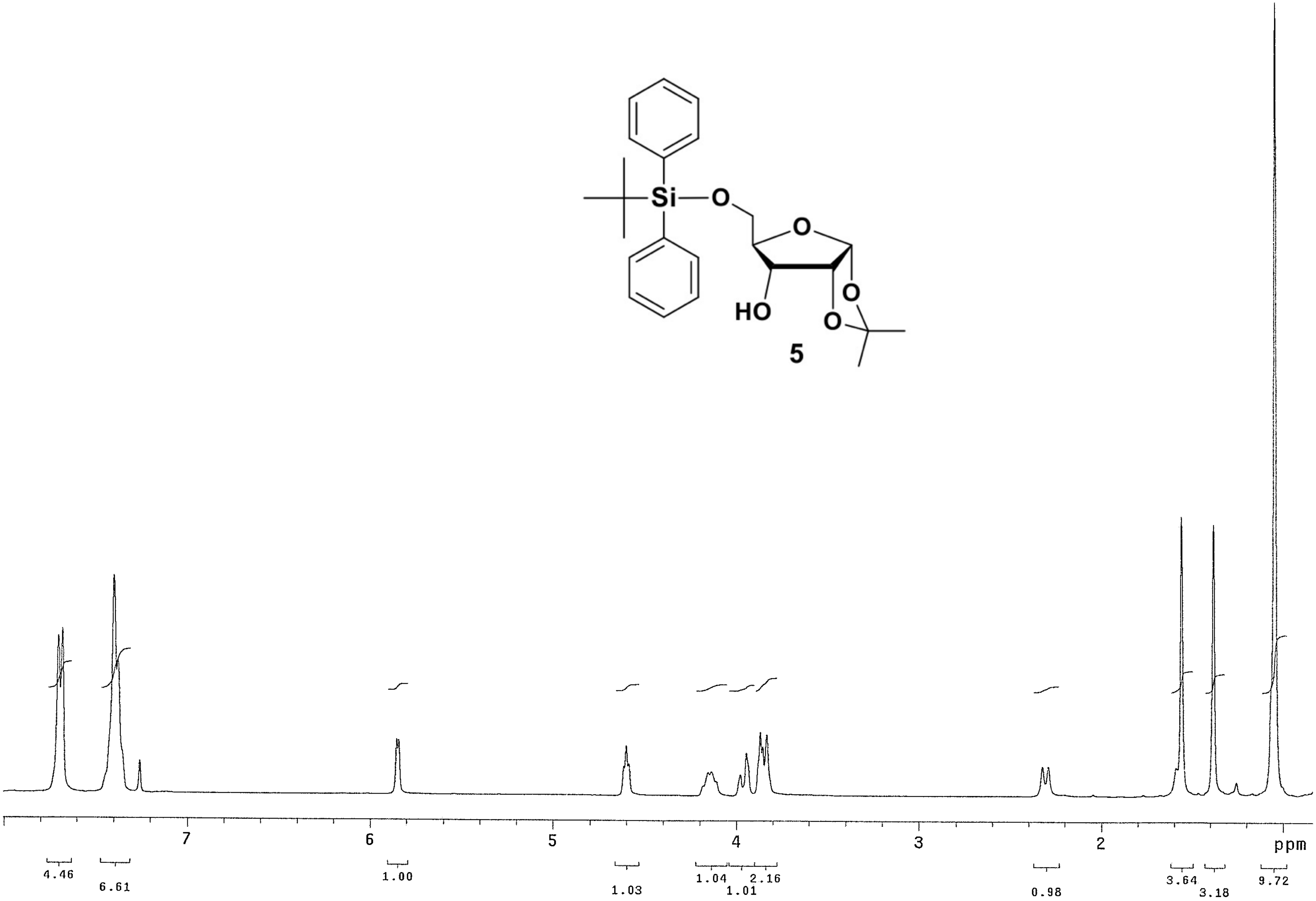
5
7
6
5
4
3
2
ppm
4.46
6.61
1.00
1.03
1.04
1.01
2.16
0.98
3.64
3.18
9.72

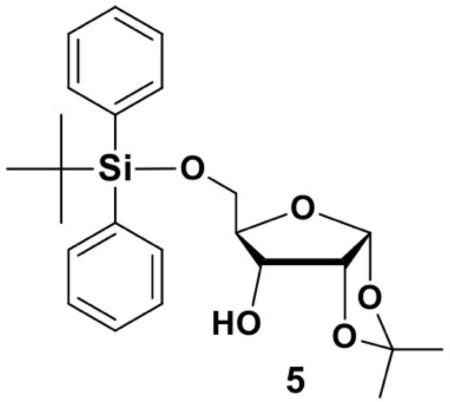

Si
O
O
HO
O
O


5

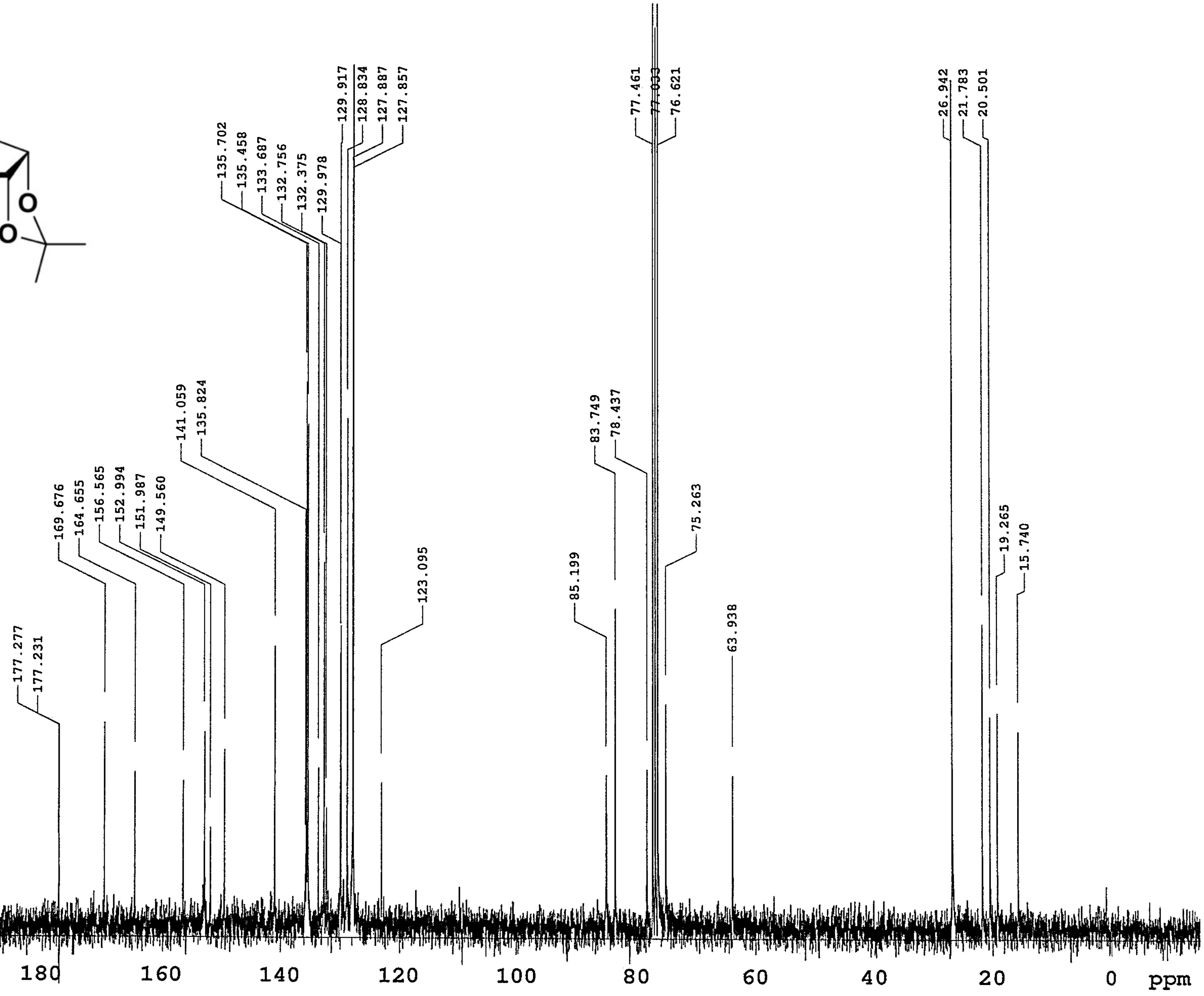

177.277
177.231
169.676
164.655
156.565
152.994
151.987
149.560
141.059
135.824
135.702
135.458
133.687
132.756
132.375
129.978
129.917
128.834
127.887
127.857
123.095
85.199
83.749
78.437
77.461
77.033
76.621
75.263
63.938
26.942
21.783
20.501
19.265
15.740
220
200
180
160
140
120
100
80
60
40
20
0
ppm

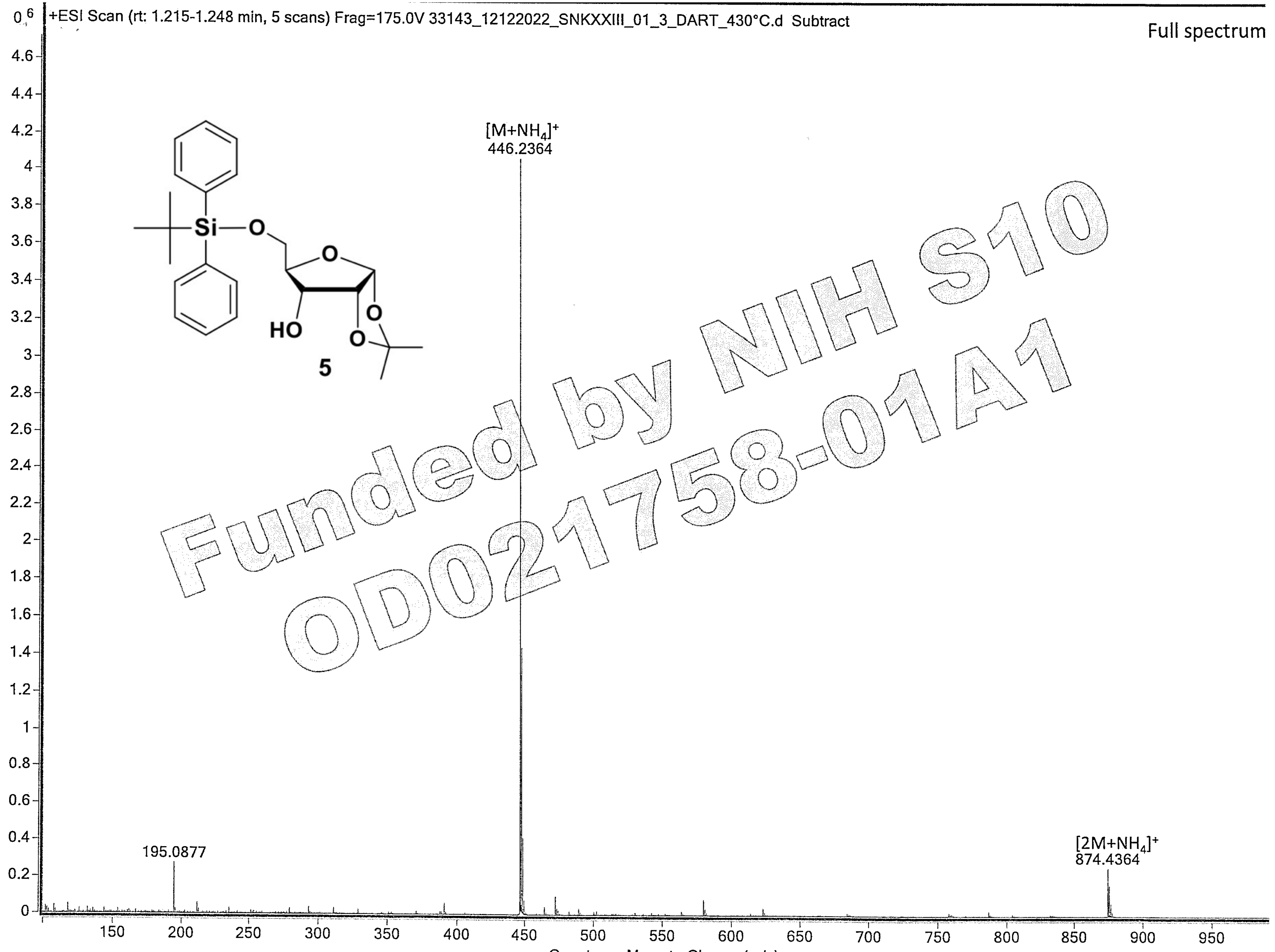

+ESI Scan (rt: 1.215-1.248 min, 5 scans) Frag=175.0V 33143_12122022_SNKXXIII_01_3_DART_430°C.d Subtract
Full spectrum
$[M+NH_4]^+$
446.2364
195.0877
$[2M+NH_4]^+$
874.4364
HO
5
Funded by NIH S10 OD021758-01A1
Counts vs. Mass-to-Charge (m/z)

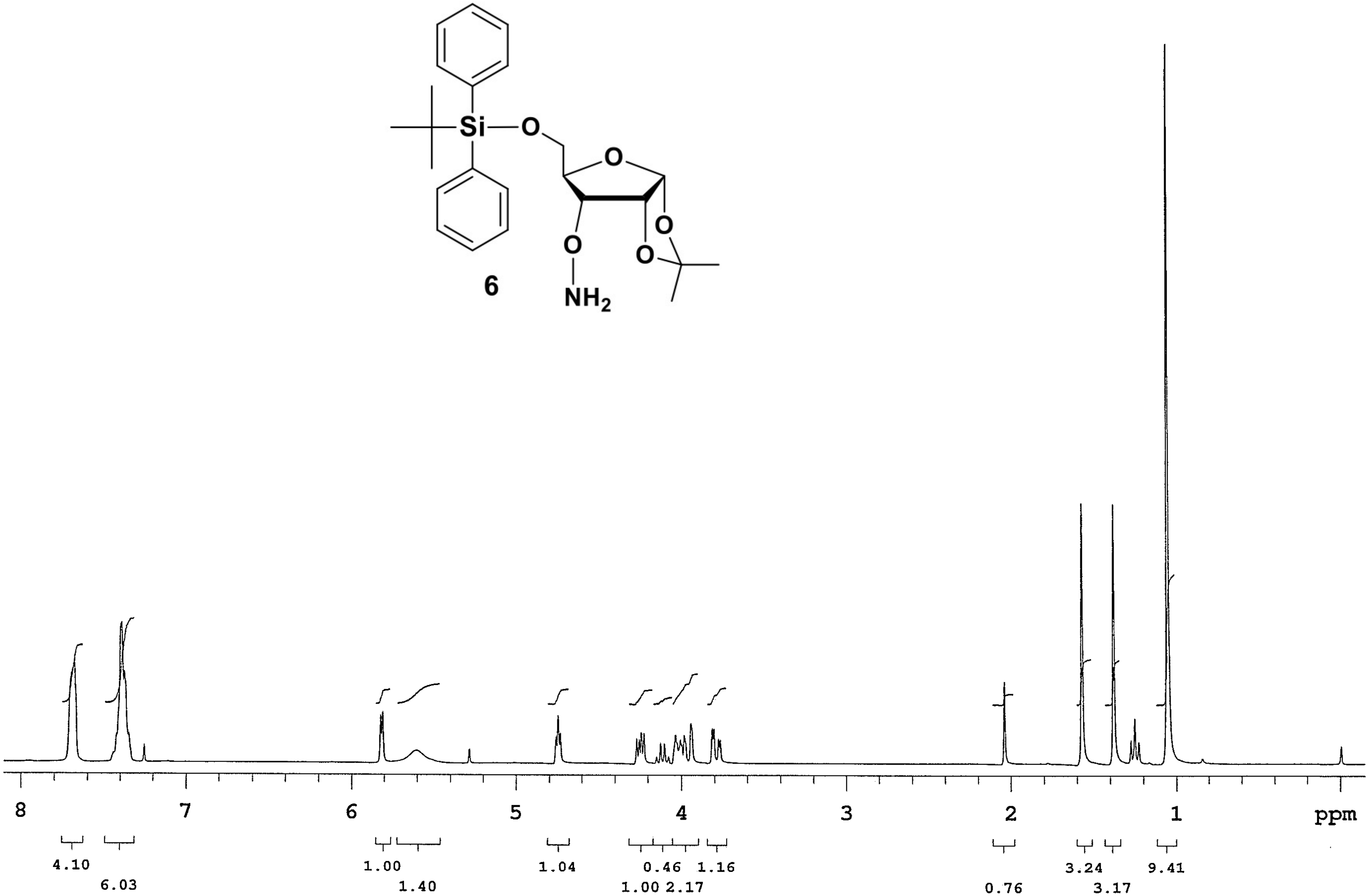
Si
O
O
O
O
O
NH2
6
8
7
6
5
4
3
2
1
ppm
4.10
6.03
1.00
1.40
1.04
0.46
1.16
1.00 2.17
0.76
3.24
3.17
9.41

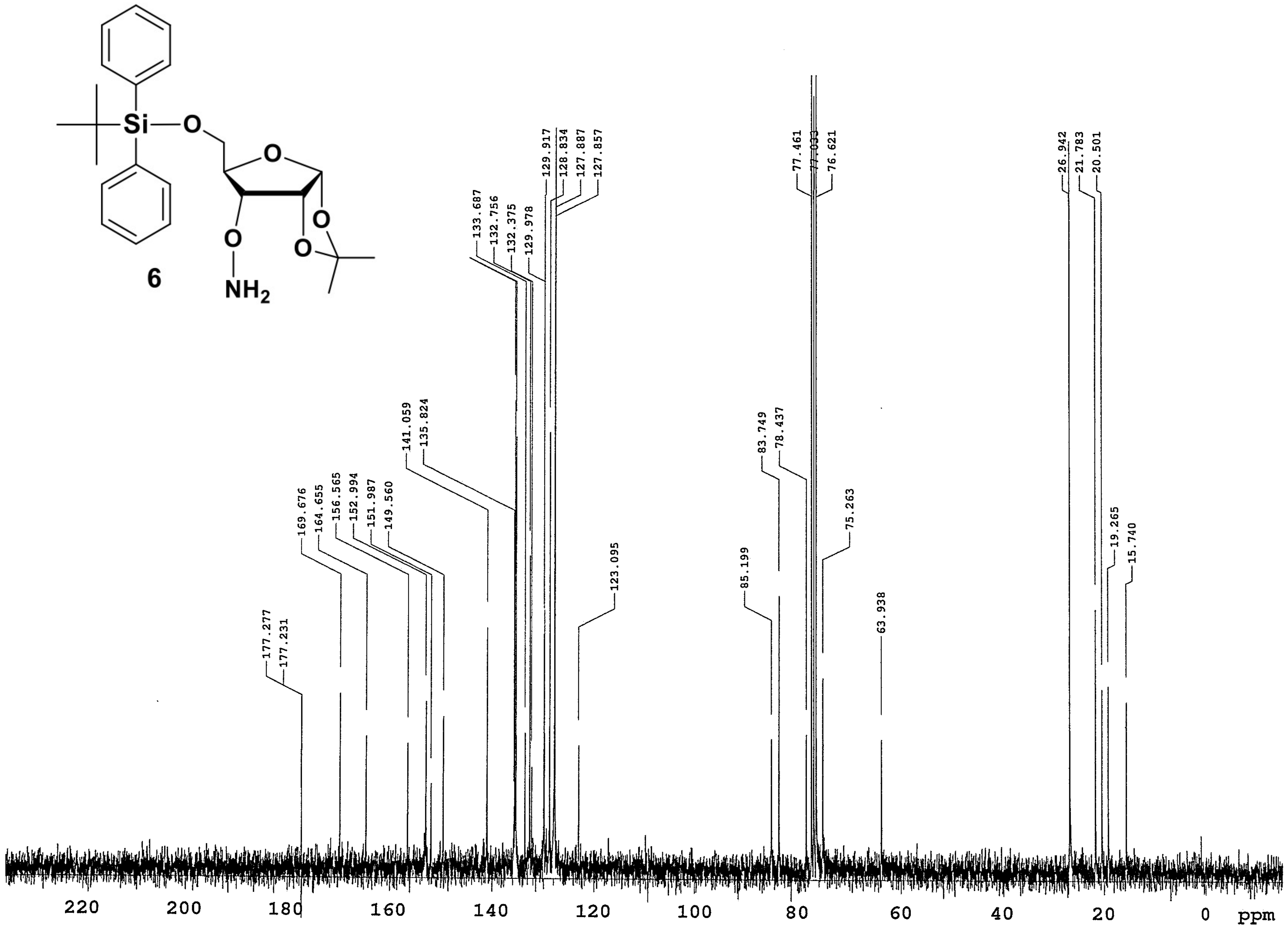
Si
O
O
O
O
O
NH2
6
177.277
177.231
169.676
164.655
156.565
152.994
151.987
149.560
141.059
135.824
133.687
132.756
132.375
129.978
129.917
128.834
127.887
127.857
123.095
85.199
83.749
78.437
77.461
77.033
76.621
75.263
63.938
26.942
21.783
20.501
19.265
15.740
220
200
180
160
140
120
100
80
60
40
20
0
ppm

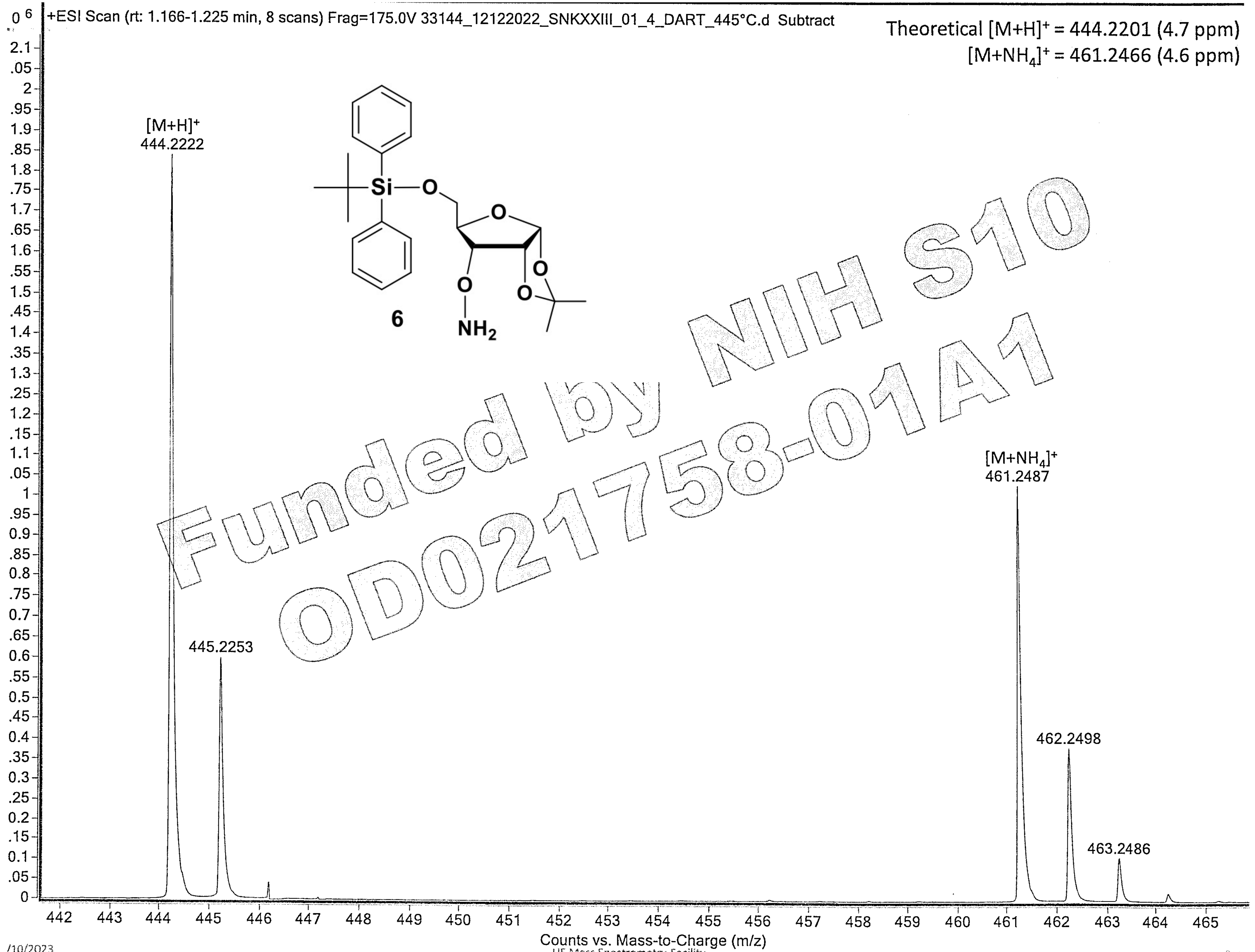

Funded by NIH S10 OD021758-01A1

/10/2023

UF Mass Spectrometry Facility

Page

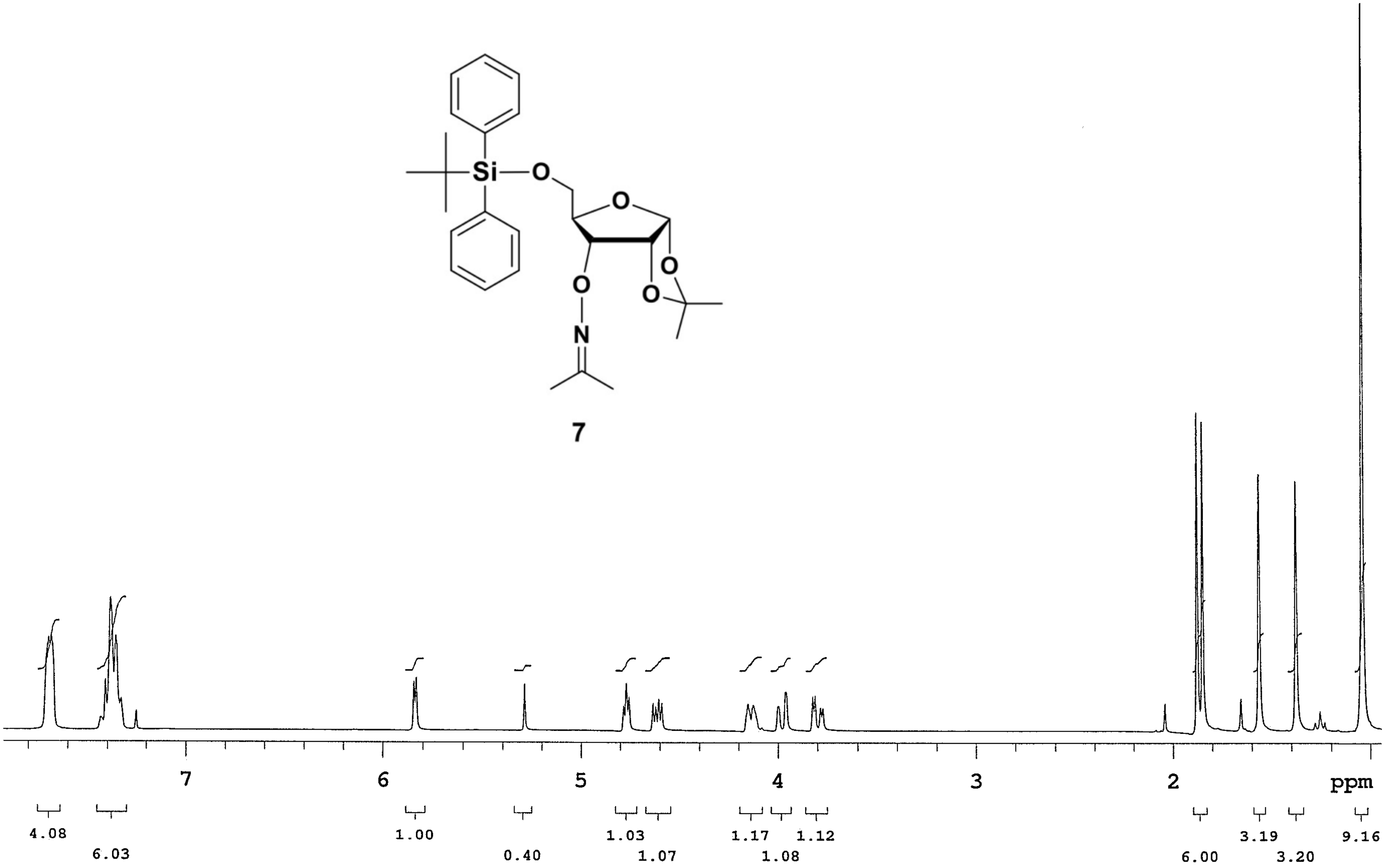

Si
O
O
O
O
O
N
7
7
6
5
4
3
2
ppm
4.08
6.03
1.00
0.40
1.03
1.07
1.17
1.08
1.12
6.00
3.19
3.20
9.16

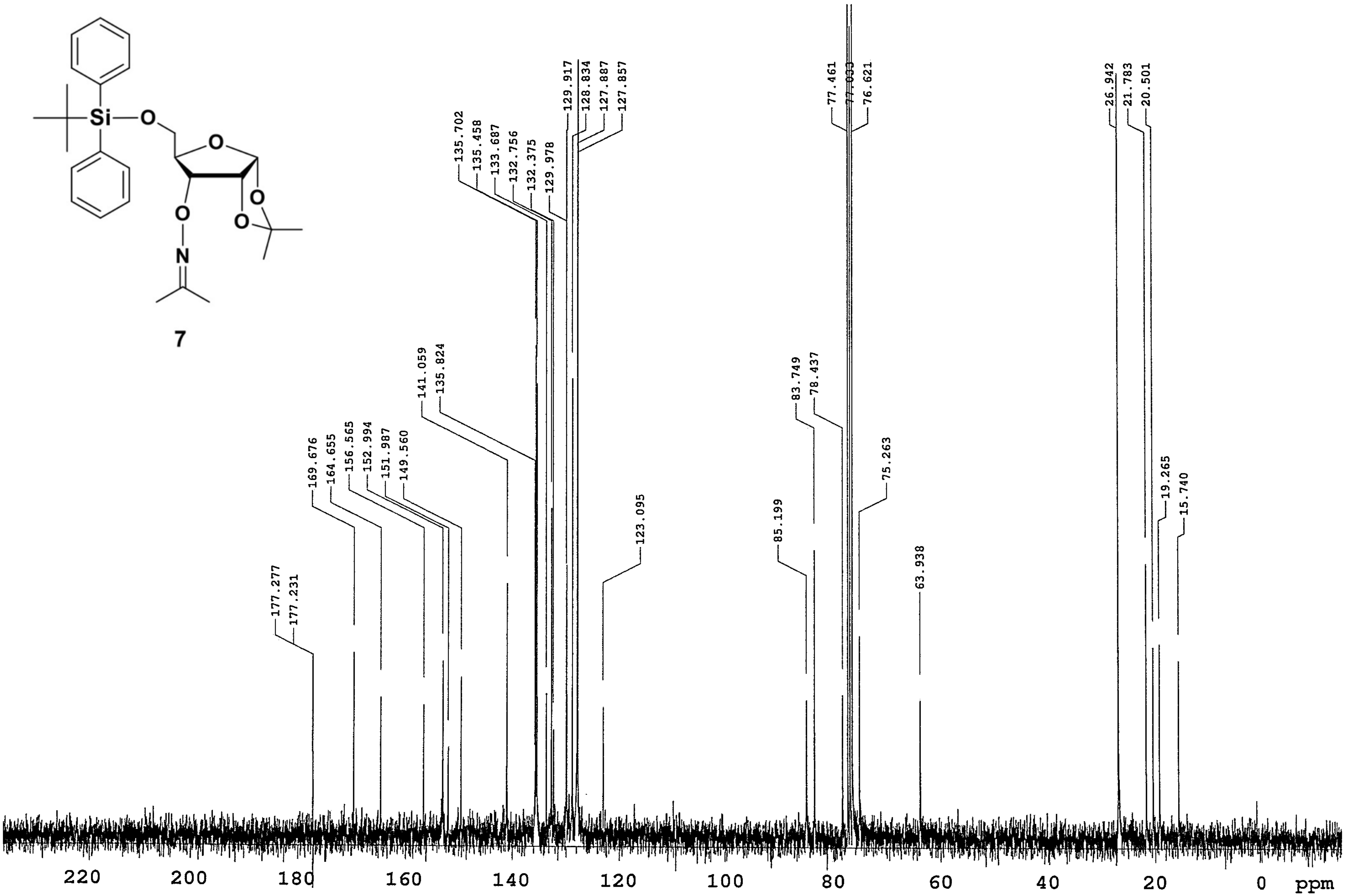
Si
O
O
O
O
O
N
177.277
177.231
169.676
164.655
156.565
152.994
151.987
149.560
141.059
135.824
135.702
135.458
133.687
132.756
132.375
129.978
129.917
128.834
127.887
127.857
123.095
85.199
83.749
78.437
77.461
77.033
76.621
75.263
63.938
26.942
21.783
20.501
19.265
15.740
220
200
180
160
140
120
100
80
60
40
20
0
ppm

**7**

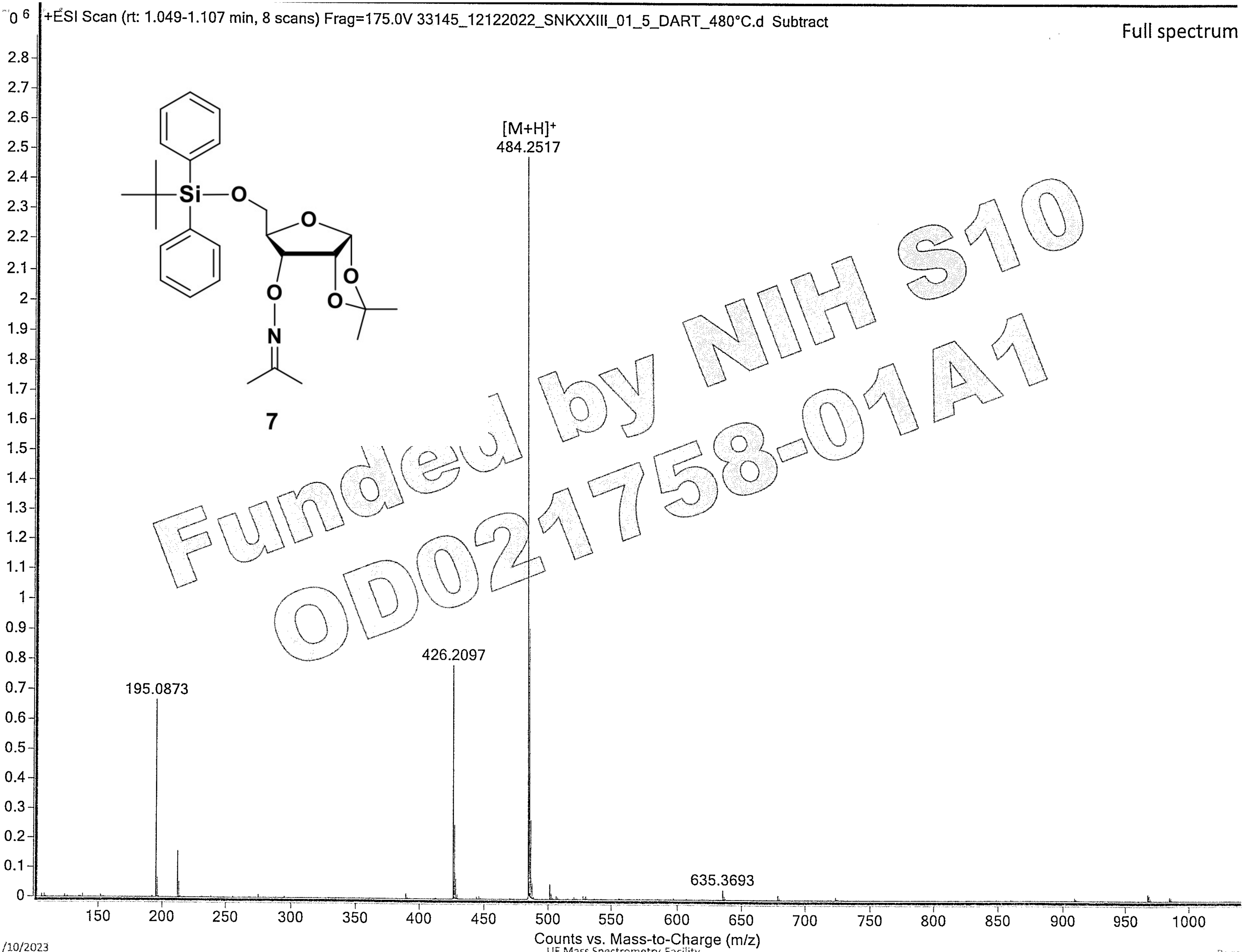

Funded by NIH S10 OD021758-01A1

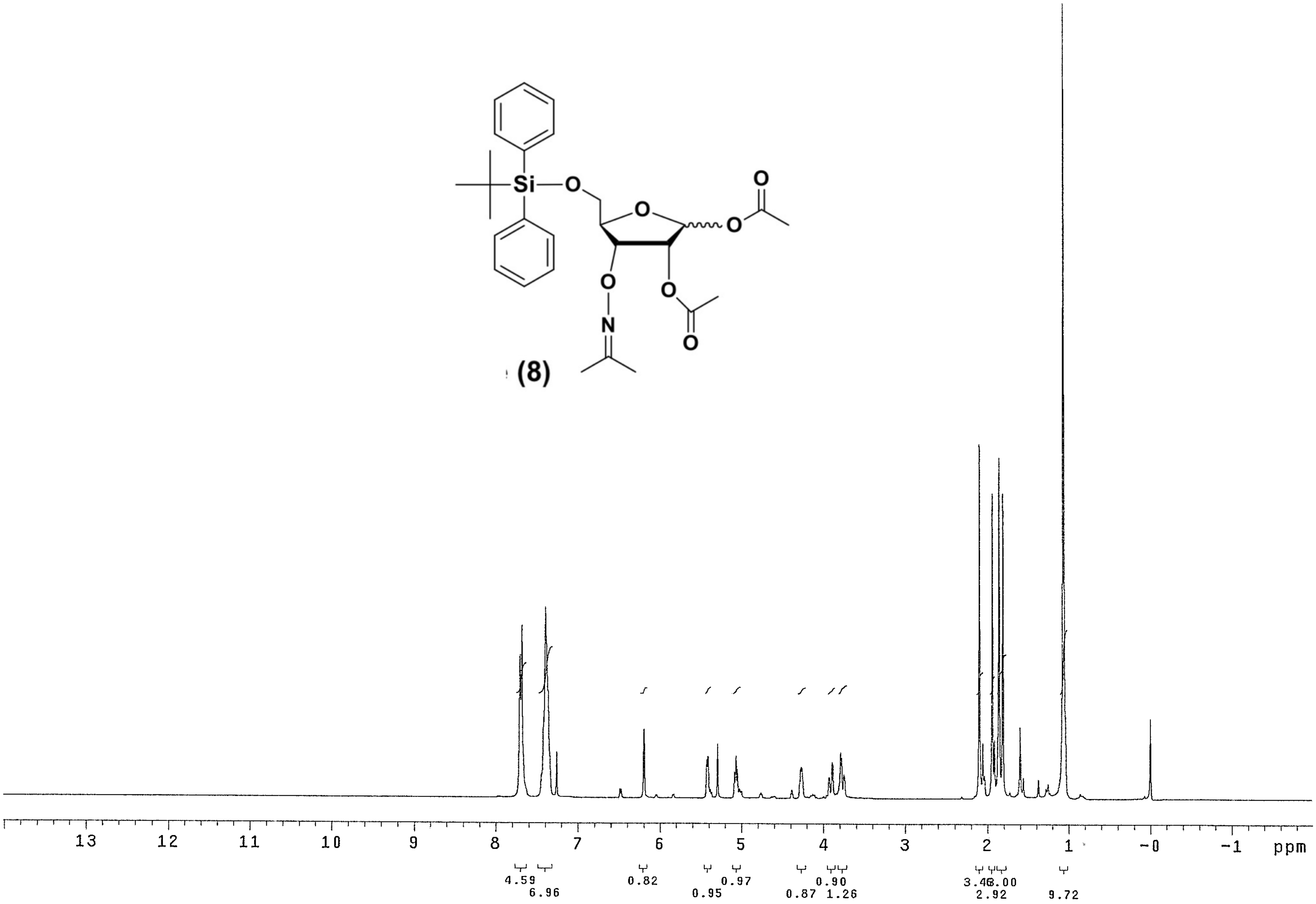
Si
O
O
O
O
O
O
O
N
(8)
13
12
11
10
9
8
7
6
5
4
3
2
1
-0
-1
ppm
4.59
6.96
0.82
0.95
0.97
0.87
0.90
1.26
3.46
3.00
2.92
9.72

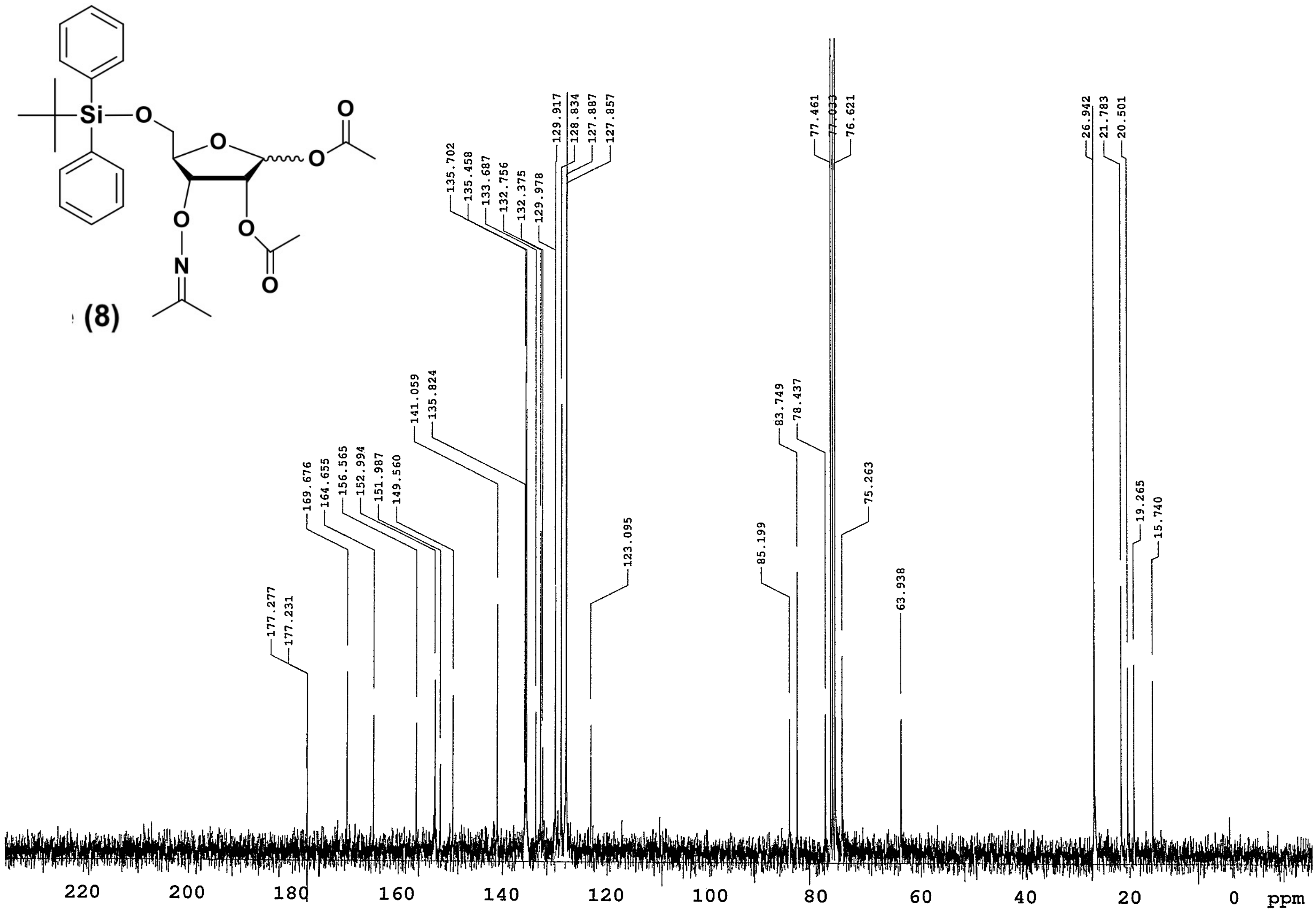
Si
O
O
O
O
O
O
N
O
177.277
177.231
169.676
164.655
156.565
152.994
151.987
149.560
141.059
135.824
135.702
135.458
133.687
132.756
132.375
129.978
129.917
128.834
127.887
127.857
123.095
85.199
83.749
78.437
77.461
77.033
76.621
75.263
63.938
26.942
21.783
20.501
19.265
15.740
220
200
180
160
140
120
100
80
60
40
20
0
ppm

(8)

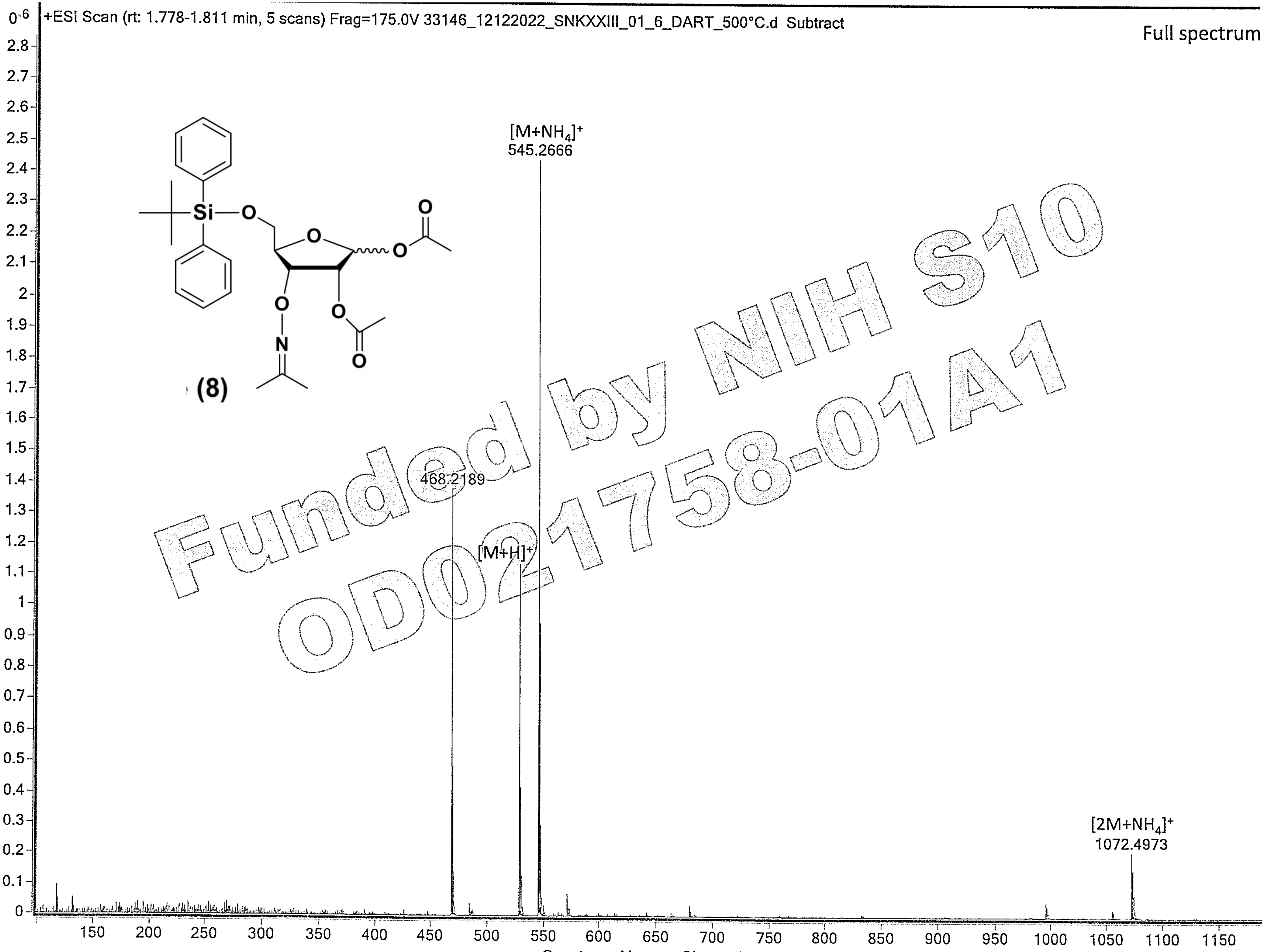


/10/2023

UF Mass Spectrometry Facility

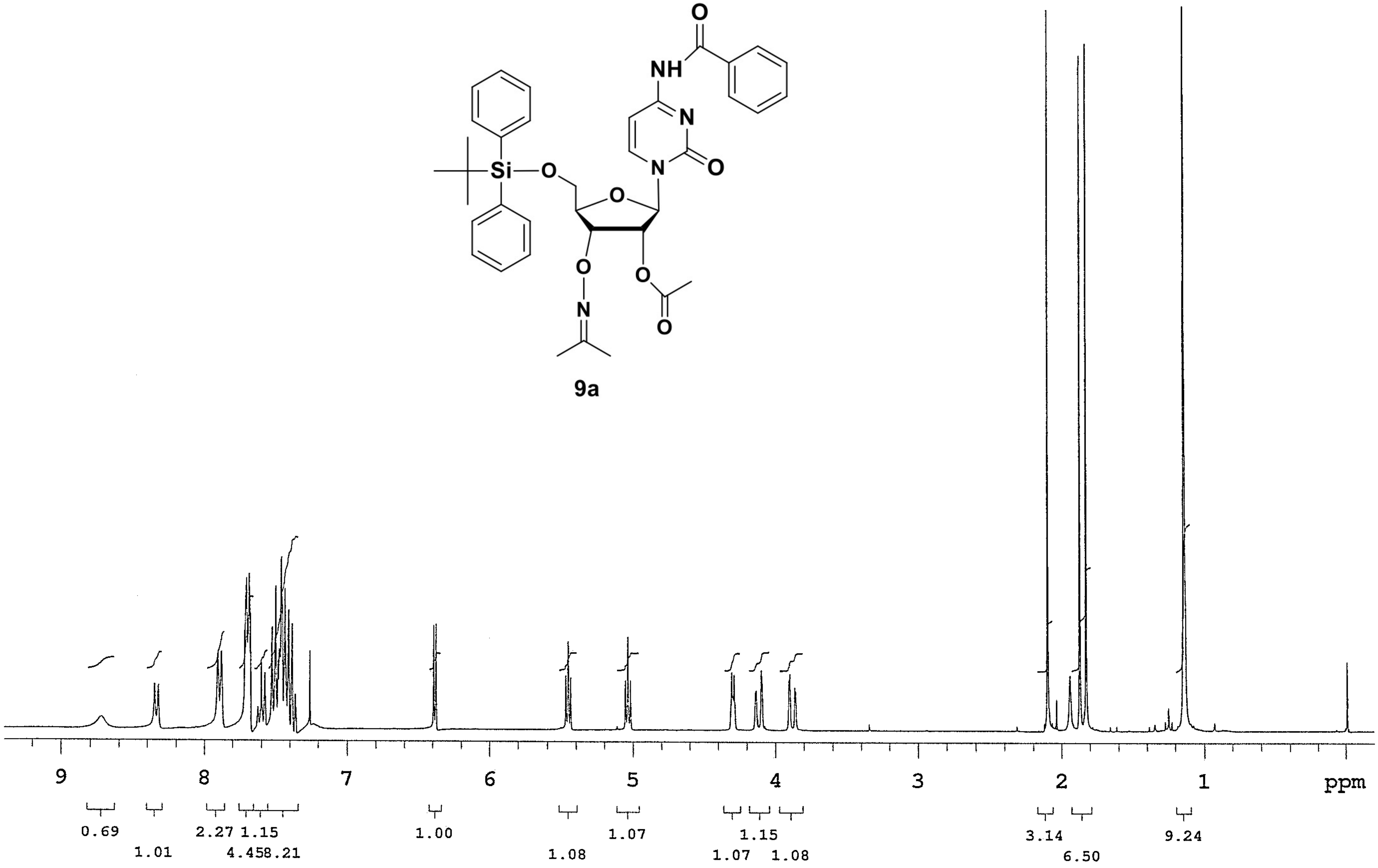

O
NH
N
O
Si
O
O
N
O
O
N
O
9a
9
8
7
6
5
4
3
2
1
ppm
0.69
1.01
2.27
1.15
4.45
8.21
1.00
1.08
1.07
1.15
1.07
1.08
3.14
6.50
9.24

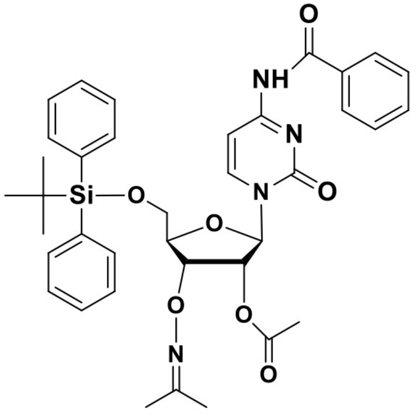

**9a**

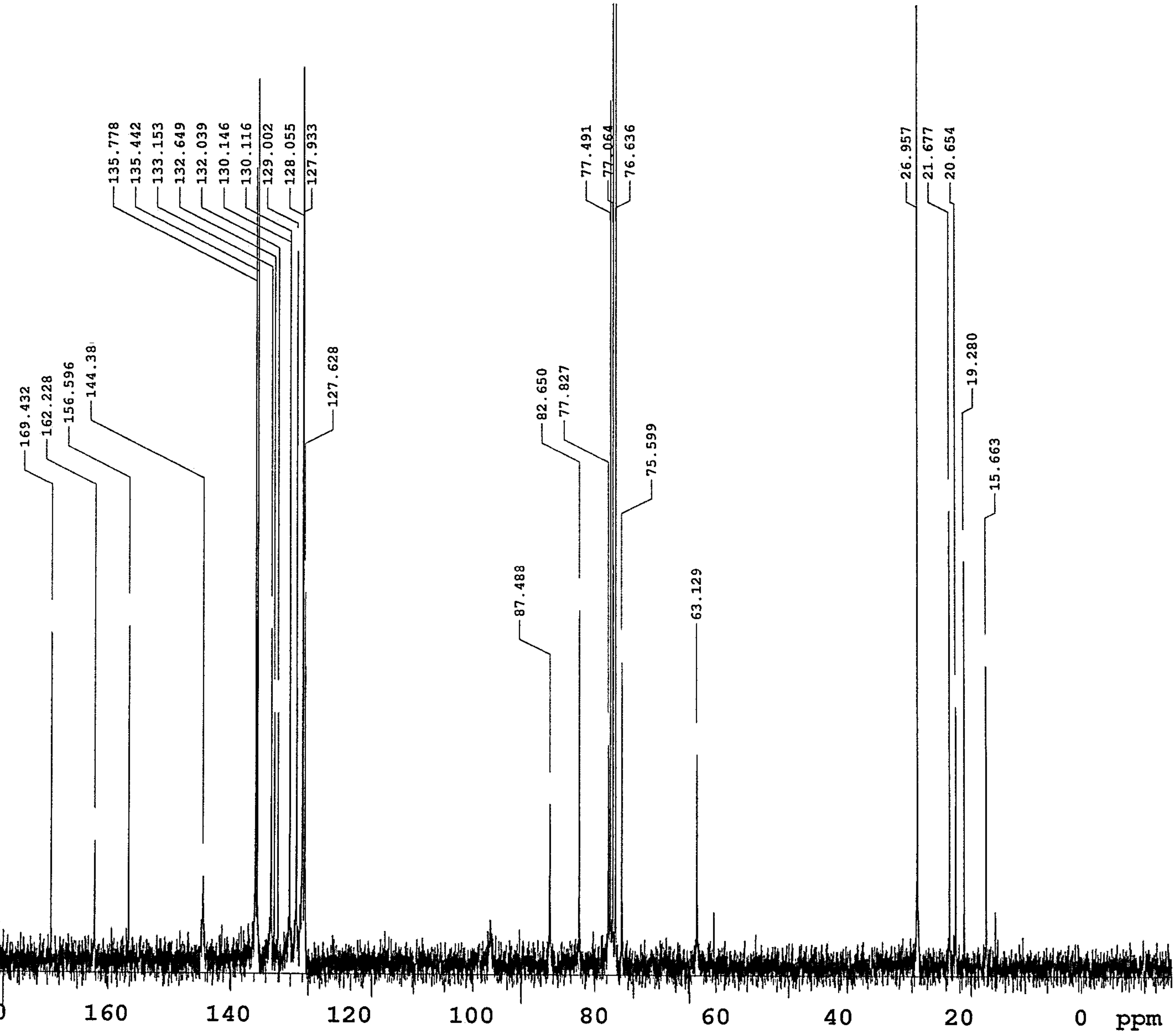

169.432
162.228
156.596
144.38
135.778
135.442
133.153
132.649
132.039
130.146
130.116
129.002
128.055
127.933
127.628
87.488
82.650
77.827
77.491
77.064
76.636
75.599
63.129
26.957
21.677
20.654
19.280
15.663
220
200
180
160
140
120
100
80
60
40
20
0
ppm

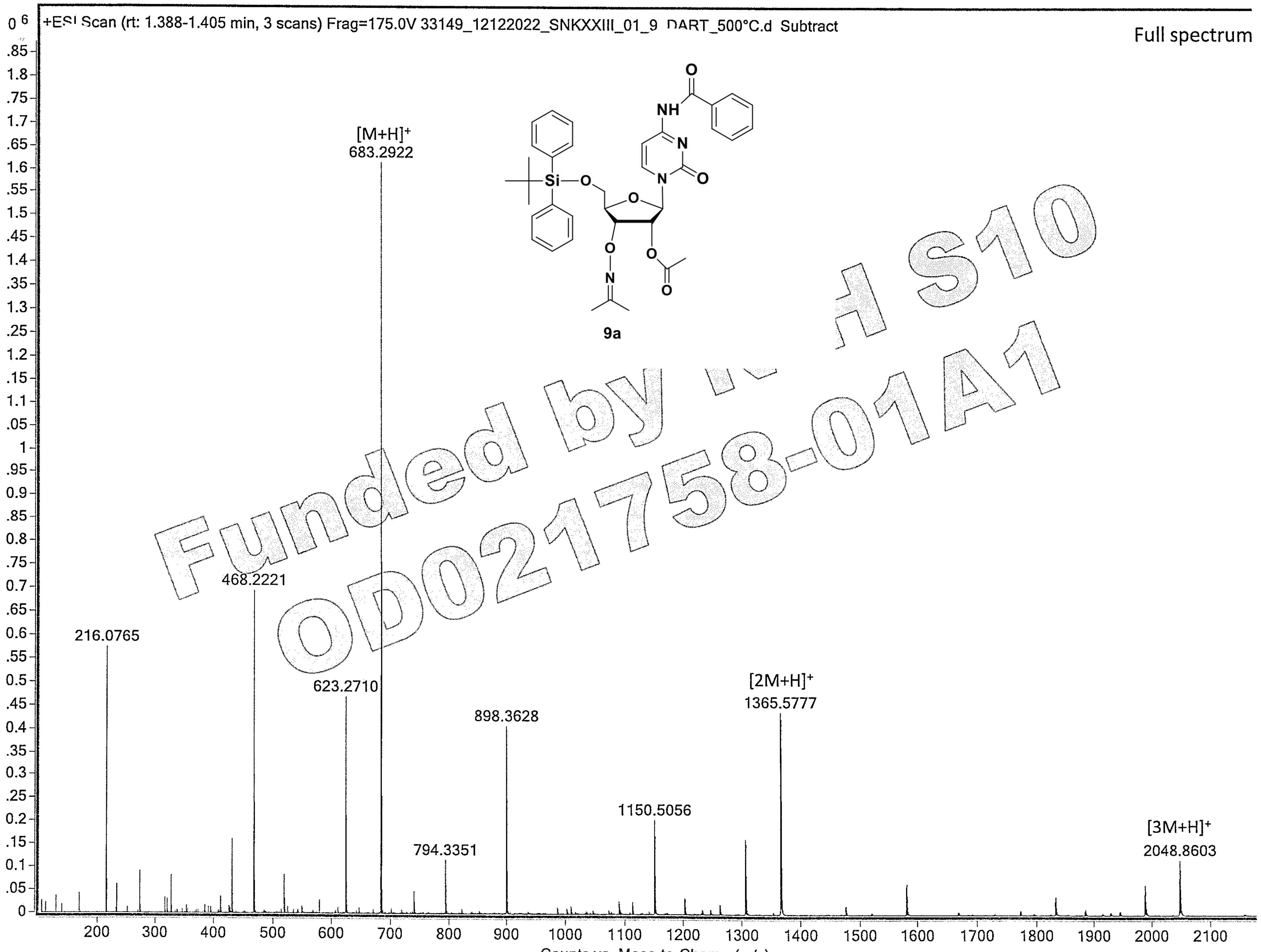

Funded by NIH S10 OD021758-01A1

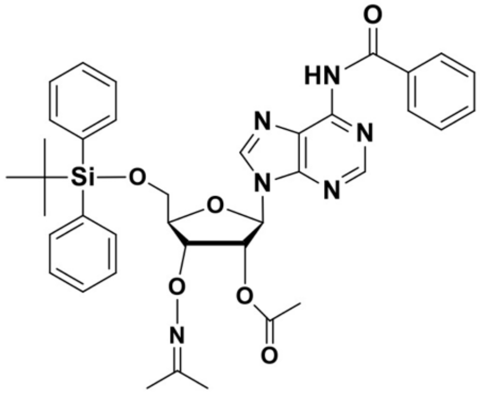
O
HN
N
N
Si
O
O
N
N
O
O
N
O

9b

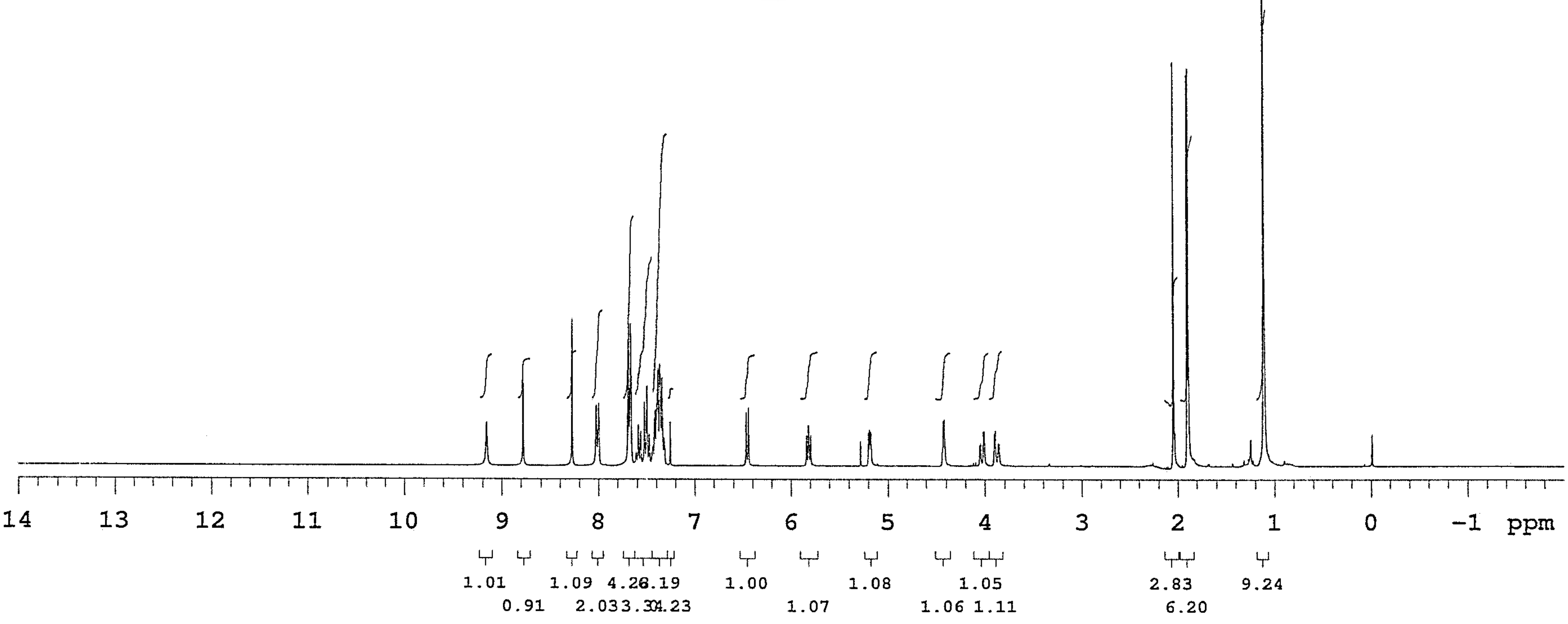
14
13
12
11
10
9
8
7
6
5
4
3
2
1
0
-1
ppm
1.01
0.91
1.09
1.00
1.07
1.08
1.06
1.05
1.11
2.83
6.20
9.24

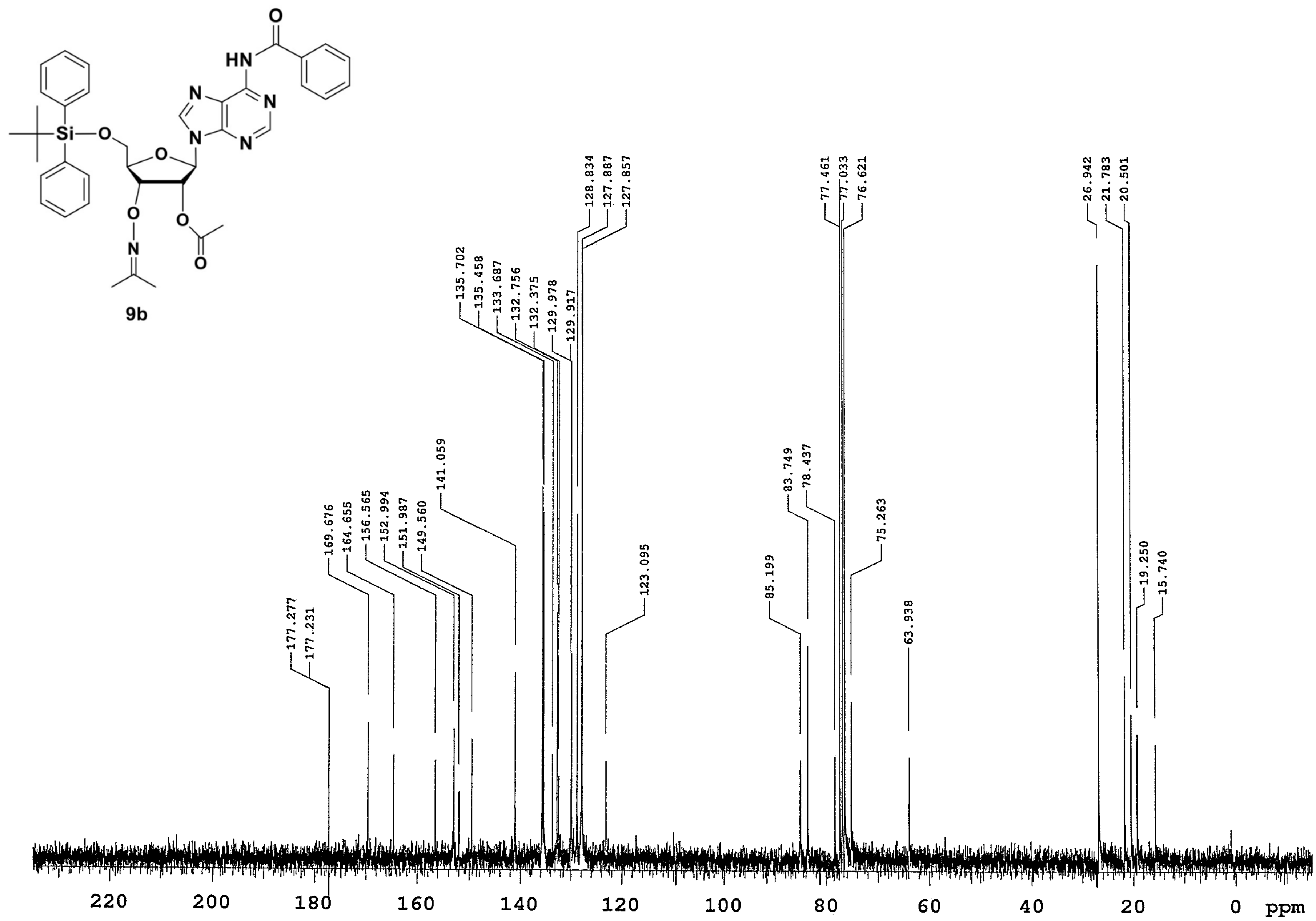

Si
O
N
HN
N
N
N
9b
177.277
177.231
169.676
164.655
156.565
152.994
151.987
149.560
141.059
135.702
135.458
133.687
132.756
132.375
129.978
129.917
128.834
127.887
127.857
123.095
85.199
83.749
78.437
77.461
77.033
76.621
75.263
63.938
26.942
21.783
20.501
19.250
15.740
220
200
180
160
140
120
100
80
60
40
20
0
ppm

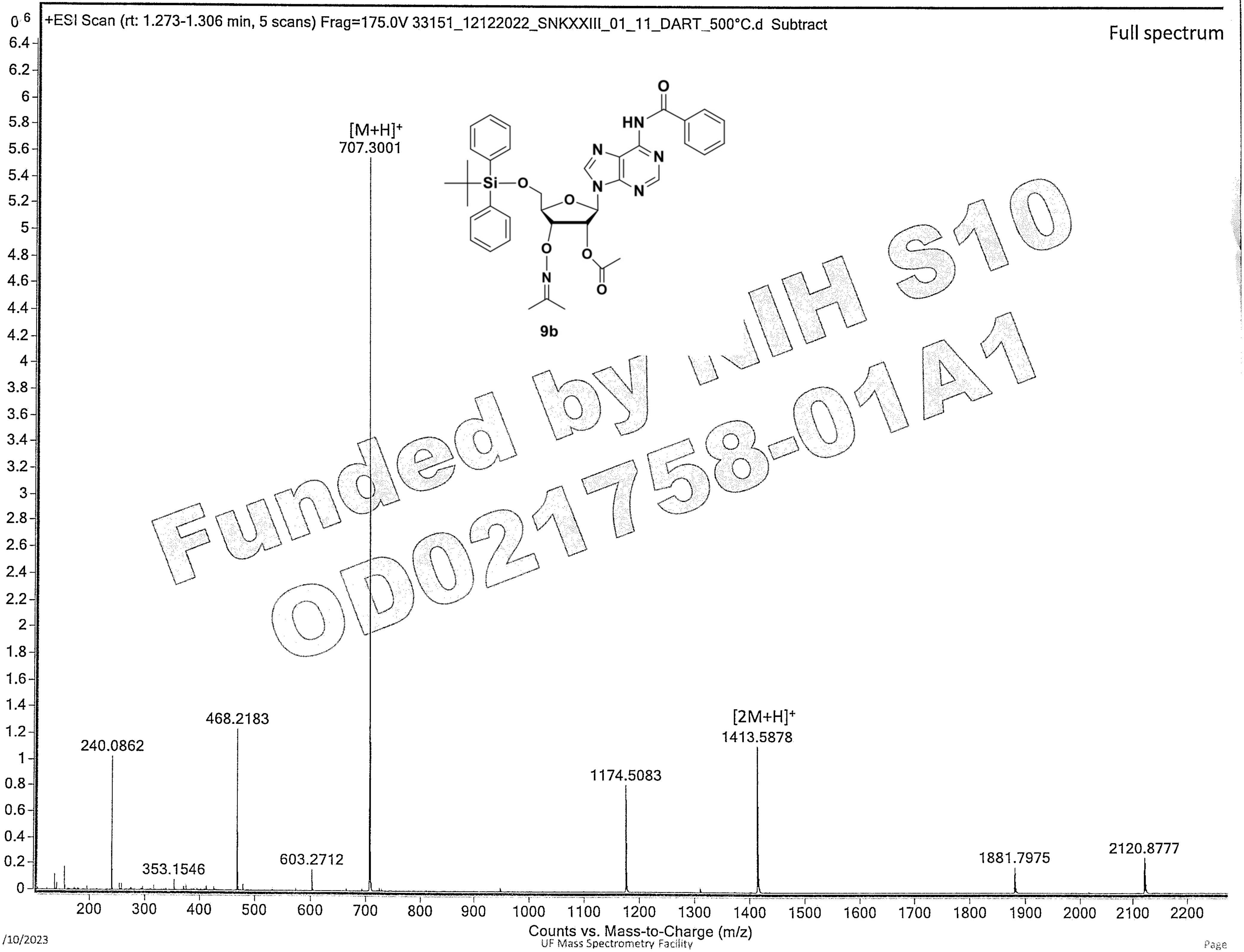

Funded by NIH S10 OD021758-01A1

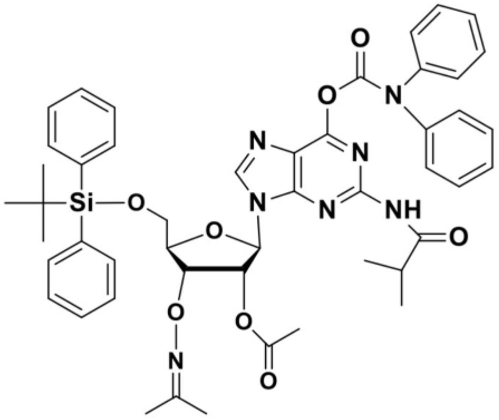
O
N
N
N
N
N
O
NH
O
Si
O
O
O
O
N
O

9c

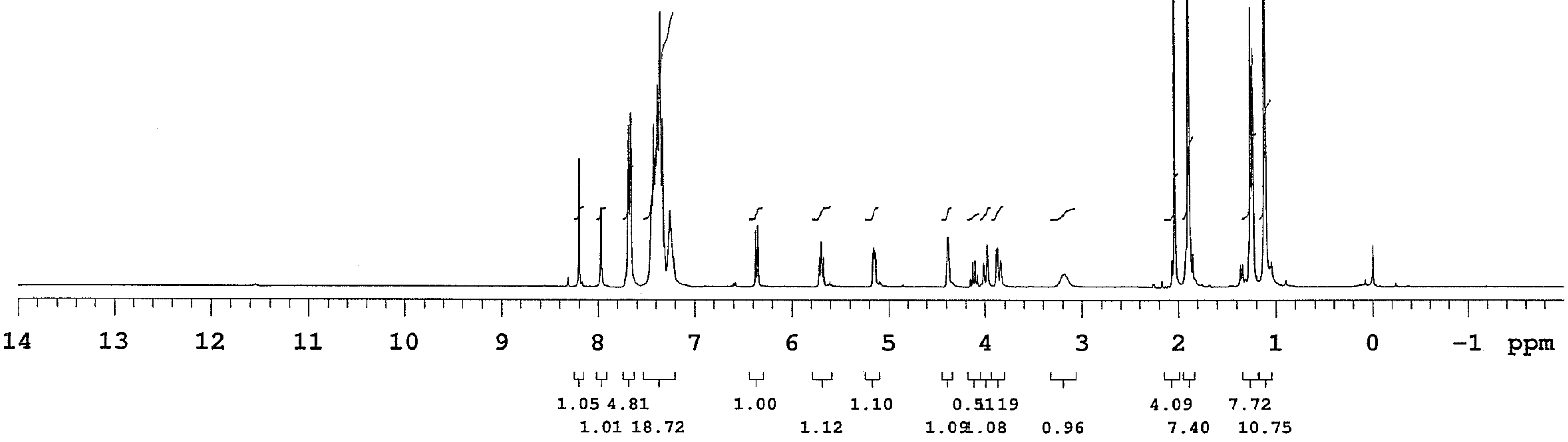
14 13 12 11 10 9 8 7 6 5 4 3 2 1 0 -1 ppm
1.05 4.81 1.00 1.10 0.51 1.19 4.09 7.72
1.01 18.72 1.12 1.09 1.08 0.96 7.40 10.75

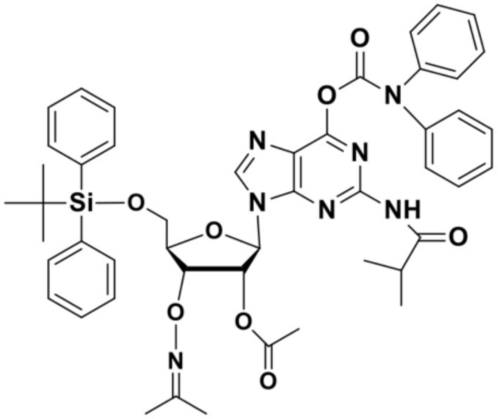

9c

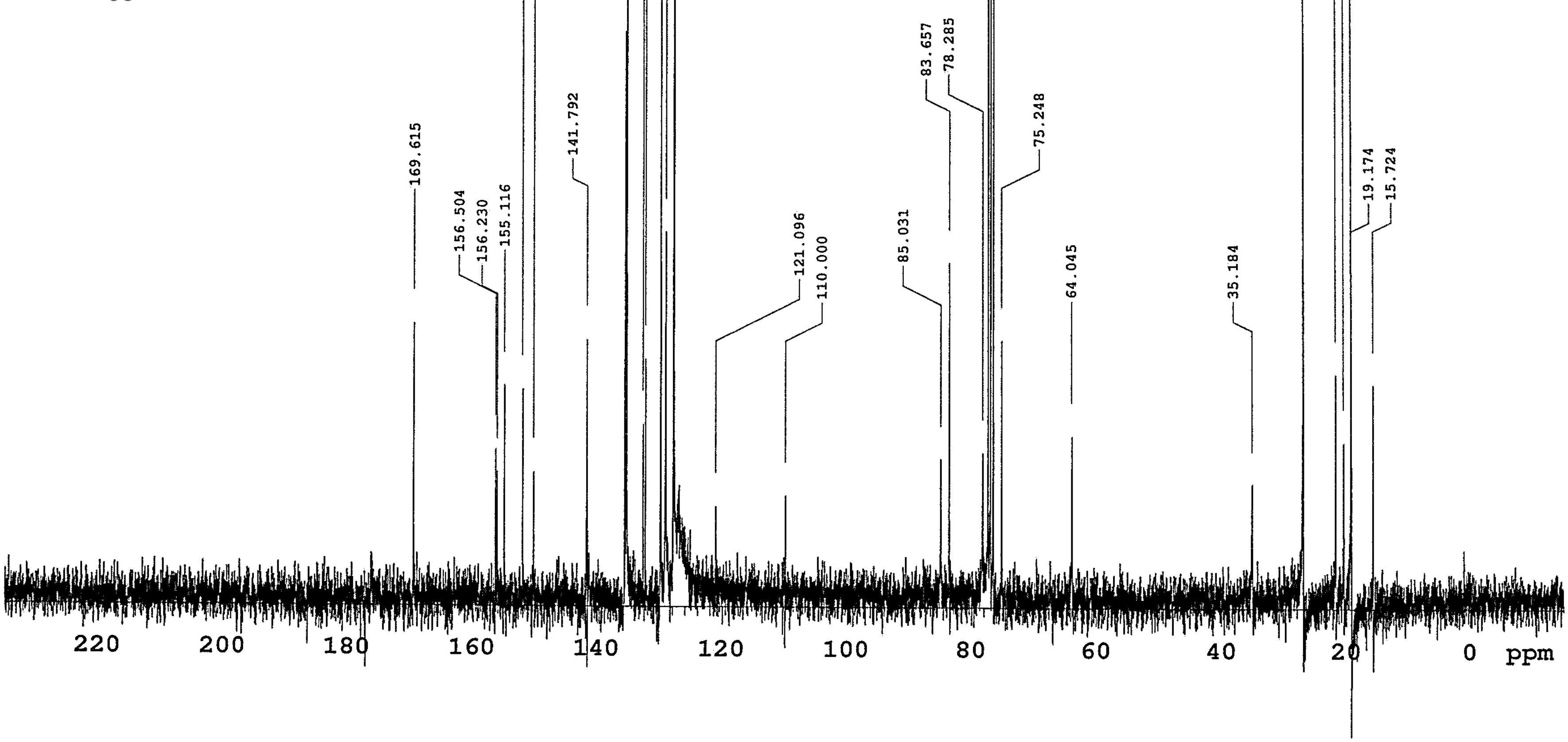

152.170
150.399
135.717
135.458
132.726
132.390
129.963
129.917
129.154
127.918
127.826
77.461
76.621
26.957
21.783
20.486
19.250
19.204
169.615
156.504
156.230
155.116
141.792
121.096
110.000
85.031
83.657
78.285
75.248
64.045
35.184
19.174
15.724
220
200
180
160
140
120
100
80
60
40
20
0
ppm

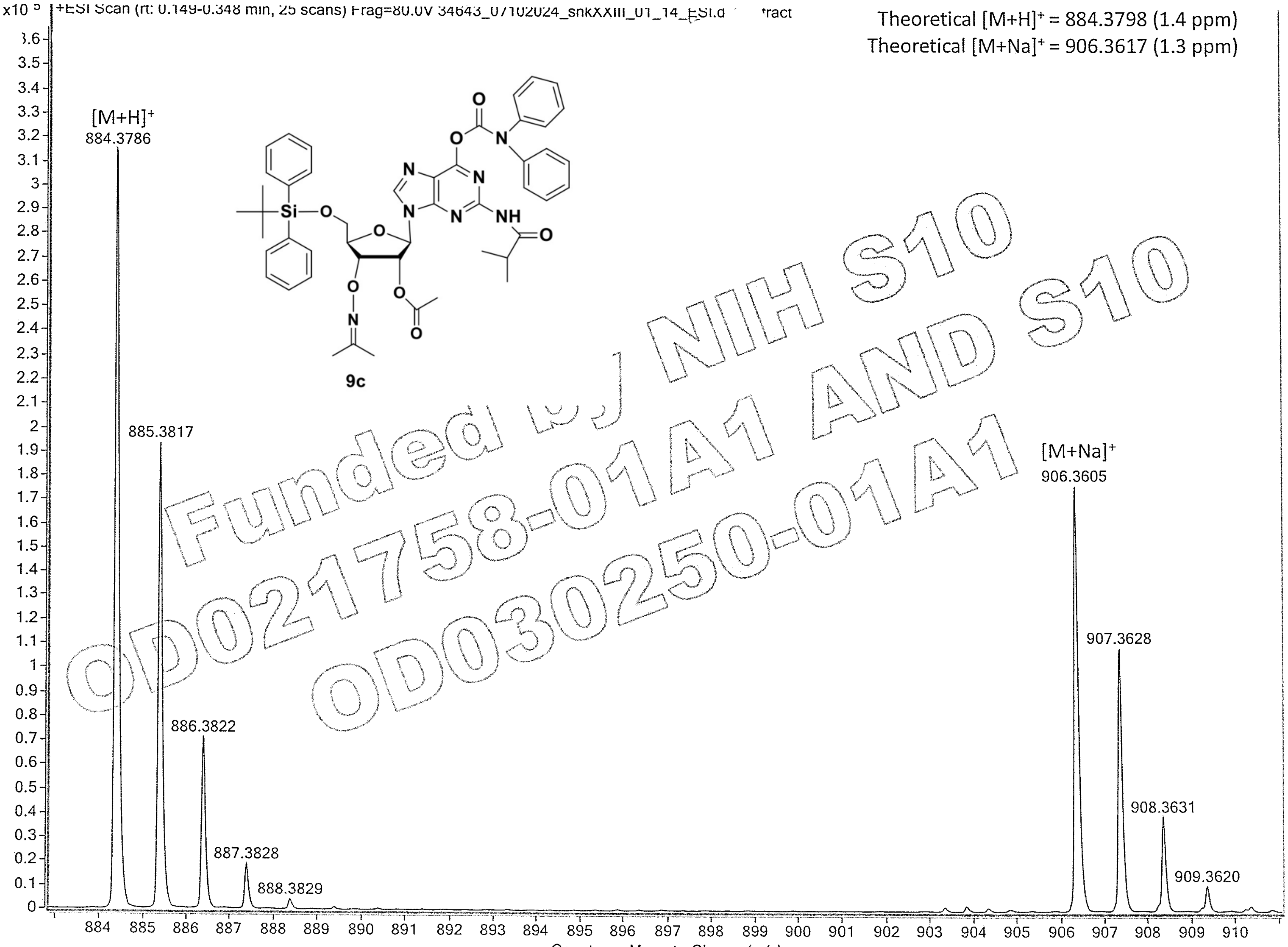

Funded by NIH S10 OD021758-01A1 AND S10 OD030250-01A1

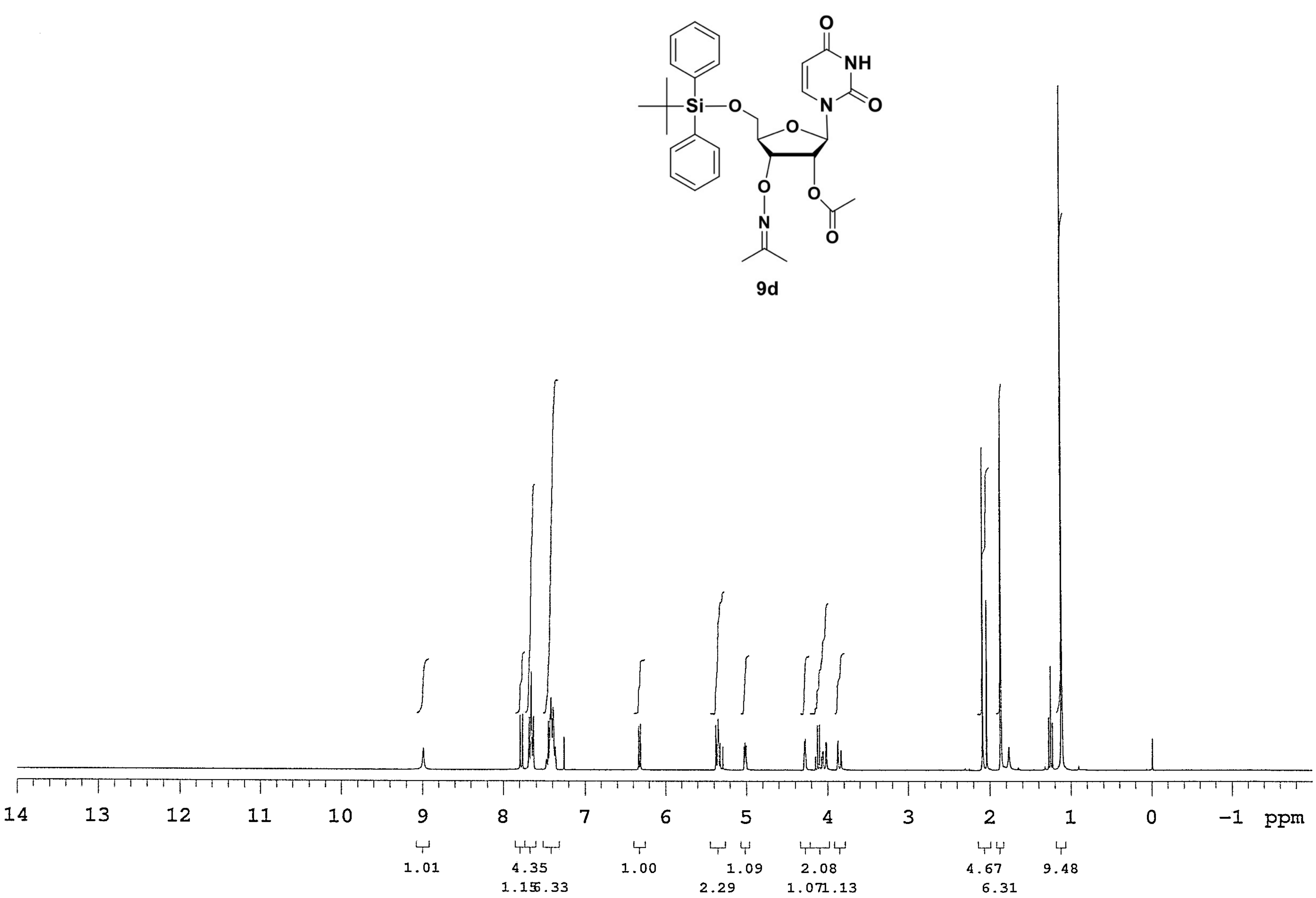

O
NH
O
N
O
Si
O
O
O
N
O
O
9d
14
13
12
11
10
9
8
7
6
5
4
3
2
1
0
−1
ppm
1.01
4.35
1.15
6.33
1.00
1.09
2.29
2.08
1.07
1.13
4.67
6.31
9.48

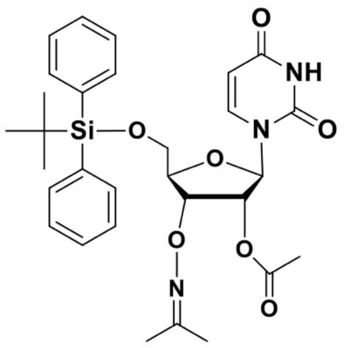
O
NH
O
N
O
Si
O
O
O
N
O
O

9d

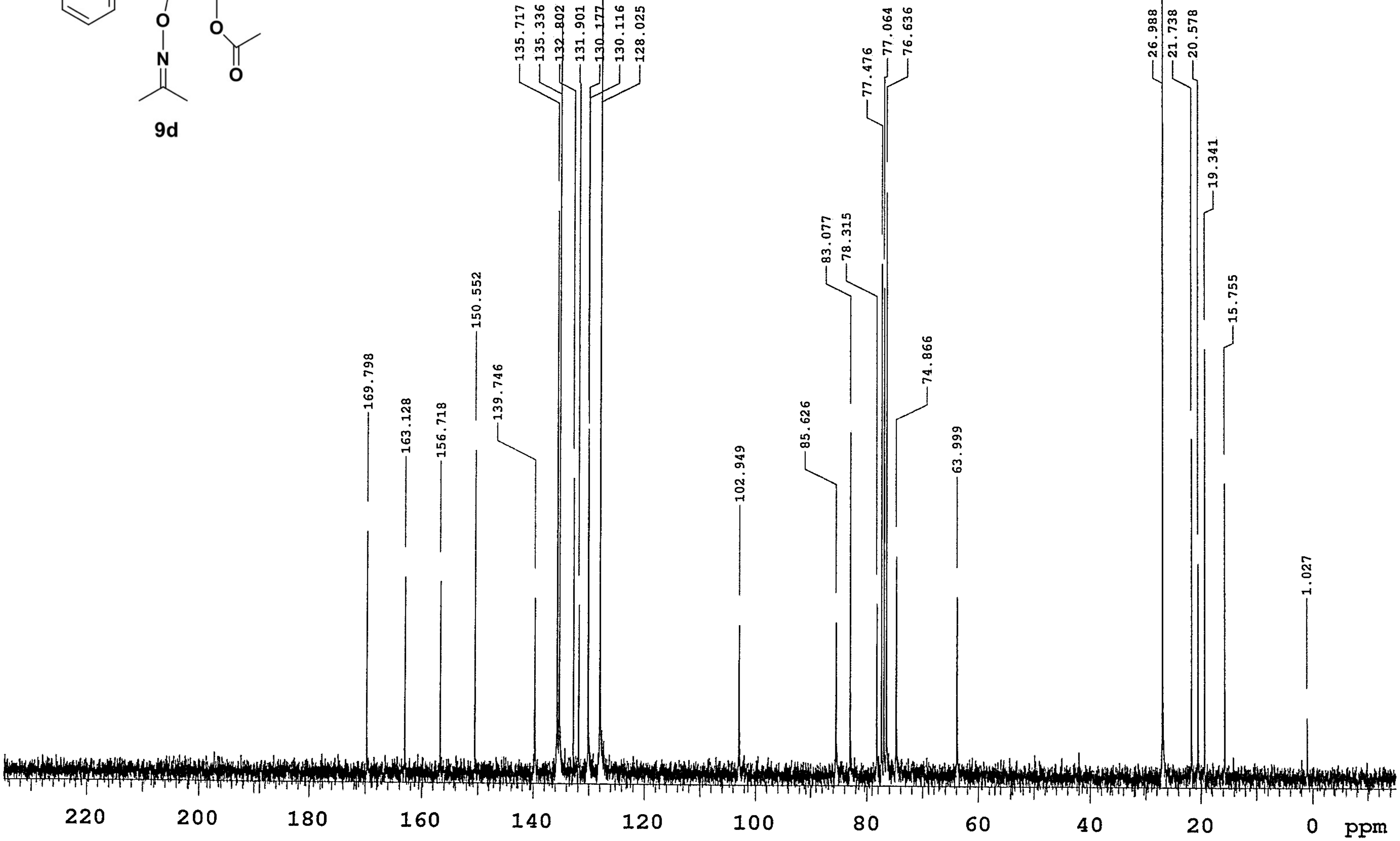
169.798
163.128
156.718
150.552
139.746
135.717
135.336
132.802
131.901
130.177
130.116
128.025
102.949
85.626
83.077
78.315
77.476
77.064
76.636
74.866
63.999
26.988
21.738
20.578
19.341
15.755
1.027
220
200
180
160
140
120
100
80
60
40
20
0
ppm

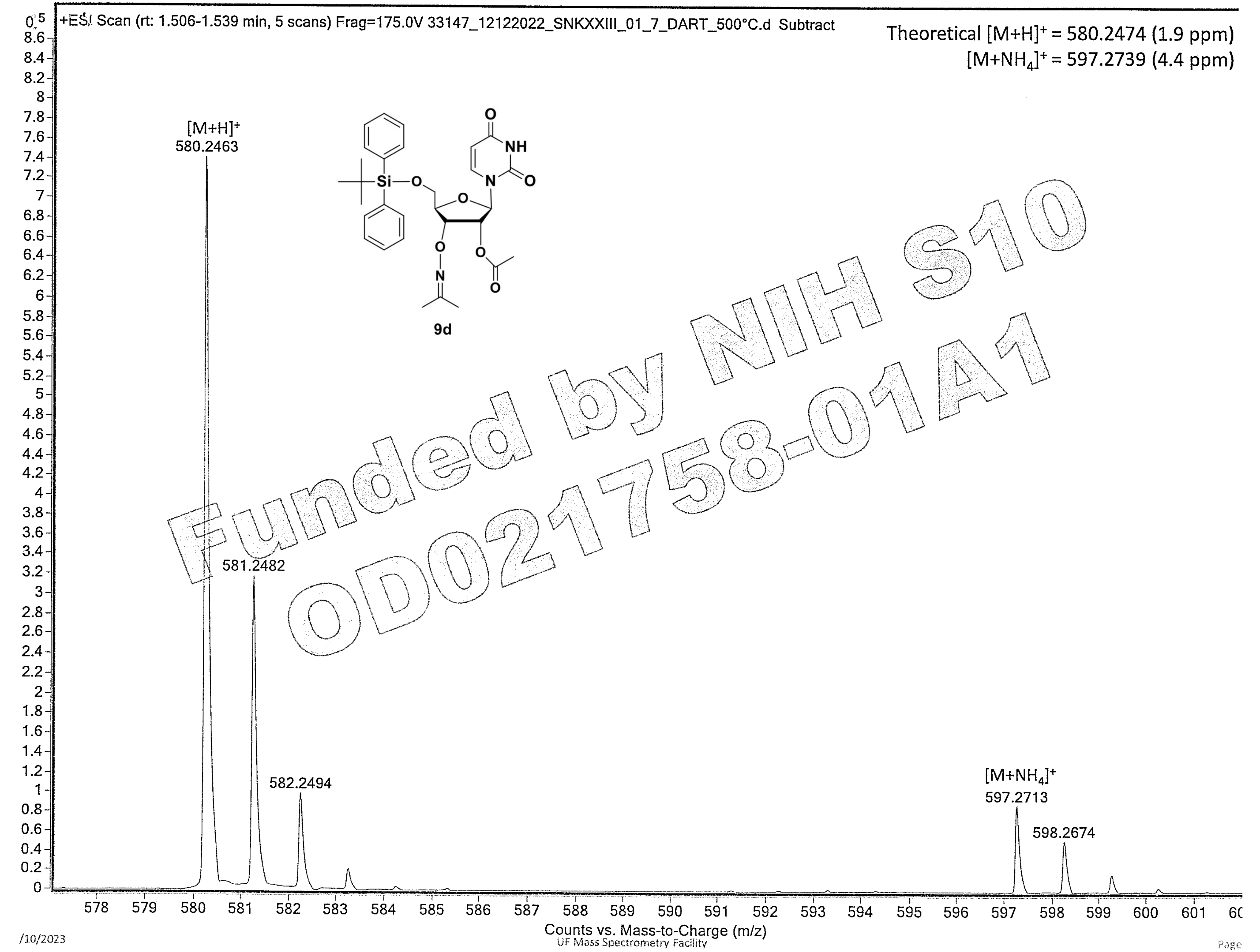

+ESI Scan (rt: 1.506-1.539 min, 5 scans) Frag=175.0V 33147_12122022_SNKXXIII_01_7_DART_500°C.d Subtract
Theoretical [M+H]+ = 580.2474 (1.9 ppm)
[M+NH4]+ = 597.2739 (4.4 ppm)
[M+H]+
580.2463
581.2482
582.2494
[M+NH4]+
597.2713
598.2674
9d
Funded by NIH S10 OD021758-01A1
Counts vs. Mass-to-Charge (m/z)

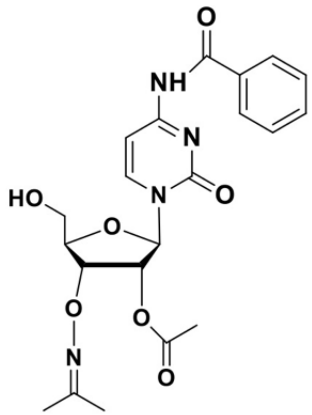

**10a**

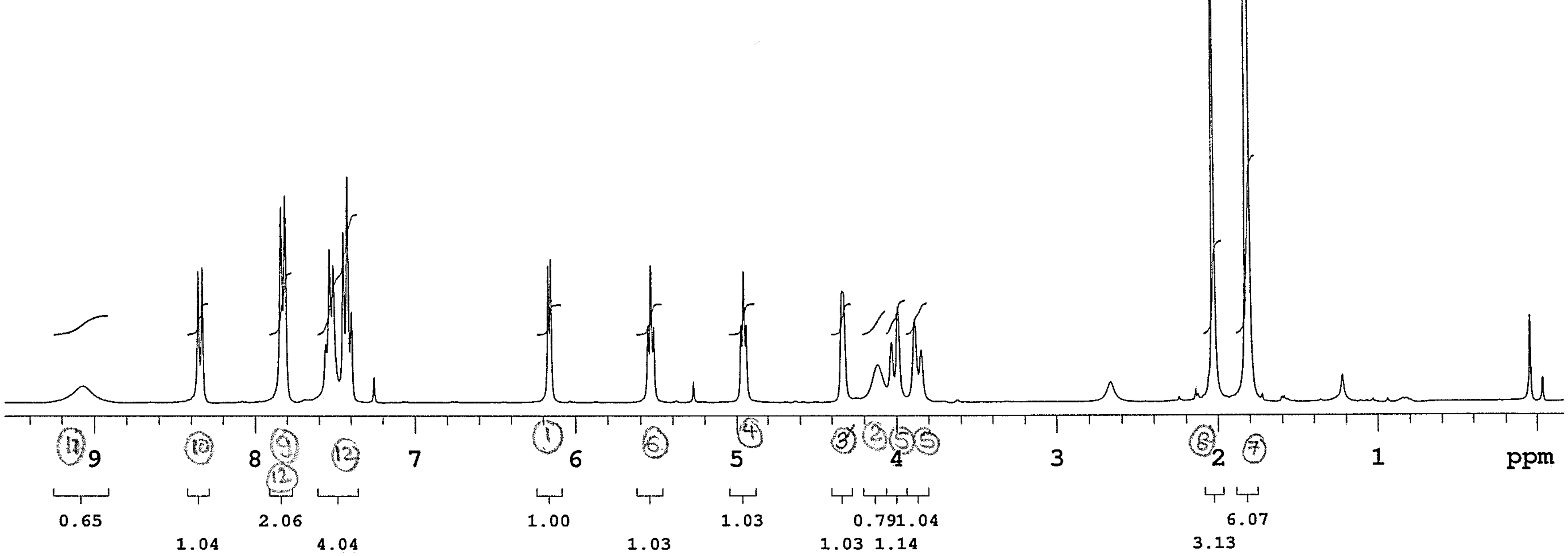

9
8
7
6
5
4
3
2
1
ppm
0.65
1.04
2.06
4.04
1.00
1.03
1.03
1.03
0.79
1.04
1.14
3.13
6.07

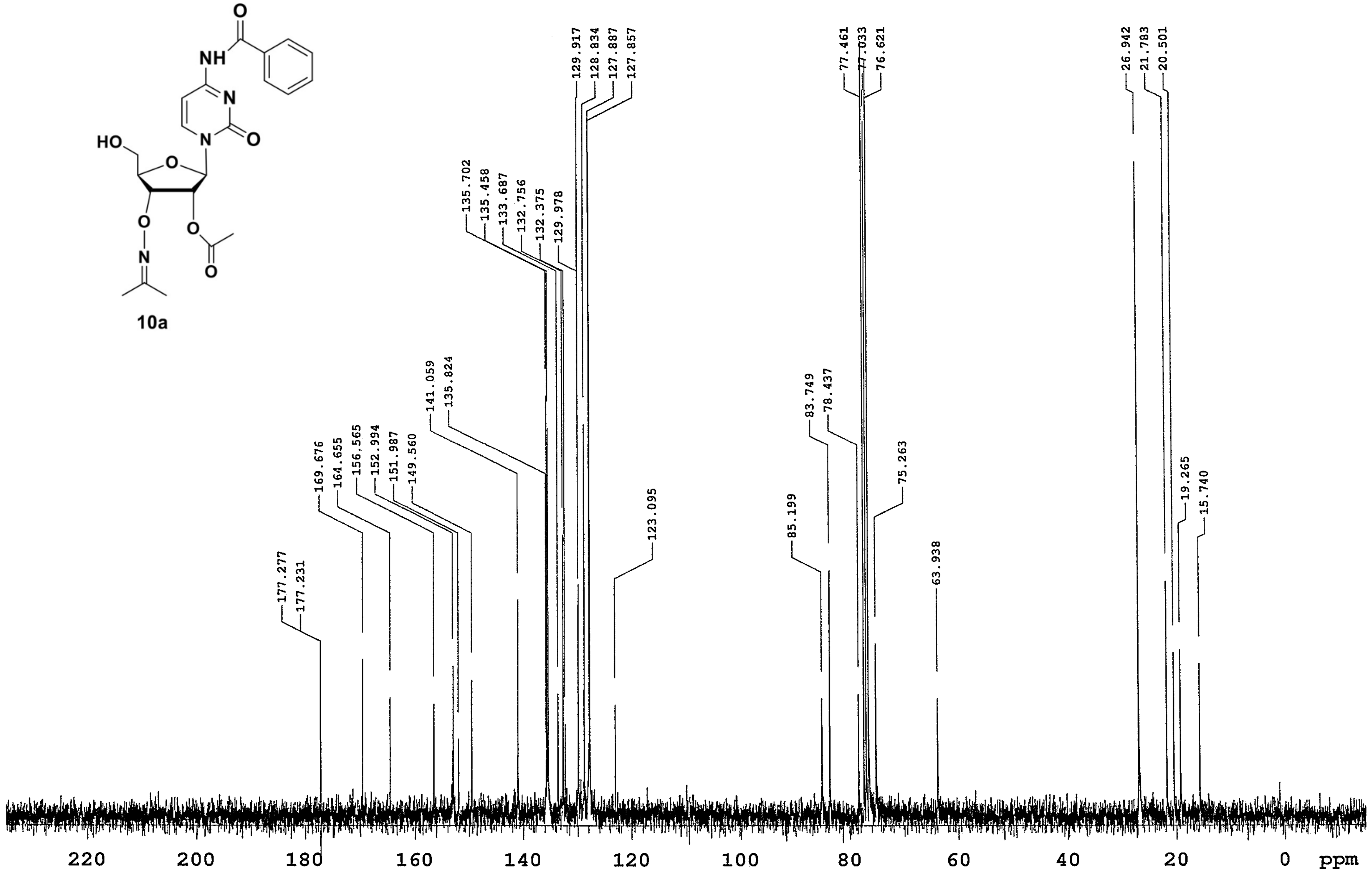
O
NH
N
HO
O
N
O
O
O
N
O
O
177.277
177.231
169.676
164.655
156.565
152.994
151.987
149.560
141.059
135.824
135.702
135.458
133.687
132.756
132.375
129.978
129.917
128.834
127.887
127.857
123.095
85.199
83.749
78.437
77.461
77.033
76.621
75.263
63.938
26.942
21.783
20.501
19.265
15.740
220
200
180
160
140
120
100
80
60
40
20
0
ppm

**10a**

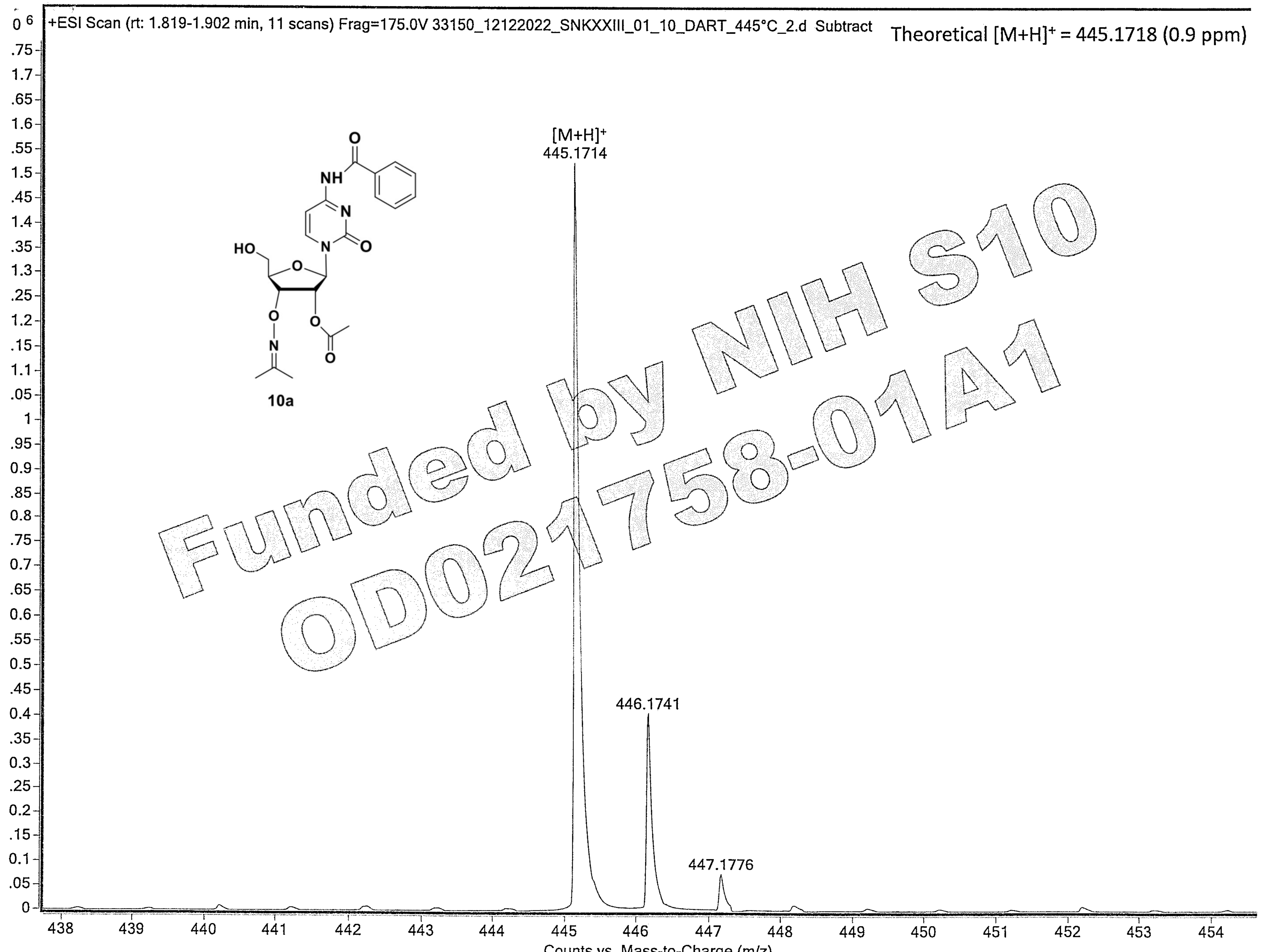

Funded by NIH S10 OD021758-01A1

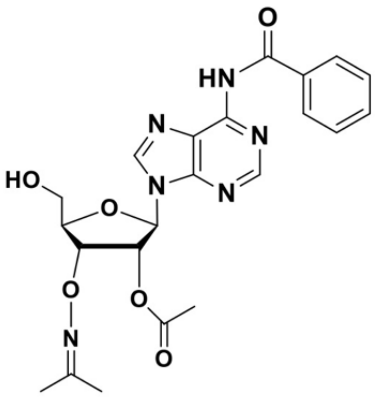

O
HN
N
N
N
N
HO
O
O
N
O
O


**10b**

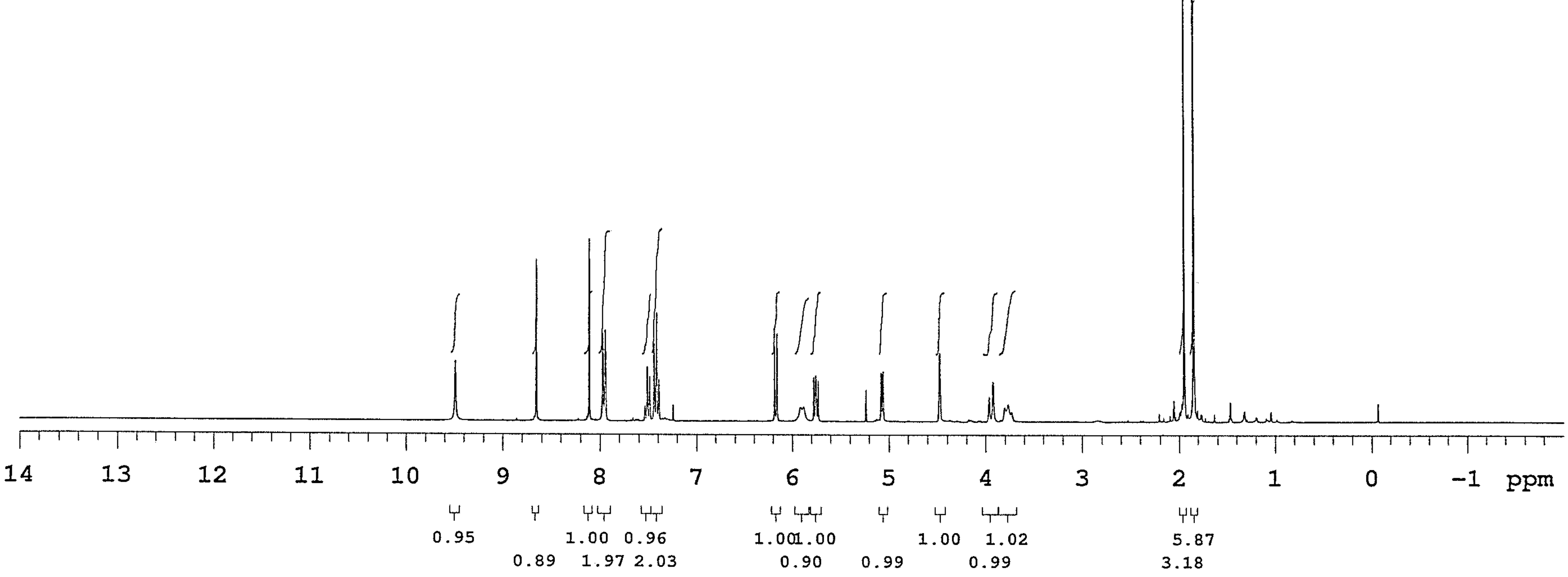

14 13 12 11 10 9 8 7 6 5 4 3 2 1 0 −1 ppm
0.95
0.89
1.00
1.97
0.96
2.03
1.00
1.00
0.90
0.99
1.00
1.02
0.99
5.87
3.18

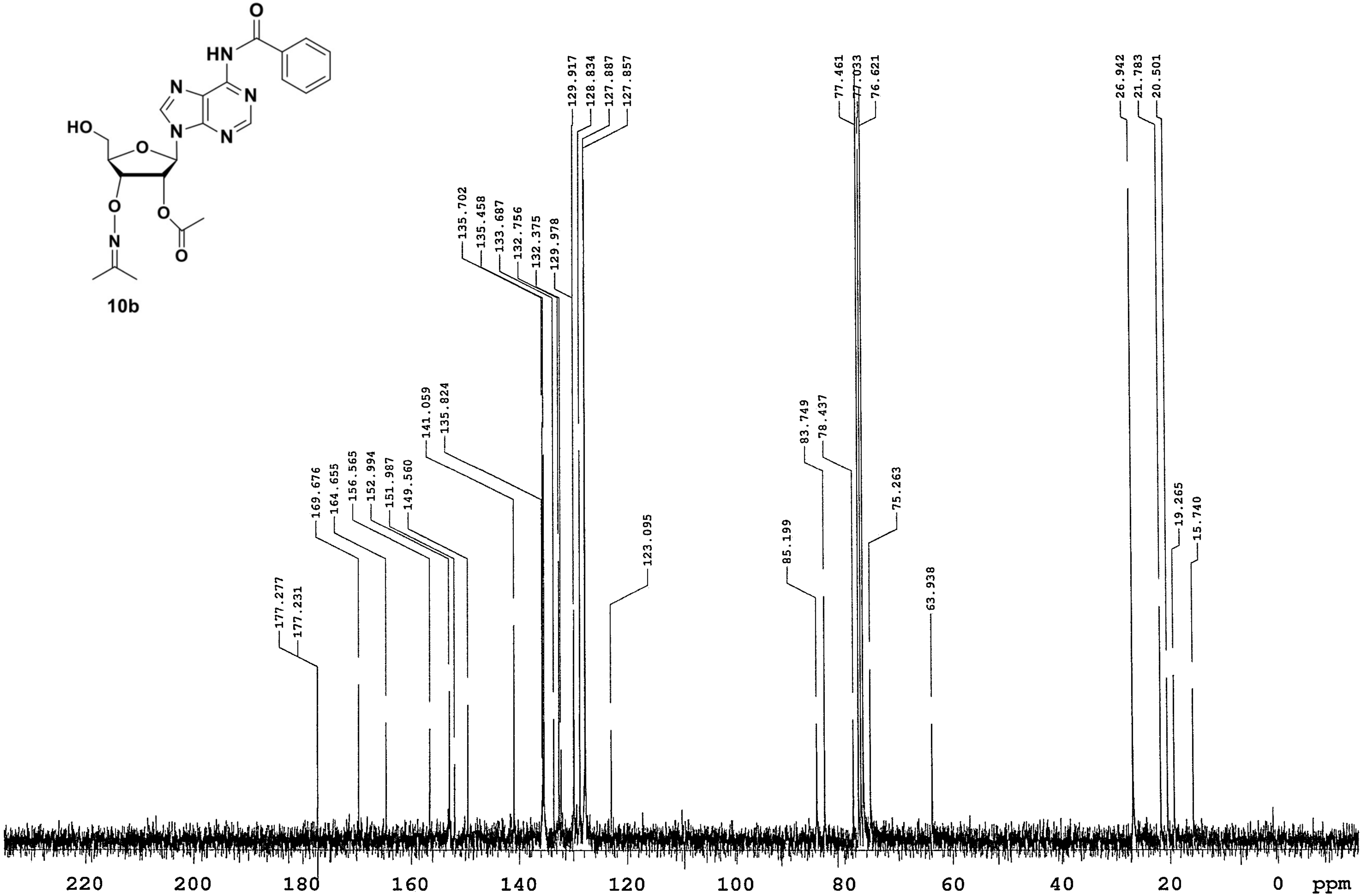
129.917
128.834
127.887
127.857
77.461
77.033
76.621
26.942
21.783
20.501
135.702
135.458
133.687
132.756
132.375
129.978
141.059
135.824
83.749
78.437
169.676
164.655
156.565
152.994
151.987
149.560
75.263
19.265
15.740
123.095
85.199
63.938
177.277
177.231
220
200
180
160
140
120
100
80
60
40
20
0
ppm

10b

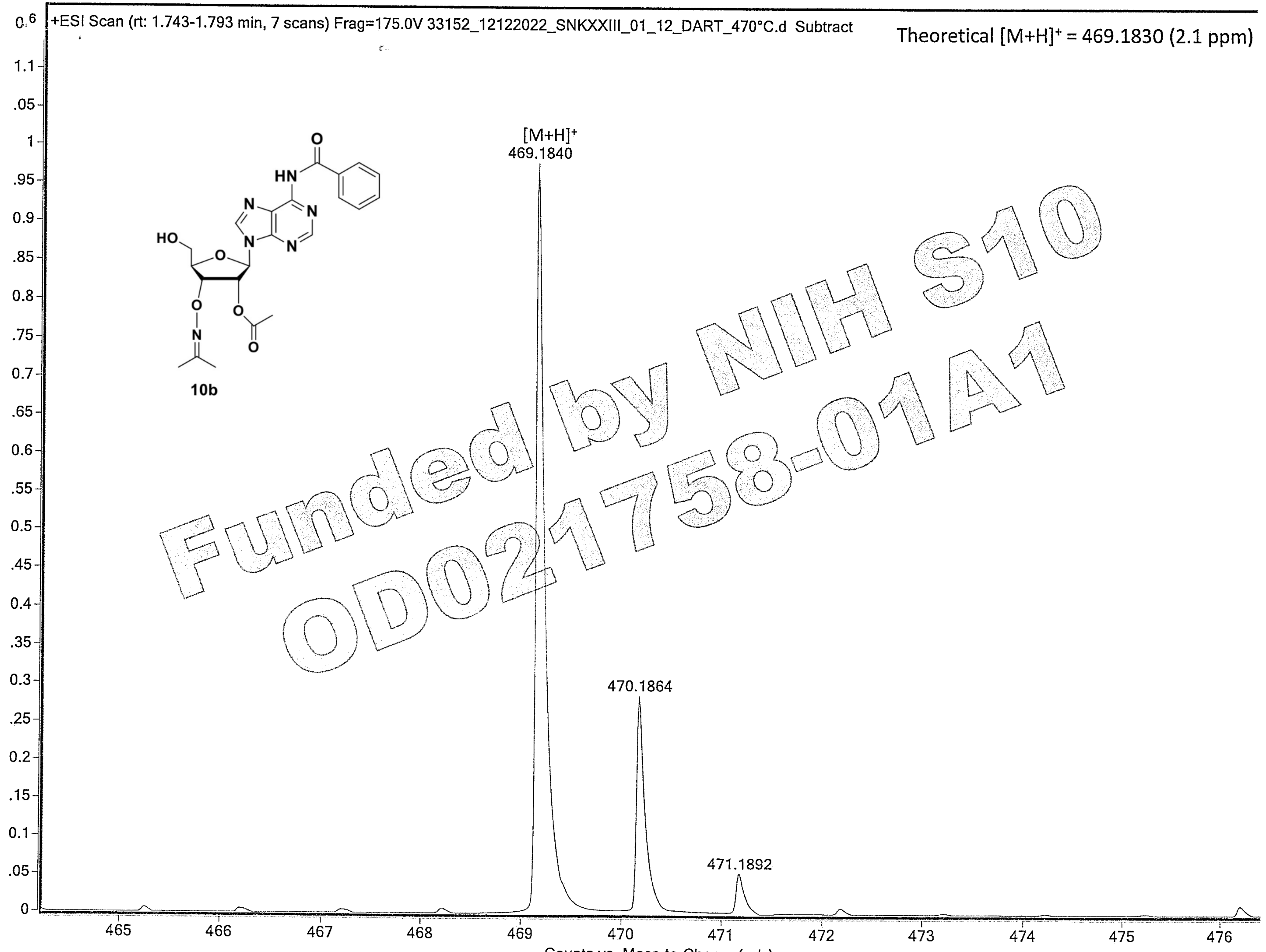

Funded by NIH S10 OD021758-01A1

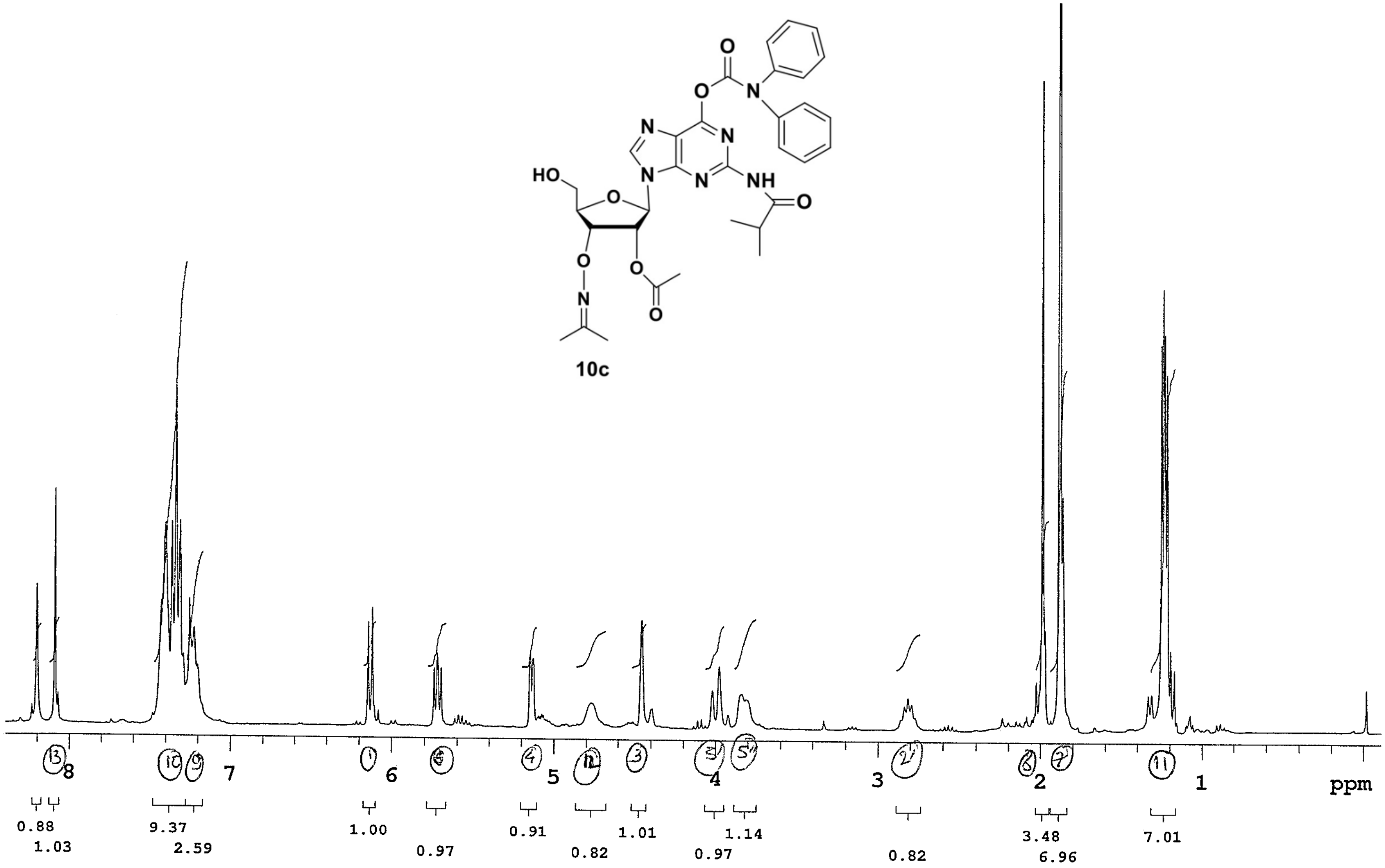

10c
8
7
6
5
4
3
2
1
ppm
0.88
1.03
9.37
2.59
1.00
0.97
0.91
0.82
1.01
0.97
1.14
0.82
3.48
6.96
7.01

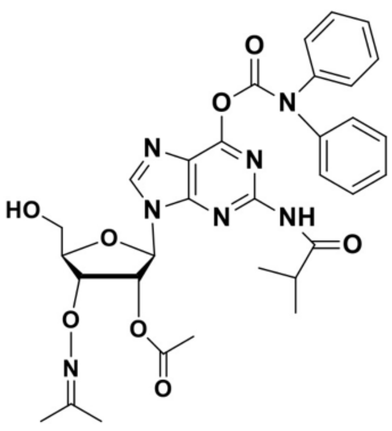

**10c**

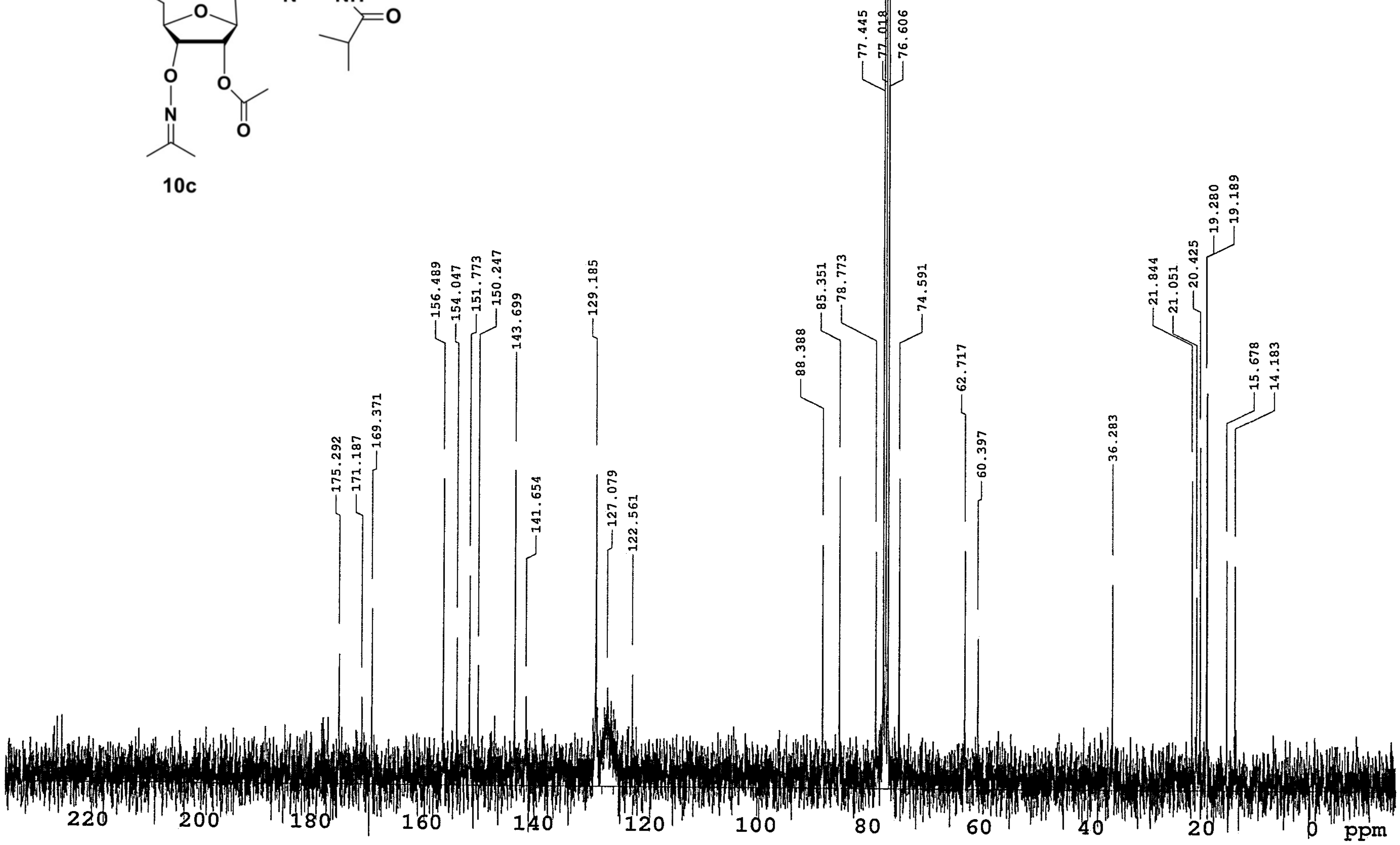

175.292
171.187
169.371
156.489
154.047
151.773
150.247
143.699
141.654
129.185
127.079
122.561
88.388
85.351
78.773
77.445
77.018
76.606
74.591
62.717
60.397
36.283
21.844
21.051
20.425
19.280
19.189
15.678
14.183
220
200
180
160
140
120
100
80
60
40
20
0
ppm

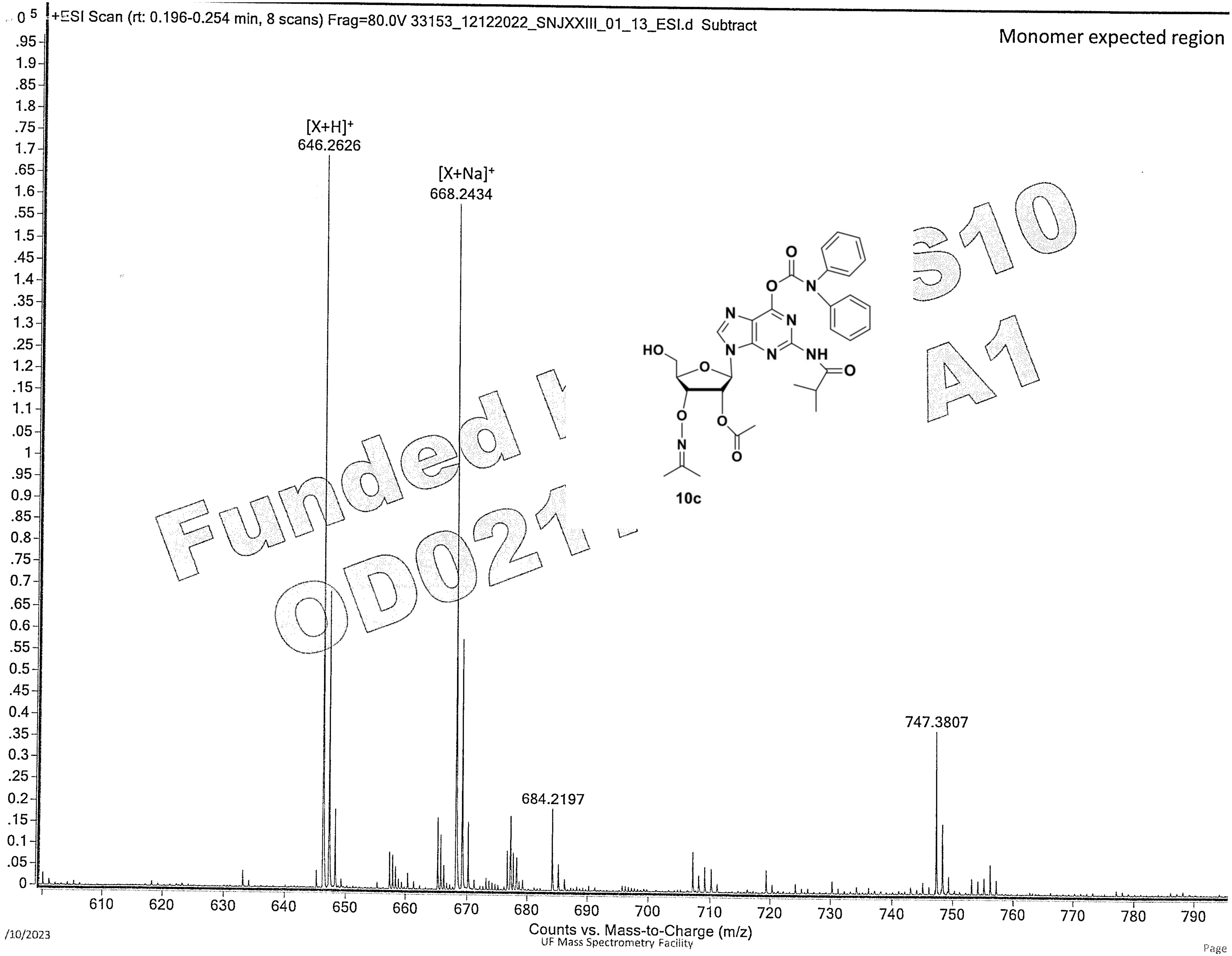
+ESI Scan (rt: 0.196-0.254 min, 8 scans) Frag=80.0V 33153_12122022_SNJXXIII_01_13_ESI.d Subtract
Monomer expected region
[X+H]+
646.2626
[X+Na]+
668.2434
684.2197
747.3807
10c
Counts vs. Mass-to-Charge (m/z)
UF Mass Spectrometry Facility

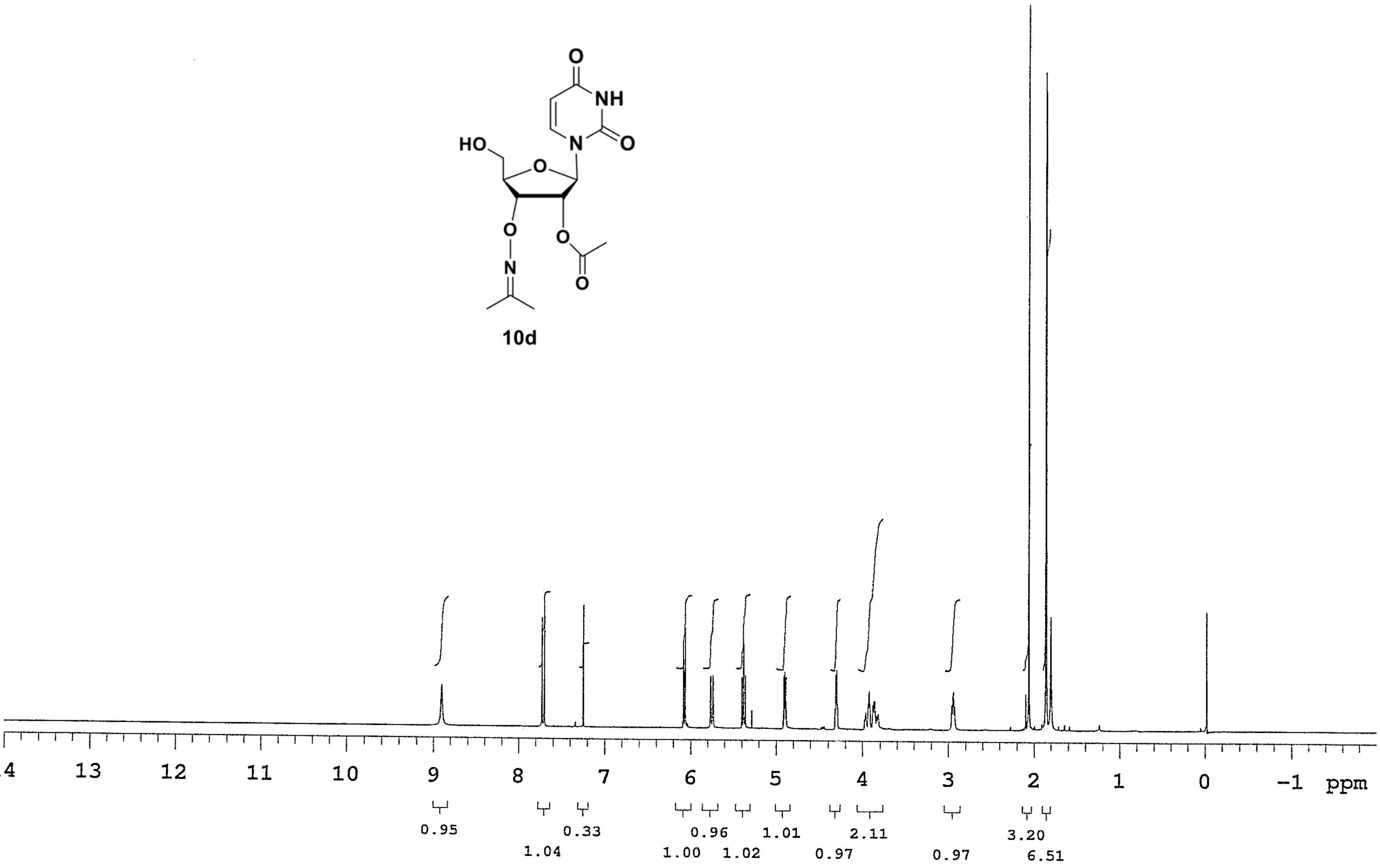

O
NH
N
O
HO
O
O
O
N
O
10d
14 13 12 11 10 9 8 7 6 5 4 3 2 1 0 -1 ppm
0.95
1.04
0.33
1.00
0.96
1.02
1.01
0.97
2.11
0.97
3.20
6.51

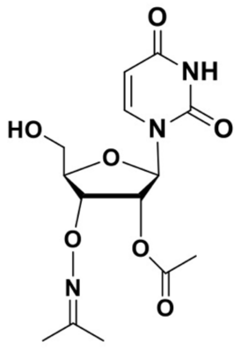

10d

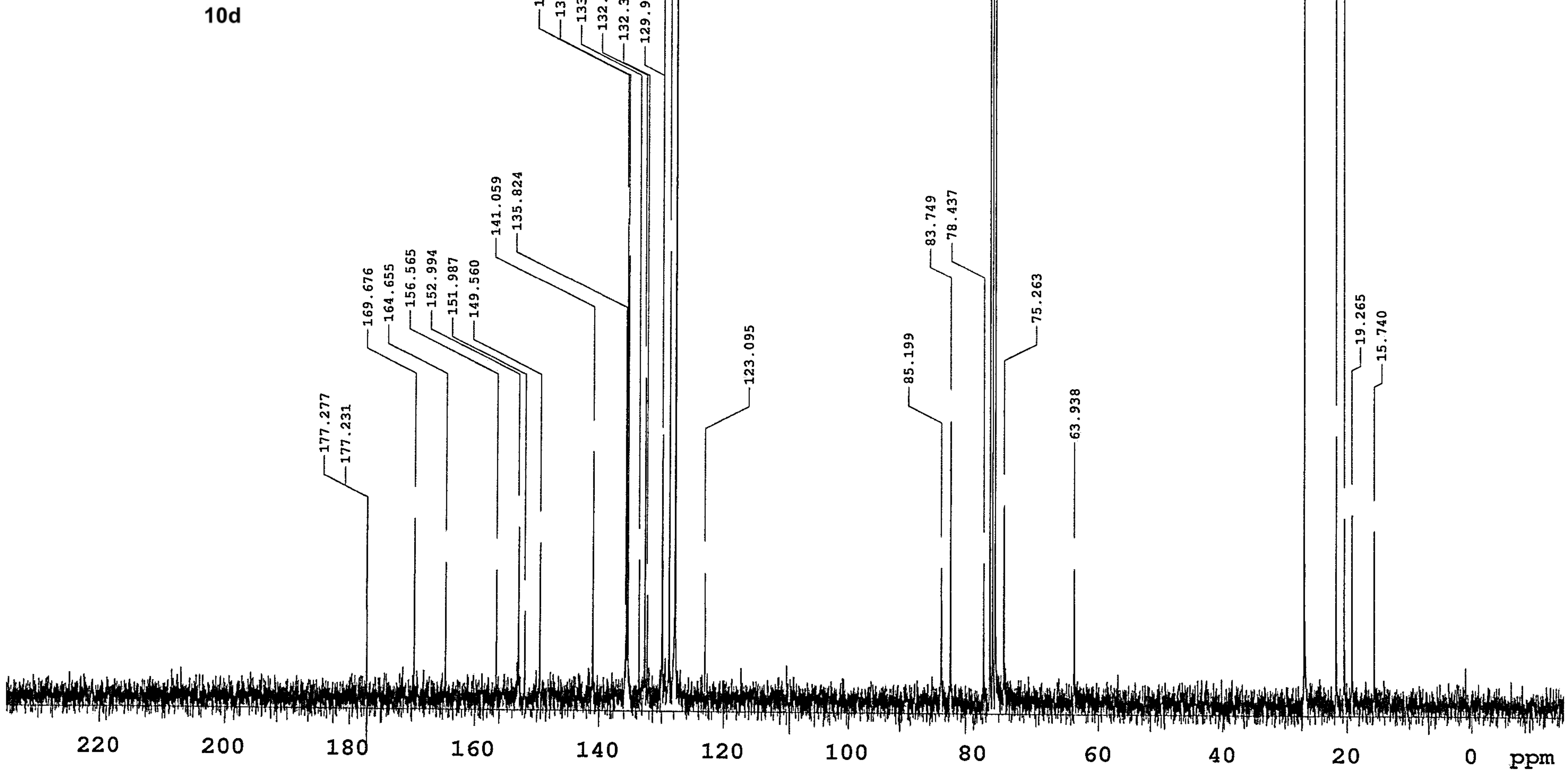

177.277
177.231
169.676
164.655
156.565
152.994
151.987
149.560
141.059
135.824
135.702
135.458
133.687
132.756
132.375
129.978
129.917
128.834
127.887
127.857
123.095
85.199
83.749
78.437
77.461
77.033
76.621
75.263
63.938
26.942
21.783
20.501
19.265
15.740
220
200
180
160
140
120
100
80
60
40
20
0
ppm

Theoretical [M+H]$^+$ = 342.1296 (2.9 ppm)
[M+NH$_4$]$^+$ = 359.1561 (3.6 ppm)

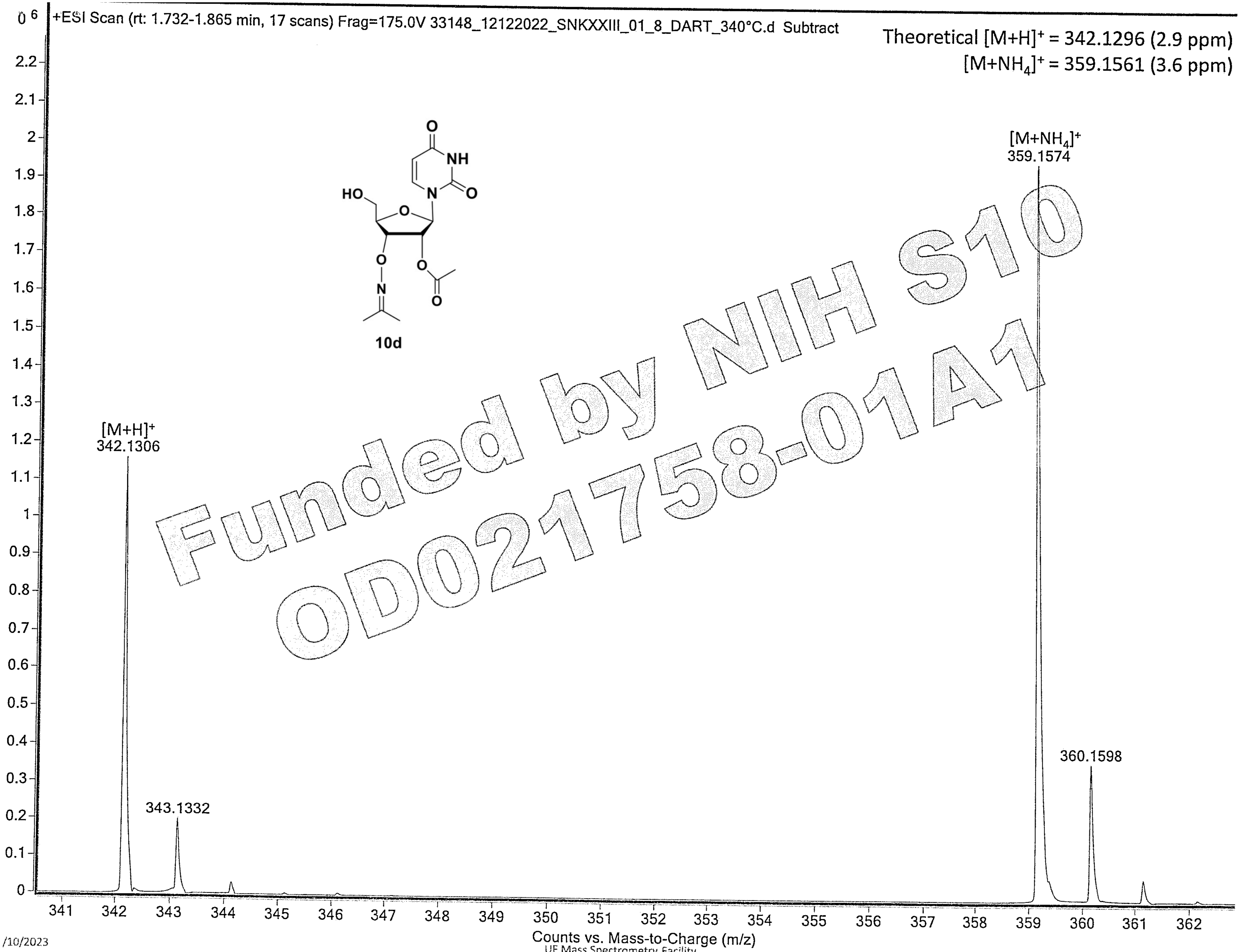


Funded by NIH S10 OD021758-01A1

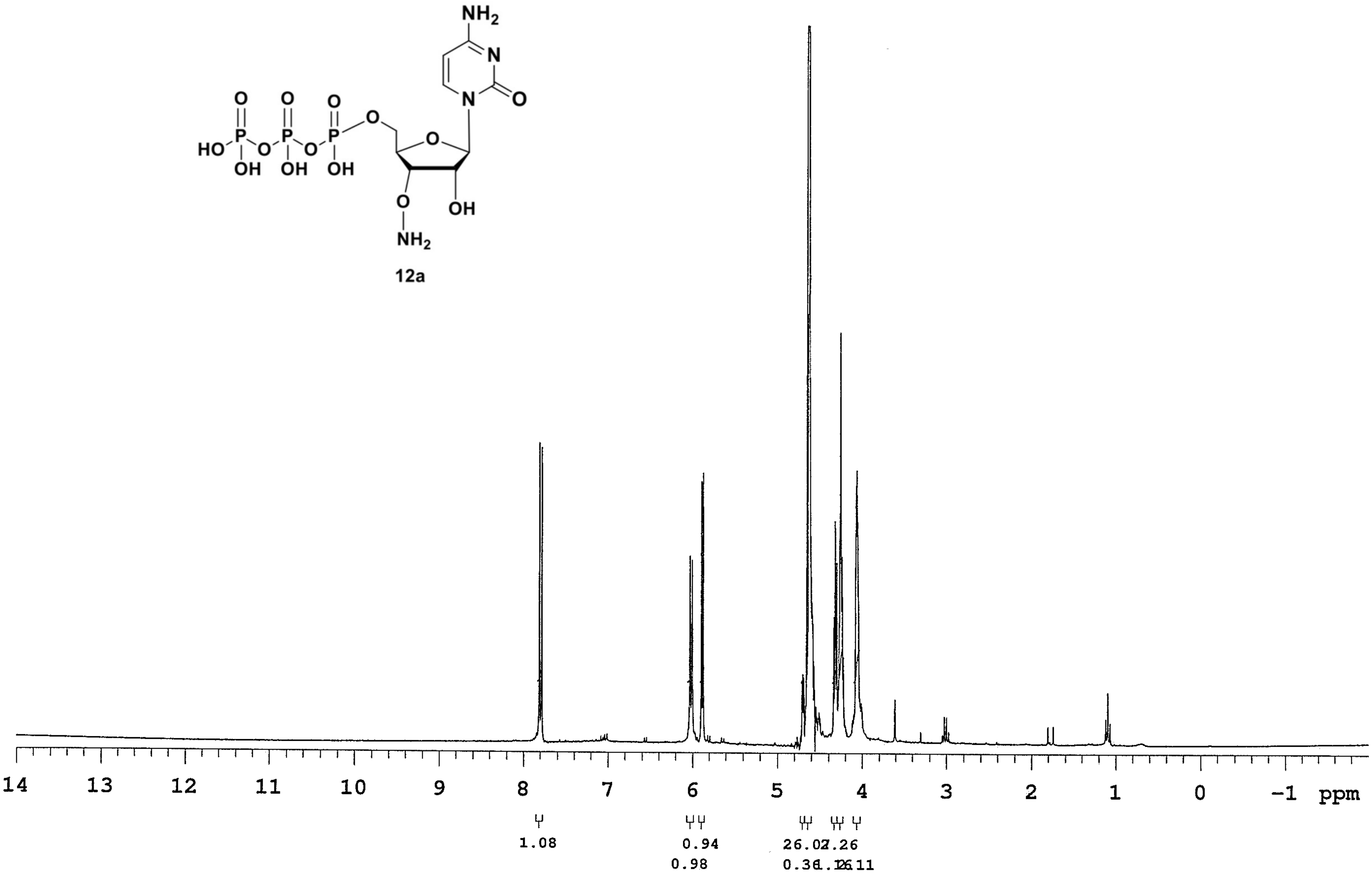
NH2
N
N
O
O
HO
P
OH
OH
OH
NH2
12a
14 13 12 11 10 9 8 7 6 5 4 3 2 1 0 -1 ppm
1.08
0.94
0.98
26.02
7.26
0.36
1.12
6.11

**012222_42_SNKXXII_CAminoxy**

| | | | | | | | |
|---|---|---|---|---|---|---|---|
| Sample Name | 012222_42_SNKXXII_CAminoxy | Pulse sequence | PHOSPHORUS | Temperature | 25 | Study owner | nkaralkar |
| Date collected | 2022-02-03 | Solvent | d2o | Spectrometer | M300-mercury300 | Operator | nkaralkar |

**012222_42_SNKXXII_CAminoxy**

**SAMPLE**

| | |
|---|---|
| date | Feb 3 2022 |
| solvent | d2o |
| file | /home/nkaralkar/vnmrsys/data/012222_42_SNKXXII_CAminoxy_20220203_01/PHOSPHORUS_01.fid |

**ACQUISITION**

| | |
|---|---|
| sw | 5830.9 |
| at | 1.405 |
| np | 16384 |
| fb | 3200 |
| bs | 64 |
| d1 | 1.000 |
| nt | 512 |
| ct | 512 |

**TRANSMITTER**

| | |
|---|---|
| tn | P31 |
| sfrq | 121.475 |
| tof | 2676.6 |
| tpwr | 59 |
| pw | 8.000 |

**DECOUPLER**

| | |
|---|---|
| dn | H1 |
| dof | 0 |
| dm | nny |
| decwave | w |
| dpwr | 41 |
| dmf | 7400 |

**PRESATURATION**

| | |
|---|---|
| satmode | n |
| wet | n |

**SPECIAL**

| | |
|---|---|
| temp | 25.0 |
| gain | 38 |
| spin | 20 |
| hst | 0.008 |
| pw90 | 16.000 |
| alfa | 10.000 |

**FLAGS**

| | |
|---|---|
| il | n |
| in | n |
| dp | y |
| hs | nn |

**PROCESSING**

| | |
|---|---|
| lb | 0.50 |
| lsfid | -2 |
| fn | 16384 |

**DISPLAY**

| | |
|---|---|
| sp | -3401.3 |
| wp | 5830.9 |
| rfl | 3401.3 |
| rfp | 0 |
| rp | -31.8 |
| lp | 636.8 |

**PLOT**

| | |
|---|---|
| wc | 234 |
| sc | 8 |
| vs | 616 |
| th | 9 |
| ai cdc ph | |

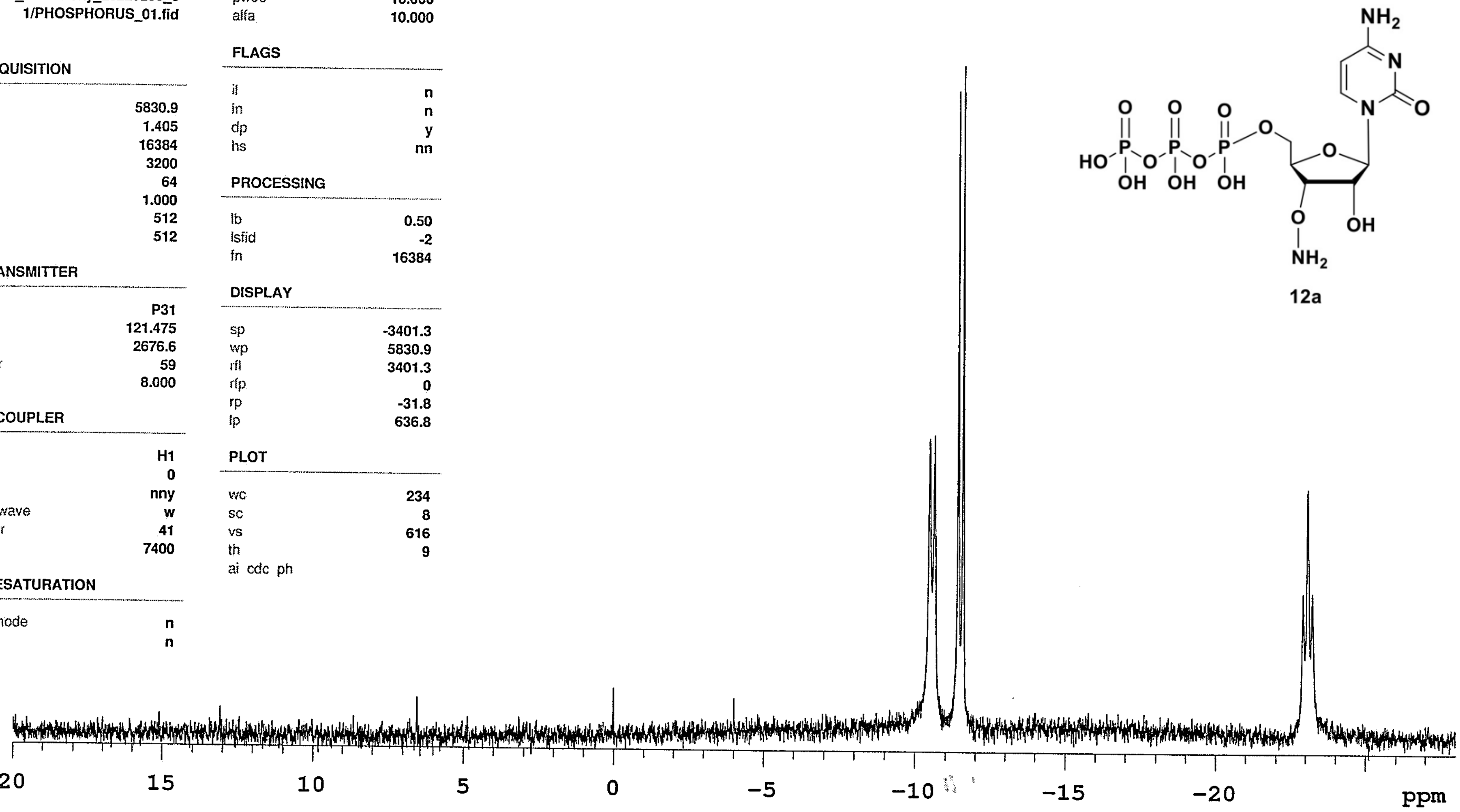


**12a**

Data file /home/nkaralkar/vnmrsys/data/012222_42_SNKXXII_CAminoxy_20220203_01/PHOSPHORUS_01.fid

Plot date 2022-02-03

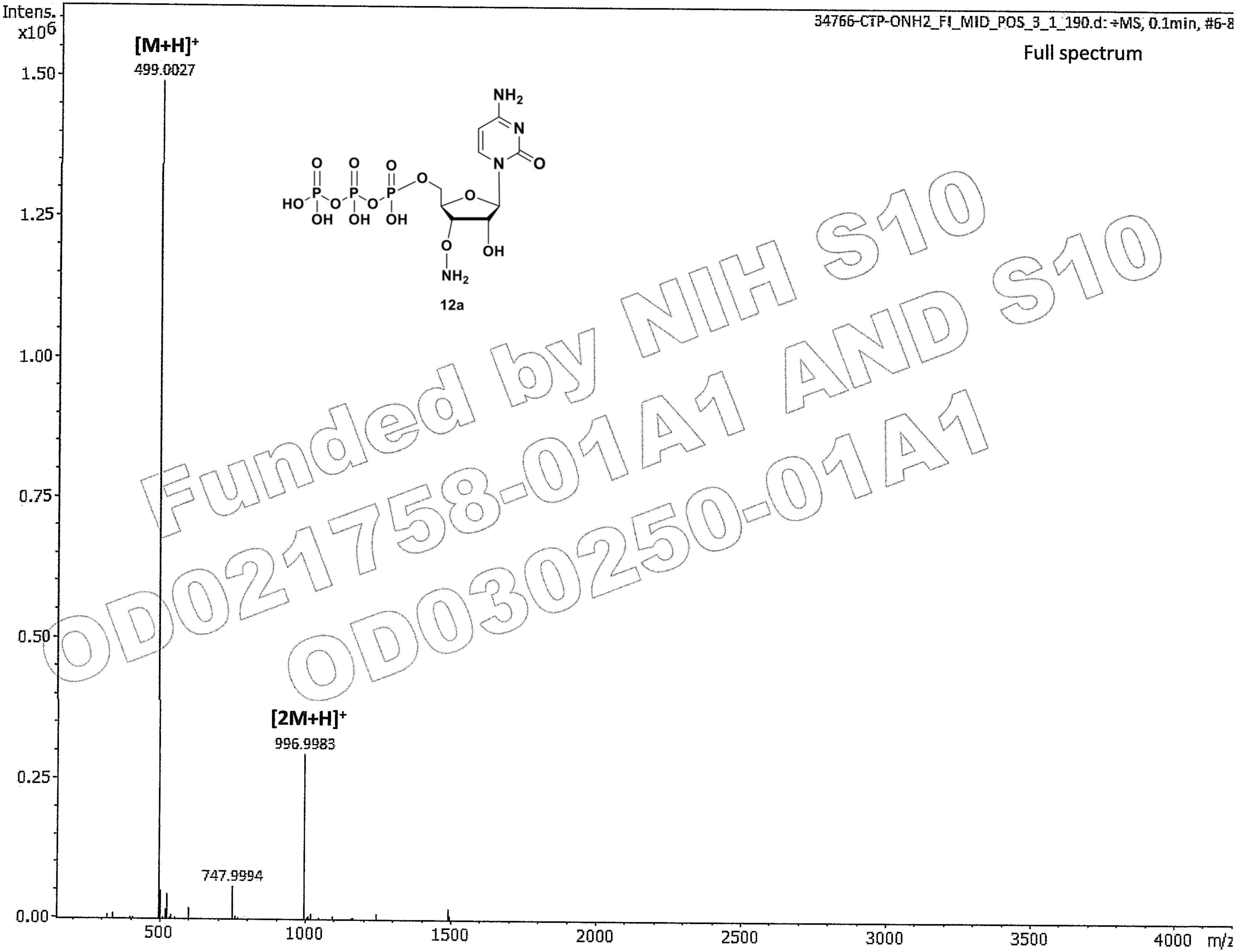


Funded by NIH S10 OD021758-01A1 AND S10 OD030250-01A1

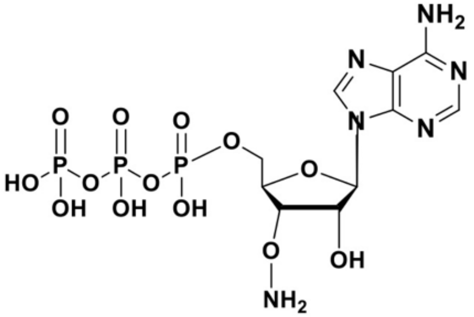
NH2
N
N
N
N
O
O
O
O
HO
P
O
P
O
P
O
OH
OH
OH
O
OH
NH2

12b

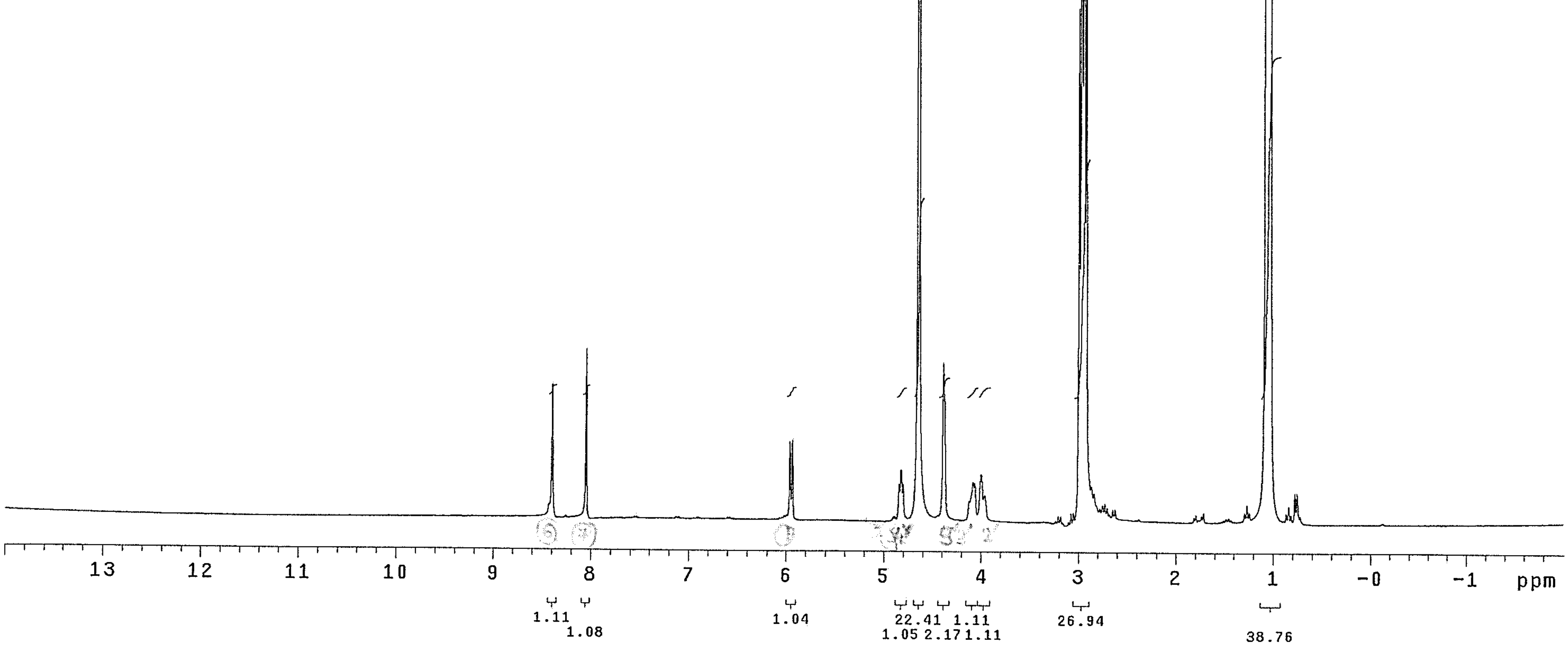
13
12
11
10
9
8
7
6
5
4
3
2
1
-0
-1
ppm
1.11
1.08
1.04
22.41
1.05
2.17
1.11
1.11
26.94
38.76

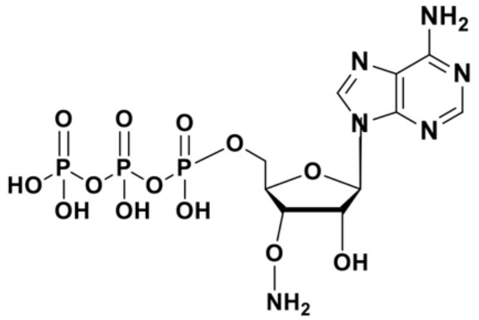
NH2
N
N
N
N
O
O
O
O
HO
P
P
P
O
O
O
OH
OH
OH
OH
O
O
NH2

12b

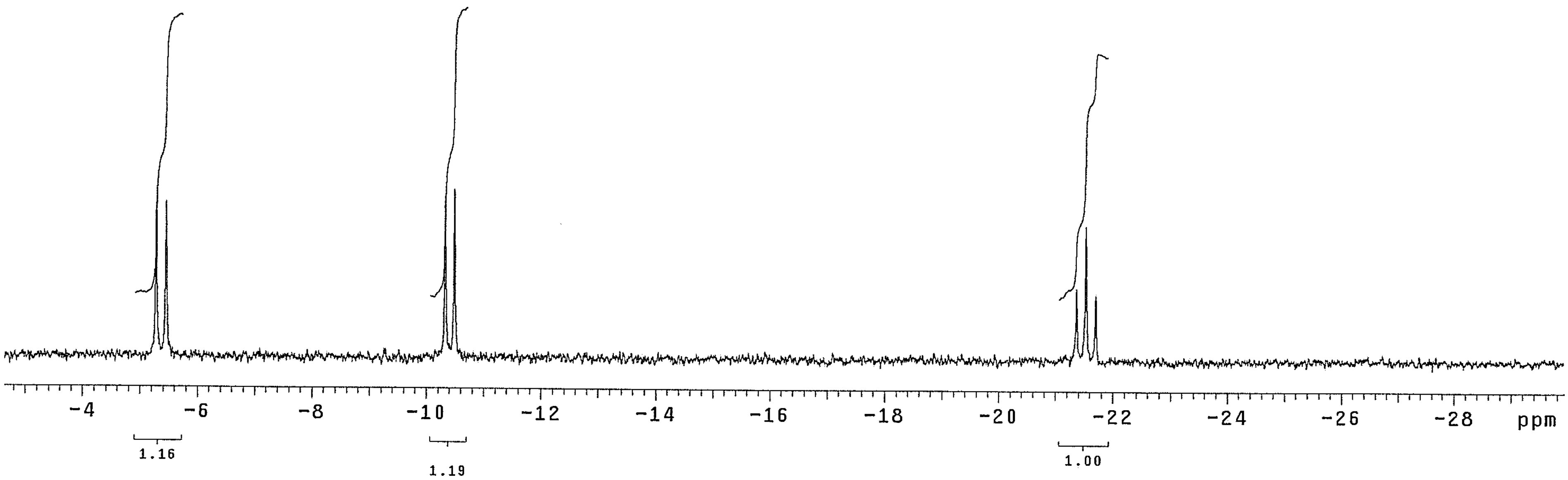
-4
-6
-8
-10
-12
-14
-16
-18
-20
-22
-24
-26
-28
ppm
1.16
1.19
1.00

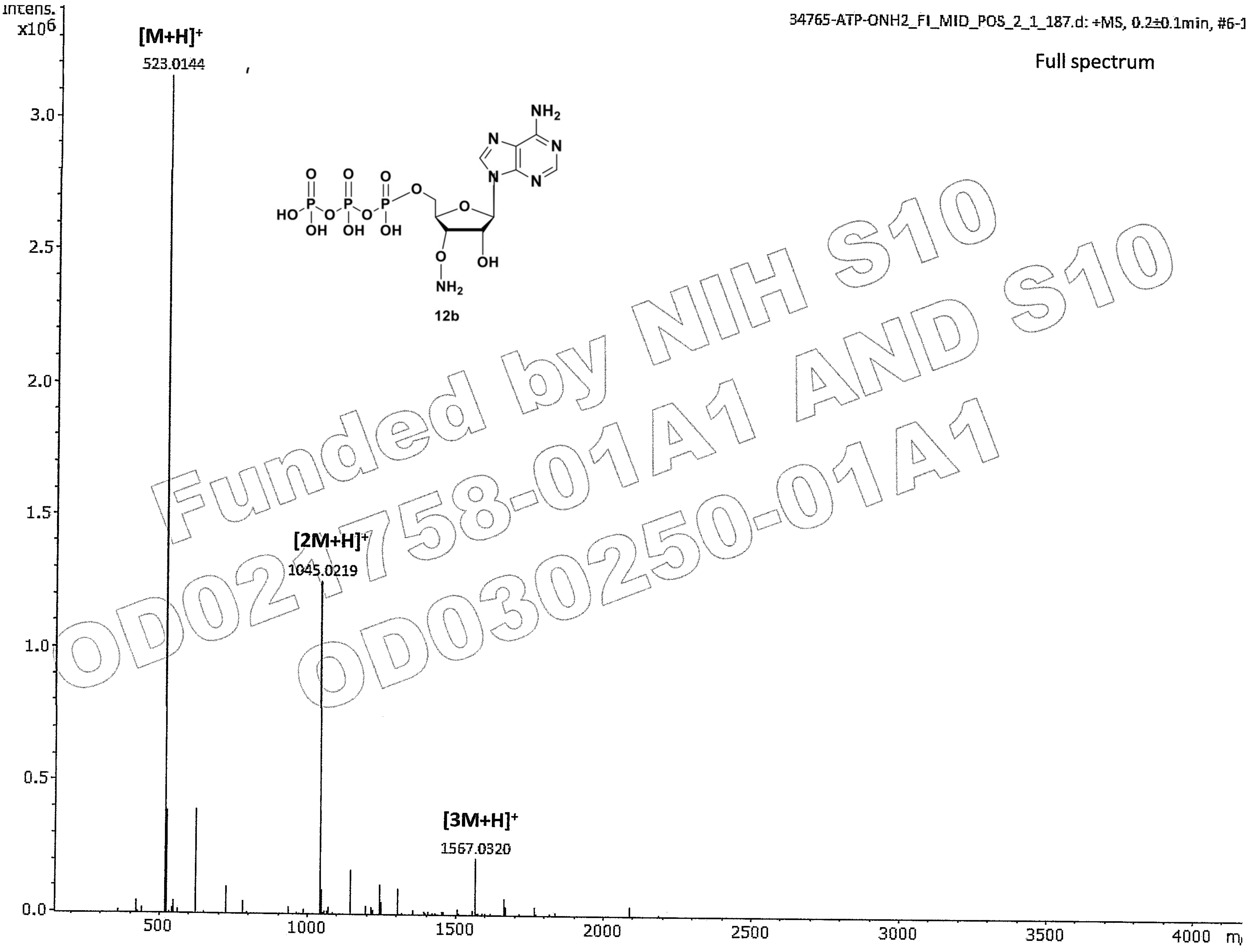

Funded by NIH S10 OD021758-01A1 AND S10 OD030250-01A1

012822_44_SNKXXII_D2O_GAminoxy

| | | | | | |
|---|---|---|---|---|---|
| Sample Name | 012822_44_SNKXXII_D2O_GAminoxy | uence | PROTON | Temperature | 25 |
| Date collected | 2022-02-08 | Solvent | d2o | Spectrometer | M300-mercury300 |

Study owner nkaralkar
Operator nkaralkar

012822_44_SNKXXII_D2O_GAminoxy

**SAMPLE**

| | |
|---|---|
| date | Feb 8 2022 |
| solvent | d2o |
| file | /home/nkaralkar/vnmrsys/data/012822_44_SNKXXII_D2O_GAminoxy_20220208_01/PROTON_01.fid |

**ACQUISITION**

| | |
|---|---|
| sw | 4800.8 |
| at | 1.706 |
| np | 16384 |
| fb | 2600 |
| bs | 32 |
| d1 | 1.000 |
| nt | 512 |
| ct | 512 |

**TRANSMITTER**

| | |
|---|---|
| tn | H1 |
| sfrq | 300.085 |
| tof | 258.6 |
| tpwr | 61 |
| pw | 9.250 |

**DECOUPLER**

| | |
|---|---|
| dn | C13 |
| dof | 0 |
| dm | nnn |
| decwave | g |
| dpwr | 42 |
| dmf | 12300 |

**PRESATURATION**

| | |
|---|---|
| satmode | n |
| wet | n |

**SPECIAL**

| | |
|---|---|
| temp | 25.0 |
| gain | 22 |
| spin | 20 |
| hst | 0.008 |
| pw90 | 18.500 |
| alfa | 10.000 |

**FLAGS**

| | |
|---|---|
| il | n |
| in | n |
| dp | y |
| hs | nn |

**PROCESSING**

| | |
|---|---|
| fn | not used |

**DISPLAY**

| | |
|---|---|
| sp | -599.3 |
| wp | 4800.2 |
| rfl | 599.9 |
| rfp | 0 |
| rp | -150.6 |
| lp | -37.8 |

**PLOT**

| | |
|---|---|
| wc | 234 |
| sc | 8 |
| vs | 1232 |
| th | 0 |
| ai cdc ph | |

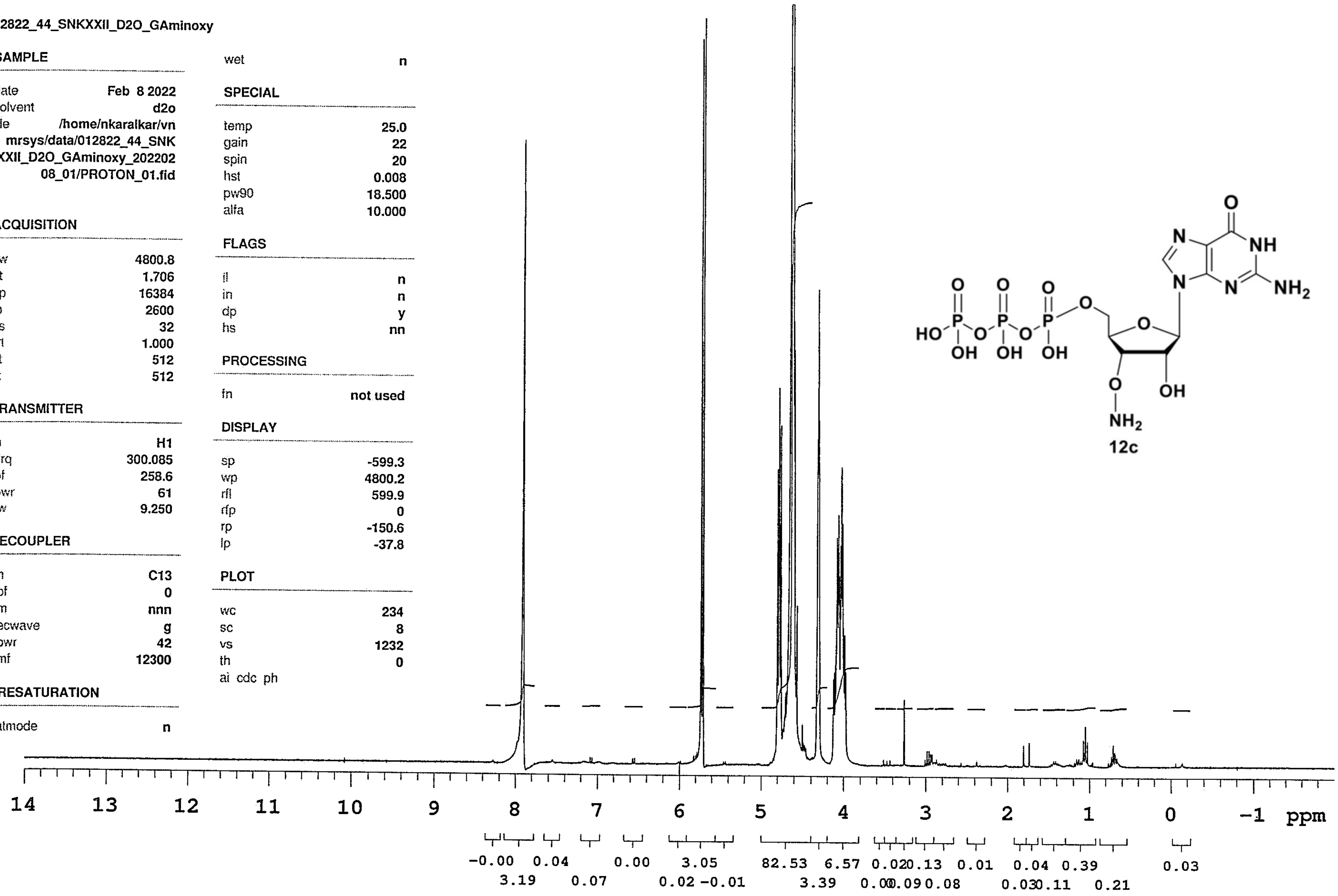


12c

Data file /home/nkaralkar/vnmrsys/data/012822_44_SNKXXII_D2O_GAminoxy_20220208_01/PROTON_01.fid

Plot date 2022-02-08

012822_44_SNKXXII_D2O_P31

| | | | |
|---|---|---|---|
| Sample Name 012822_44_SNKXXII_D2O_P31 | Pulse sequence PHOSPHORUS | Temperature 25 | Study owner nkaralkar |
| Date collected 2022-02-08 | Solvent d2o | Spectrometer M300-mercury300 | Operator nkaralkar |

012822_44_SNKXXII_D2O_P31

**SAMPLE**

| | |
|---|---|
| date | Feb 8 2022 |
| solvent | d2o |
| file | /home/nkaralkar/vnmrsys/data/012822_44_SNKXXII_D2O_P31_20220208_01/PHOSPHORUS_01.fid |

**ACQUISITION**

| | |
|---|---|
| sw | 5830.9 |
| at | 1.405 |
| np | 16384 |
| fb | 3200 |
| bs | 64 |
| d1 | 1.000 |
| nt | 512 |
| ct | 512 |

**TRANSMITTER**

| | |
|---|---|
| tn | P31 |
| sfrq | 121.475 |
| tof | 2676.6 |
| tpwr | 59 |
| pw | 8.000 |

**DECOUPLER**

| | |
|---|---|
| dn | H1 |
| dof | 0 |
| dm | nny |
| decwave | w |
| dpwr | 41 |
| dmf | 7400 |

**PRESATURATION**

| | |
|---|---|
| satmode | n |
| wet | n |

**SPECIAL**

| | |
|---|---|
| temp | 25.0 |
| gain | 38 |
| spin | 20 |
| hst | 0.008 |
| pw90 | 16.000 |
| alfa | 10.000 |

**FLAGS**

| | |
|---|---|
| il | n |
| in | n |
| dp | y |
| hs | nn |

**PROCESSING**

| | |
|---|---|
| lb | 0.50 |
| lsfid | -2 |
| fn | 16384 |

**DISPLAY**

| | |
|---|---|
| sp | -3401.4 |
| wp | 5830.9 |
| rfl | 3401.4 |
| rfp | 0 |
| rp | -16.5 |
| lp | 581.9 |

**PLOT**

| | |
|---|---|
| wc | 234 |
| sc | 8 |
| vs | 782 |
| th | 11 |
| ai cdc ph | |

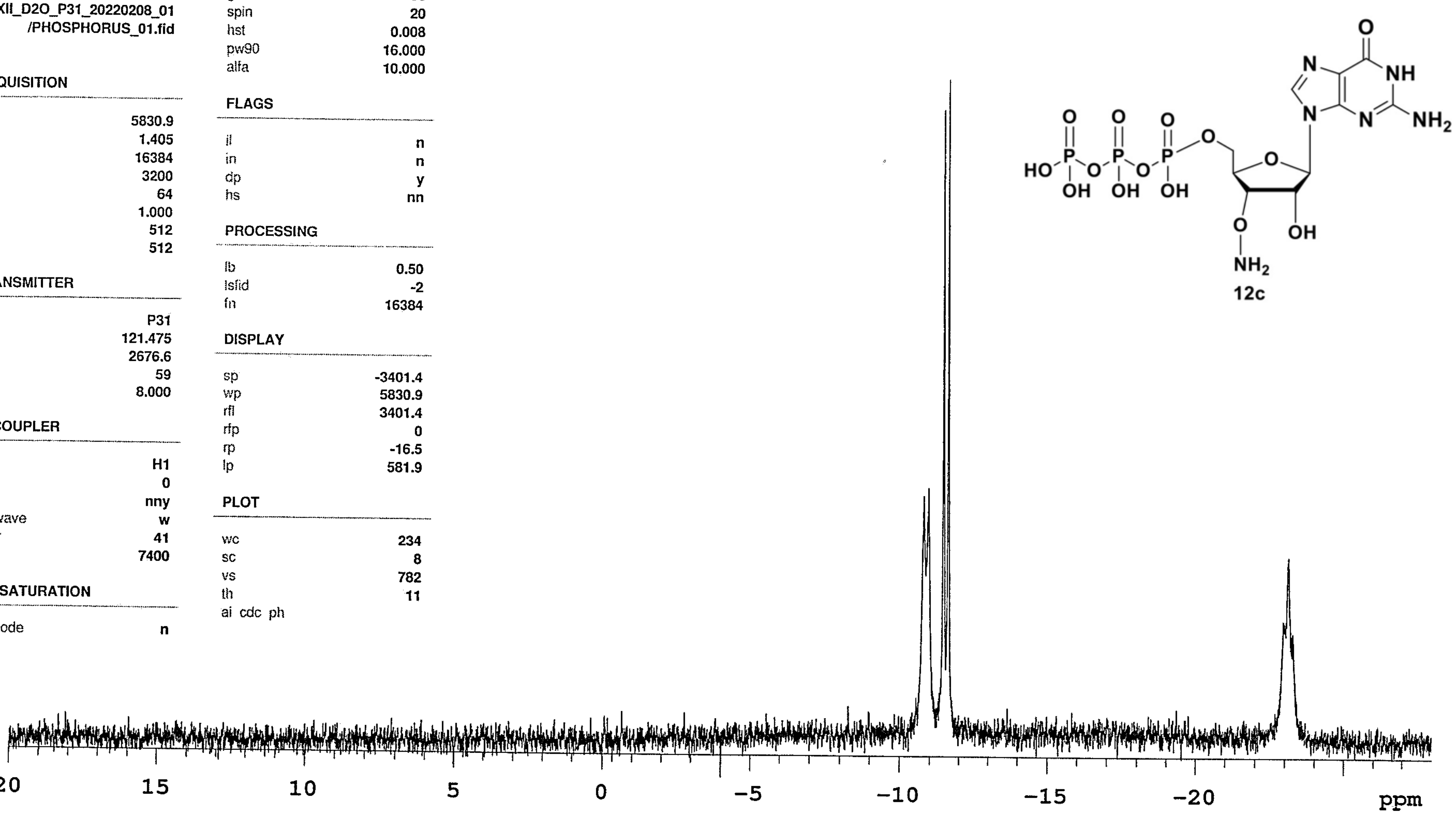


12c

Data file /home/nkaralkar/vnmrsys/data/012822_44_SNKXXII_D2O_P31_20220208_01/PHOSPHORUS_01.fid

Plot date 2022-02-08

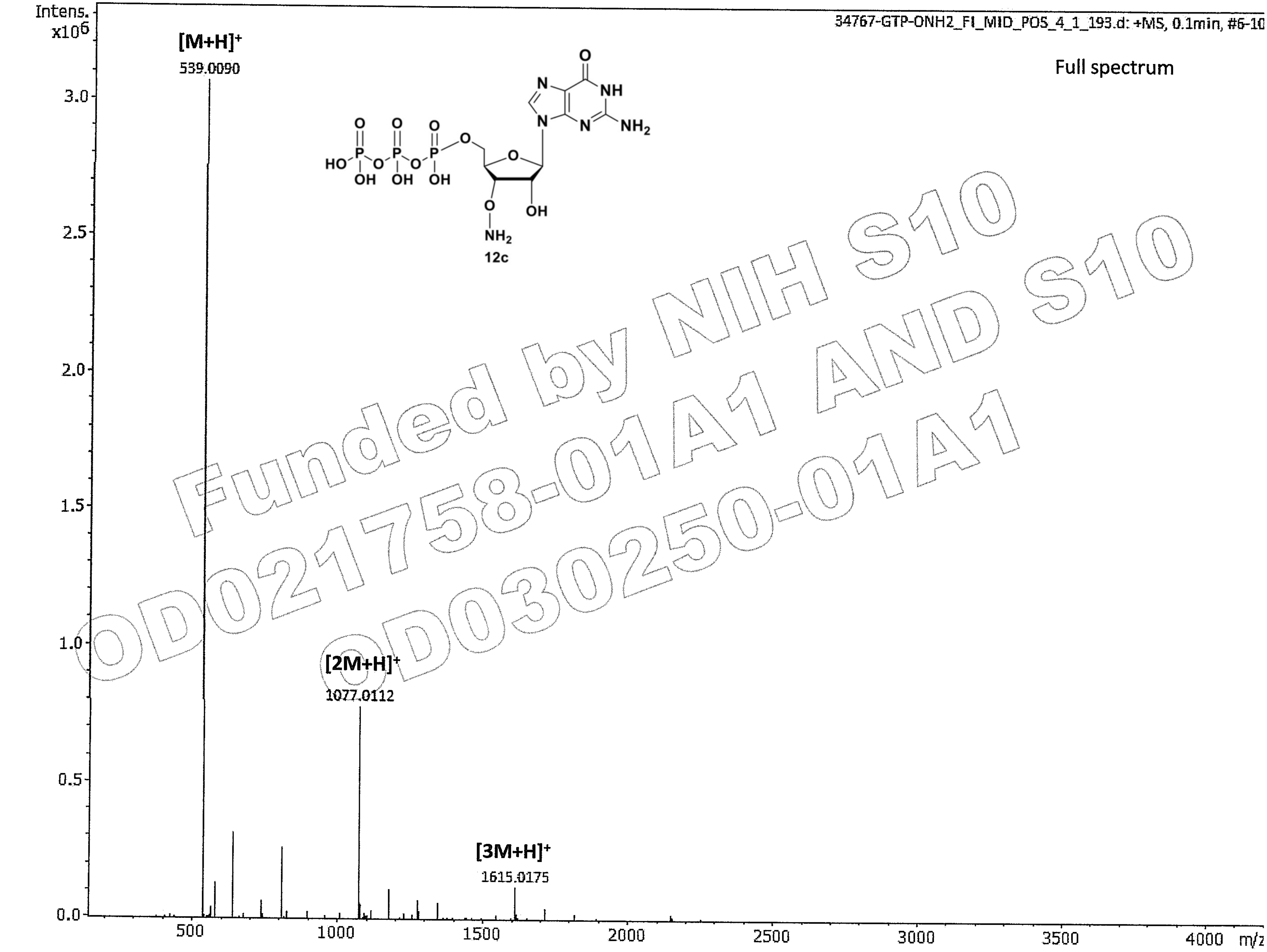

Funded by NIH S10 OD021758-01A1 AND S10 OD030250-01A1

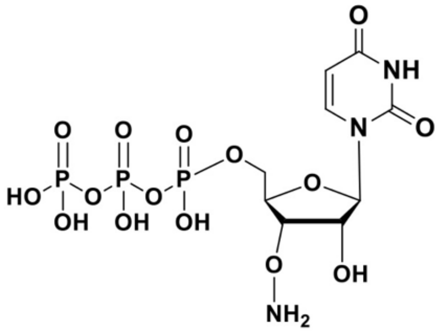

12d

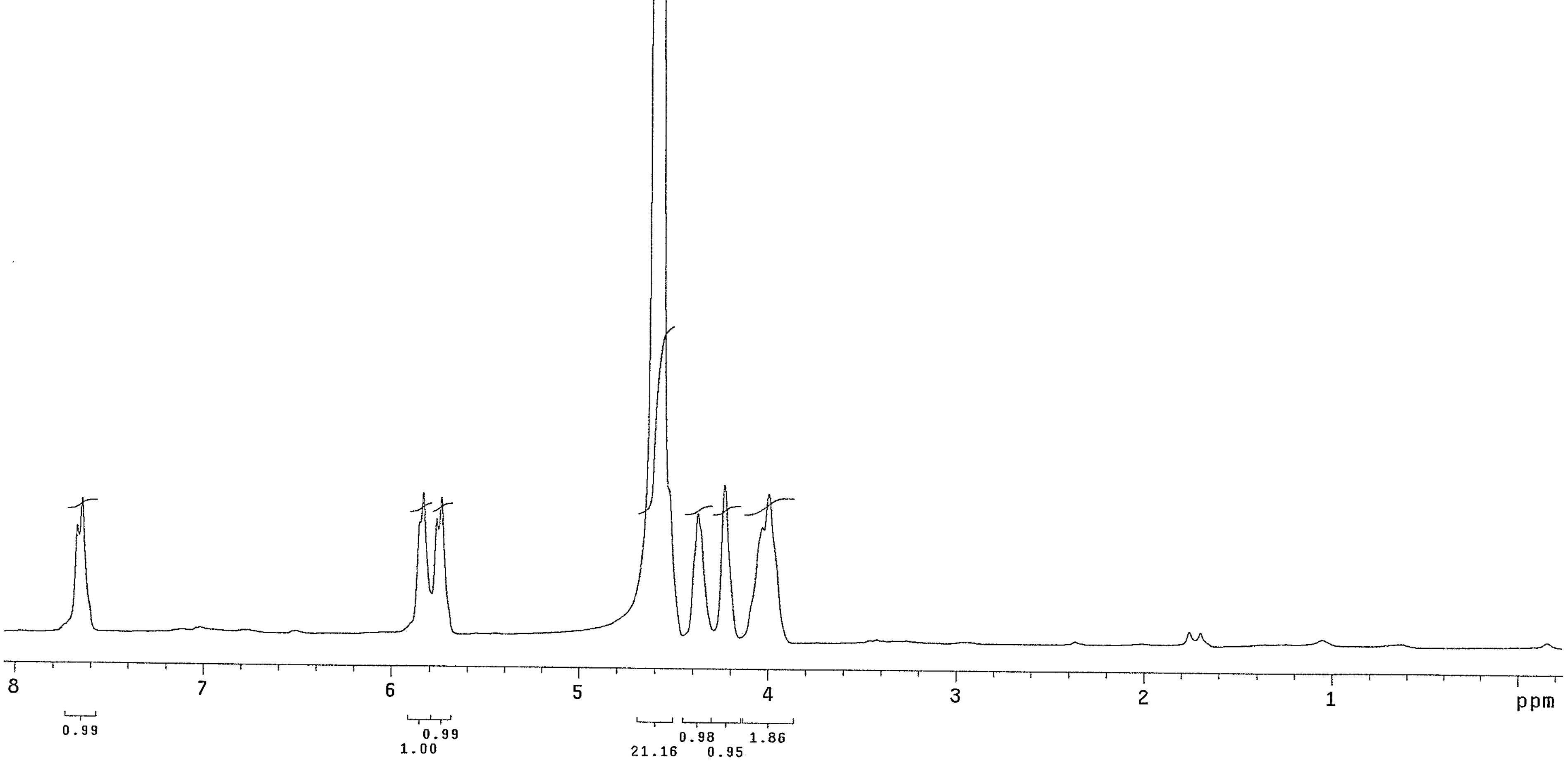

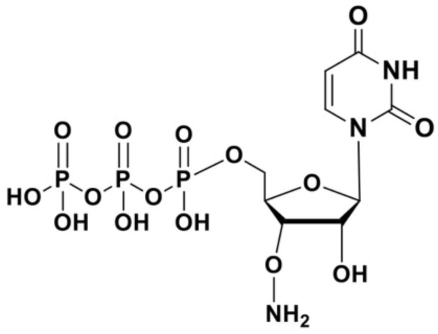
O
NH
O
N
O
O
O
O
HO
P
O
P
O
P
OH
OH
OH
O
O
OH
NH2

12d

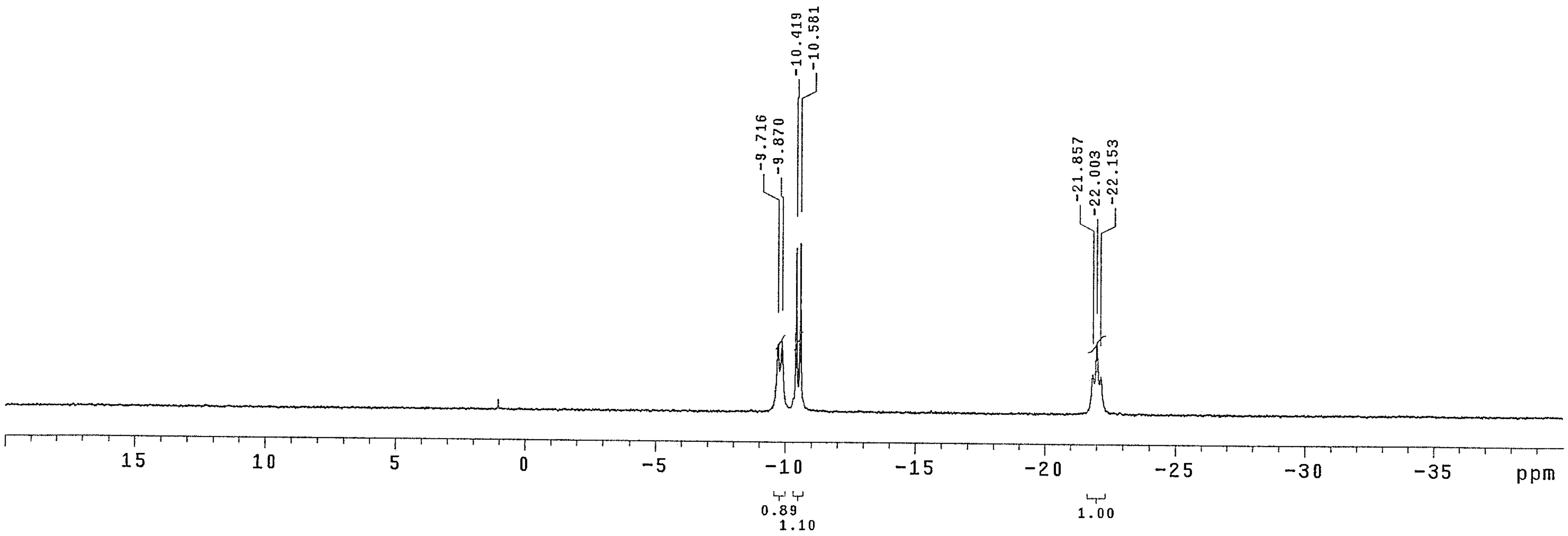
-9.716
-9.870
-10.419
-10.581
-21.857
-22.003
-22.153
15
10
5
0
-5
-10
-15
-20
-25
-30
-35
ppm
0.89
1.10
1.00

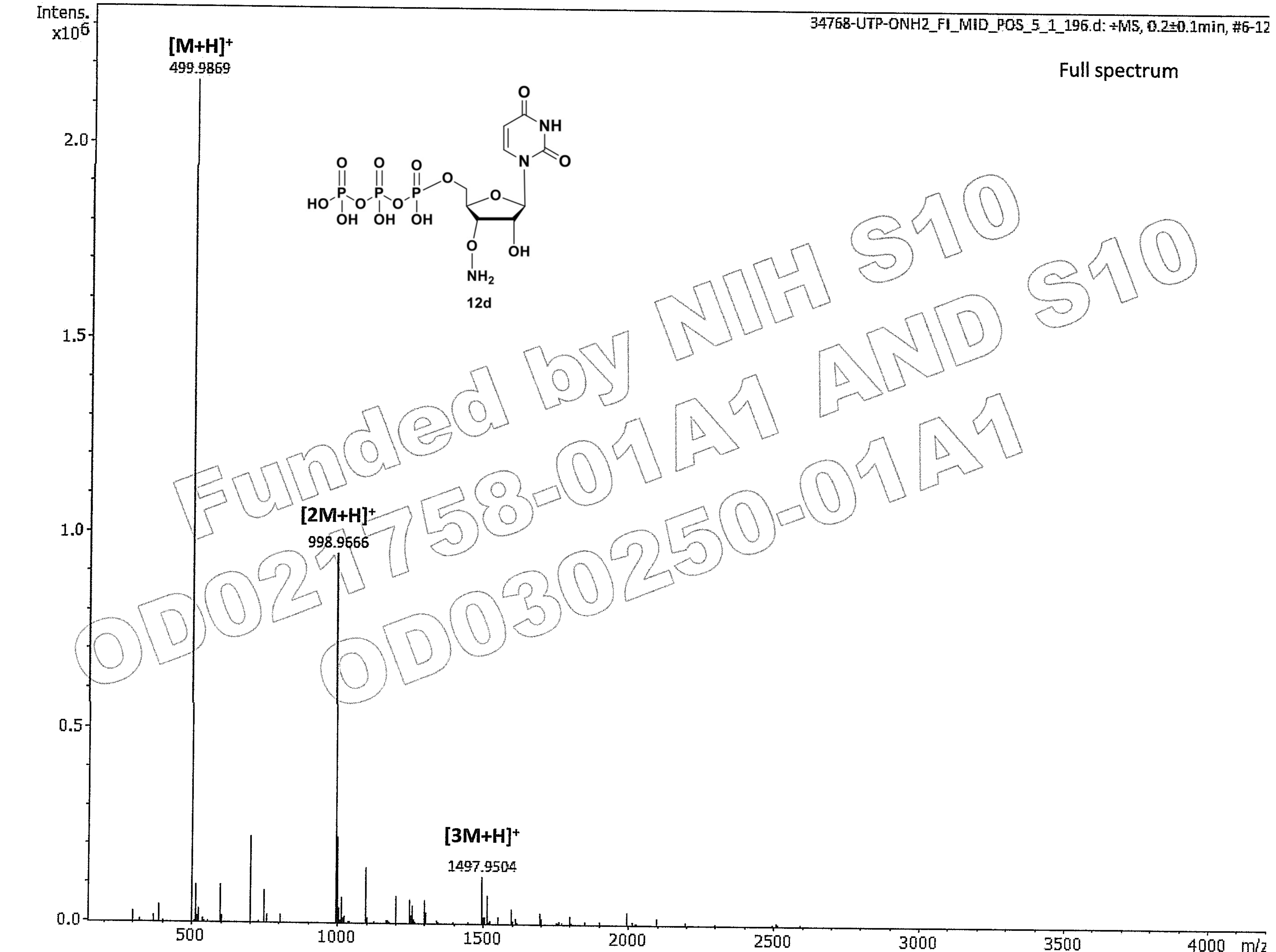

Funded by NIH S10 OD021758-01A1 AND S10 OD030250-01A1
